# Cysteine reactivity profiling links metabolic rewiring to the dysfunctional human T cell proteome

**DOI:** 10.64898/2026.03.04.709614

**Authors:** Hiroyuki Kojima, Charlotte R. Wayne, Luis F. Somarribas Patterson, Henry Sanford, Tzu-Jou Chen, Ya-Hui Lin, Joshua D. Schoenfeld, Lisa McGary, Yan-Ting Chen, Korbinian N. Kropp, Beatrice Zhang, Jahan Rahman, Tiffany L. Zhang, Nathalie Ropek, Cameron Roberts, Yuxi Ai, Kartikeya M. Menon, A. Ari Hakimi, Jiankun Lyu, Christopher A. Klebanoff, Omar Abdel-Wahab, Santosha A. Vardhana, Ekaterina V. Vinogradova

## Abstract

T cells exposed to persistent antigen in the context of chronic viral infections or tumors lose cytotoxic capacity. Post-transcriptional regulatory mechanisms contributing to human T cell dysfunction remain poorly understood. Complementary proteomic and cysteine reactivity profiling of human T cells undergoing chronic antigen-driven dysfunction identified post-translational changes associated with activation of multiple stress pathways, including the stress kinase MAP2K4, which were validated in primary TILs. Functional analysis revealed that many reactivity changes occurred within nucleotide-binding proteins, reflecting ATP loss due to chronic antigen-driven suppression of oxidative TCA cycle metabolism. Targeting MAP2K4 restored T cell cytotoxic capacity during prolonged tumor cell co-culture. These findings reveal a mechanism by which metabolic rewiring alters protein function and lay the groundwork for cysteine-directed therapeutics to enhance cancer immunotherapy.

## INTRODUCTION

Persistent T cell receptor (TCR) stimulation due to chronic antigen exposure is necessary and sufficient to impair the function of both CD4^+^ and CD8^+^ T cells during chronic viral infections as well as tumorigenesis^1, 2, 3, 4, 5, 6, 7, 8^. This dysfunctional state is associated with genome-wide alterations in both the T cell transcriptome^9^ and epigenome^10, 11, 12^. More recently, T cell dysfunction has been linked to proteotoxic stress^13, 14, 15^ and changes in T cell metabolism, including the accumulation of mitochondrial reactive oxygen species (ROS), which is sufficient to compromise mitochondrial ATP production and drive a terminally dysfunctional state in mouse T cells^4, 16, 17^. Loss of mitochondrial ATP production may disrupt ATP-dependent protein synthesis and folding, which are among the most bioenergetically demanding processes in eukaryotic cells^18^. Moreover, ROS can directly alter protein function by oxidizing proteinaceous thiols. Neither the functional status of the exhausted T cell proteome, nor whether it is impacted by T cell metabolic state, has been established.

In this study, we leveraged advanced cysteine-targeting chemical proteomic workflows – activity-based protein profiling with sample multiplexing using tandem mass tags (TMT-ABPP)^19, 20^ – to investigate the impact of chronic antigen stimulation on the functional proteomic landscape of dysfunctional human T cells. We recently deployed this approach to characterize reactivity changes following acute TCR stimulation, demonstrating that cysteine can serve as a “sensor” for structural and functional protein changes under different conditions, including shifts in cellular redox homeostasis^20^. We hypothesized that cysteine reactivity profiling could identify post-translationally regulated proteins critical to T cell dysfunction but overlooked by traditional transcriptomic and epigenetic analyses. Here, we demonstrate extensive remodeling of both the T cell proteome and cysteine reactivity landscape during chronic TCR stimulation, including widespread reactivity changes in nucleotide-binding proteins. Parallel metabolomic analyses demonstrated that suppressed oxidative metabolism and mitochondrial ATP synthesis, rather than mitochondrial ROS, are the primary drivers of these functional changes in human T cells. Cysteine reactivity profiling combined with ATP add-back provided mechanistic insights into how metabolic rewiring reshapes the functional proteome and linked these changes to the activation of stress-response pathways. We showed activation of key stress-response pathways, including the integrated stress response (ISR), endoplasmic reticulum (ER) stress signaling, and the stress-activated kinase MAP2K4. We further validated MAP2K4 activation across several orthogonal *in vivo* models of T cell exhaustion and showed that pharmacologic inhibition of MAP2K4 with covalent small-molecule electrophiles restores cytotoxic T cell function during prolonged antigen encounter. Together, these findings underscore the unique value of cysteine reactivity profiling in characterizing a previously inaccessible layer of post-translational regulation during T cell dysfunction and identifying novel targets for cancer immunotherapy.

## RESULTS

### Transcriptome-independent functional proteomic changes during chronic antigen-driven human T cell dysfunction

We first sought to develop an *in vitro* system that would allow us to assess the impact of chronic antigen exposure on the proteomic and cysteine reactivity landscape of primary human T cells, by comparing CD3^+^ T cells from the peripheral blood of healthy donors that had been activated with αCD3 and αCD28 antibodies for 2 days (“activated” state, D2) with activated T cells that had been passaged for an additional 2, 6, or 13 days in the presence of IL-7 and IL-15 to promote cytokine-driven expansion (“acute” state, D4A, D8A, and D15A) or in the presence of IL-7 and IL-15 along with continued αCD3-mediated TCR stimulation (“chronic” state, D4C, D8C, and D15C) (Fig. 1a). Flow cytometry analysis revealed that chronic stimulation of activated human T cells increased expression of multiple exhaustion markers, including inhibitory receptors (PD-1, LAG3, CTLA4, CD39) and transcription factors (TOX, T-Bet/TBX21) associated with human T cell dysfunction compared with acute T cells, which retained markers of activation (HLA-DR, CD38) but not exhaustion (Extended Data Fig. 1). Unbiased analysis of human T cell transcriptomes cultured under acute or chronic conditions using bulk RNA sequencing demonstrated progressive upregulation of genes uniquely associated with intratumoral T cell dysfunction across seven cancer types (“chronic activation score”)^2^ (Fig. 1b, Extended Data Fig. 2a; Data S1). Similarly, single-cell RNA sequencing of cells expanded in the absence or presence of chronic stimulation for 15 days revealed that while acute T cell transcriptomes mapped to a variety of cell states, nearly all chronically stimulated CD8^+^ T cells most closely mapped to exhausted CD8^+^ T cells from patient tumors (Fig. 1c,d)^6^. Moreover, chronic TCR stimulation reproduced functional hallmarks of T cell exhaustion, including reduced T cell proliferation (Extended Data Fig. 2b) and attenuated intracellular cytokine accumulation following restimulation (Extended Data Fig. 1). Finally, we adapted an established antigen-specific serial cytotoxicity assay^21^ in which human T cells transduced with a transgenic TCR targeting the NY-ESO-1 cancer testis antigen (LY) were acutely or chronically stimulated as described in Figure 1a and then co-cultured with NY-ESO-1 expressing cancer cell lines for an additional eight days, with replenishment of cancer cells every 2 days. Re-stimulated acute T cells (D8A) killed target cells in a TCR-specific, dose-dependent fashion, while chronically stimulated T cells (D8C) were markedly impaired in killing capacity (Fig. 1e,f).

**Fig. 1.**
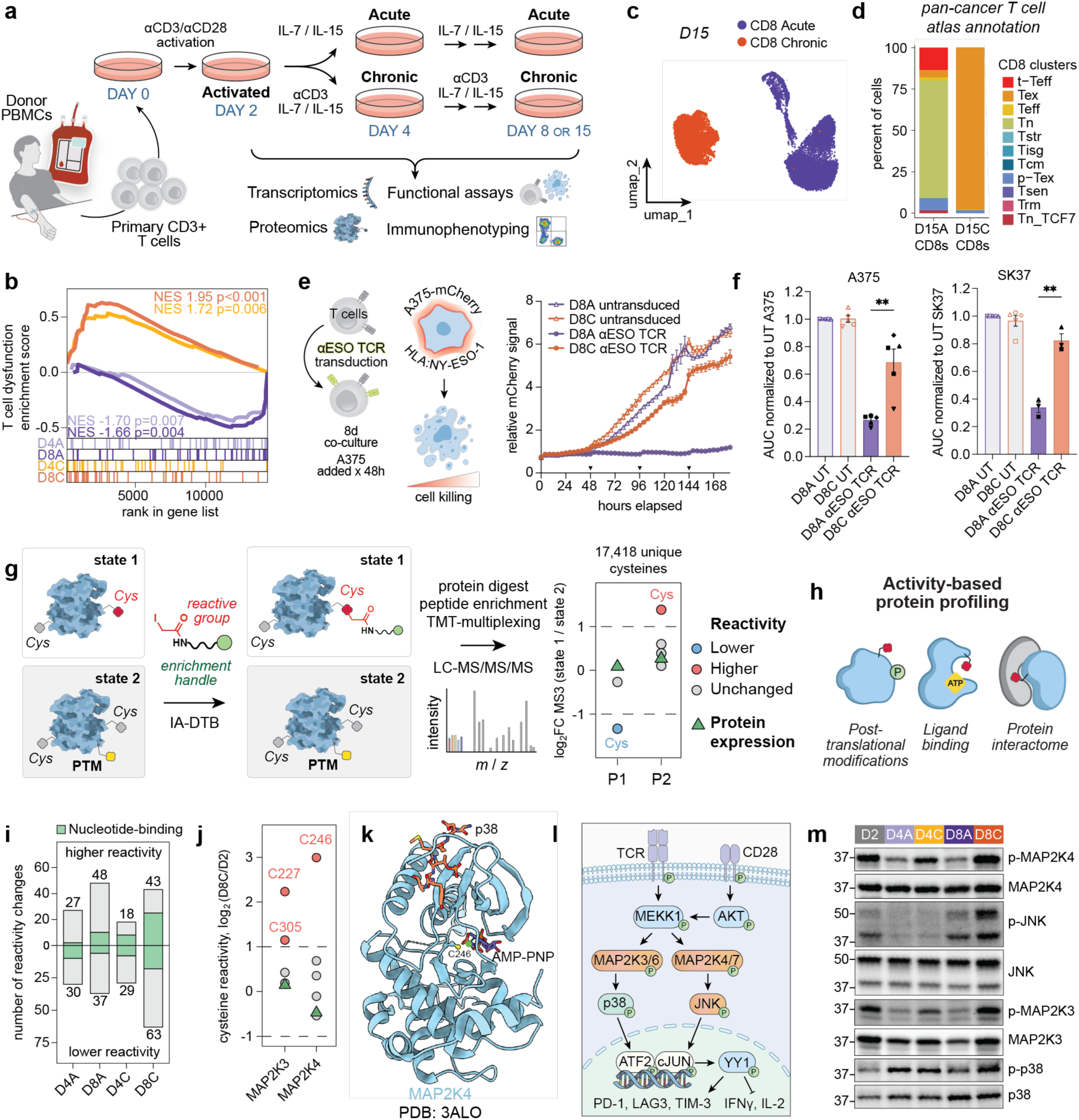
Reactivity profiling facilitates characterization of post-translational changes in T cell dysfunction. (**a**) Experimental setup for generation and multi-omic profiling of primary human T cells undergoing cytokine-mediated expansion with or without chronic TCR stimulation. (**b**) Gene Set Enrichment Analysis (GSEA) on a list of 43 genes in the “chronic activation program” associated with pan-cancer T cell dysfunction^2^. Each condition is compared to D2. NES – normalized enrichment score. (**c**) Uniform Manifold Approximation and Projection (UMAP) of single-cell transcriptomes from D15A and D15C CD8^+^ T cells. (**d**) Annotation of acute and chronic CD8^+^ T cell phenotypes (D15A and D15C) based on label transfer of single-cell transcriptomic data from Chu et al.^6^. (**e**) Antigen-specific cancer cell killing by D8A and D8C T cells transduced with NY-ESO-1-specific TCR. A representative cell viability curve of NY-ESO-1-expressing A375 melanoma cells engineered to express mCherry following 184 hours of co-culture with T cells, normalized to 0 h condition is shown. Inverted triangles on x axis indicate timepoints when A375 melanoma cells were added to the co-culture. (**f**) Area under the curve (AUC) of relative mCherry signal in A375 and SK37 melanoma cells over 48-184 hours for n = 3-5 independent donors, normalized to untransduced (UT) D8A. Statistical comparison by two-tailed paired t test (**, p < 0.01). Donor-matched conditions are indicated by differently shaped data points. (**g**) Schematic of reactivity profiling experiments using a cysteine-reactive chemical probe, iodoacetamide-desthiobiotin (IA-DTB) (Data S2-3, S2-4). PTM – post-translational modification. See Materials and Methods for more details. (**h**) Schematic showing biochemical drivers of reactivity changes in TMT-ABPP. (**i**) Bar graph of reactivity changes across cysteine reactivity profiling in n = 5 donors. Nucleotide-binding proteins (based on Gene Ontology annotation) are shown in green. Higher/lower reactivity was determined by comparison to whole proteome or median cysteine reactivity. (**j**) Cysteine reactivity log_2_ fold change values between D8C and D2 T cells for MAP2K3 and MAP2K4. Protein expression log_2_ fold change values are shown as green triangles. Cysteine reactivity data is from n = 5 donors and protein expression data is from n = 6 donors. (**k**) Crystal structure of autoinhibited non-phosphorylated MAP2K4 in complex with AMP-PNP and p38 peptide (PDB: 3ALO^22^) with differentially reactive cysteine C246 labeled near AMP-binding site. (**l**) Schematic of MAPK pathways driven by chronic αCD3/αCD28 stimulation. Proteins with more phosphorylation in D8C than in D2 T cells are shown in orange. (**m**) Western blot analysis of phosphorylation levels of MAP2K4 (Ser257), JNK (Thr183/Tyr185), MAP2K3 (Ser189), and p38 (Thr180/Tyr182) relative to protein expression.

We then performed complementary mass spectrometry-based proteomics experiments to examine protein-level changes and their correlation with gene expression assessed by RNA-sequencing during persistent TCR stimulation (Extended Data Fig. 2c; Data S2-1 and S2-2). Using isobaric tandem mass tag (TMT) multiplexing and normalization by protein content enabled us to directly compare protein abundances in activated, acutely, and chronically stimulated T cells at multiple timepoints. PCA revealed clear separation among activated, acute, and chronic T cell populations that increased over time (Extended Data Fig. 2d). Importantly, key exhaustion markers were consistently upregulated at both transcriptomic and proteomic levels in chronically stimulated cells (Extended Data Fig. 2e). However, a substantial number of proteins exhibited changes only at the proteomic level, including proteins involved in mitochondrial ROS detoxification (GPX1, GPX4), peroxisomal function (PXMP2), and cell-cycle progression (CDKN2A/p16^23^), suggesting the existence of post-transcriptional regulatory mechanisms in T cell dysfunction (Extended Data Fig. 2e,f)^24, 25^.

We recently developed a chemical proteomic platform termed TMT-ABPP, which uses broadly reactive iodoacetamide probes to globally assess cell state-dependent changes in cysteine reactivity (Fig. 1g)^19, 20^. This method is enabled by sample multiplexing using tandem mass tags (TMT) and offers a new approach to characterize structural and functional proteomic landscapes that have been overlooked by traditional gene and protein expression analyses across different T cell states (Fig. 1h). Cysteine is a highly nucleophilic amino acid that exhibits reactivity towards electrophilic chemical probes. Cysteine reactivity can be influenced by multiple factors, including direct and indirect post-translational modifications and protein interaction networks with proteins, nucleic acids, and metabolites (Fig. 1h)^20, 26^. We mapped cysteine reactivity changes across activated (D2), acute (D4A and D8A), and chronic (D4C and D8C) T cells by treating cell lysates with a broad-spectrum cysteine-reactive iodoacetamide-desthiobiotin (IA-DTB) probe, followed by trypsin and Lys-C digestion, streptavidin-mediated enrichment of desthiobiotinylated cysteine-containing peptides, TMT-based multiplexing, and analysis via LC/LC-MS/MS/MS (Fig. 1g). We identified reactivity changes using a cutoff of >2-fold signal intensity differences normalized to activated cells in proteins with at least two quantified cysteine-containing peptides or with a single quantified peptide and matching unenriched proteomics data (Data S2-3, S2-4).

Reactivity profiling revealed more than 150 reactivity changes across different cell states, with the largest number observed in chronically stimulated D8C cells (Fig. 1i). Global analysis of identified reactivity changes revealed an enrichment in nucleotide-binding proteins under chronically stimulated conditions (Fig. 1i). Among these, we observed state-dependent reactivity differences in cysteines within nucleotide-binding domains of stress-associated kinases MAP2K4 and MAP2K3 (Fig. 1j,k and Extended Data Fig. 2g)^22^, which are upstream of JNK and p38 stress-induced pathways (Fig. 1l). This suggests that the increased reactivity of MAP2K4_C246 and MAP2K3_C227 in D8C T cells may result from reduced nucleotide occupancy of the kinase active site and/or allosteric effects of activation loop phosphorylation. Consistent with this, MAP2K4_C246 was previously shown to exhibit increased reactivity following gel filtration of mitotic HeLa cell lysates, potentially reflecting the removal of bound nucleotides; its reactivity was also associated with the phosphorylation and activation state of MAP2K4^26^. We confirmed activation of MAP2K3 and MAP2K4-JNK pathways in chronically stimulated (D8C), but not acute (D8A), T cells by Western blot (Fig. 1m and Extended Data Fig. 2h). Indeed, chronic T cell stimulation-driven p38 activation has been shown to increase T cell dysfunction^27^, whereas inhibition of p38α can increase T cell expansion, expression of stemness markers, and antitumor efficacy^28^. These findings highlight the potential of TMT-ABPP to identify therapeutically relevant functional protein changes not identified using classical approaches such as gene or protein expression analyses.

### Impaired nucleotide synthesis alters the functional proteome of exhausted T cells

We had previously shown that chronic stimulation of mouse T cells causes mitochondrial dysfunction and impairs nucleotide triphosphate synthesis^4^, leading us to hypothesize that the reactivity changes observed during chronic TCR stimulation of human T cells were a result of compromised nucleotide pools. PCA based on steady-state abundances of the 156 targeted polar metabolites quantified across activated, acute, and chronic T cells by QQQ-LC/MS showed that T cells could be reproducibly separated based on polar metabolite abundance alone (Fig. 2a). Interestingly, nucleotides were the strongest drivers of separation between D2, D8A, and D8C cells, with nucleotide di- and triphosphates accumulating in D2 cells and intermediates in nucleotide salvage accumulating in D8C cells (Fig. 2a). Analysis of differentially abundant metabolites showed that compared with D2 cells, D8C cells exhibited a substantial decrease in mitochondrial TCA cycle metabolites (malate, fumarate, oxoglutarate, succinate), as well as mitochondrial precursors of nucleotide biosynthesis (dihydroorotate, aspartate), accumulating upstream glycolytic intermediates (glucose, glucose-6P, fructose-6P) and metabolites reflective of nucleotide salvage (adenosine, xanthosine, uridine, guanosine) (Fig. 2b,c and Extended Data Fig. 3a; Data S3-1). As both *de novo* synthesis of purines and pyrimidines depend on mitochondrial ATP as well as oxidative synthesis of aspartate^29, 30^, activation of nucleotide salvage is consistent with reduced mitochondrial electron transport in chronically stimulated cells^31^. Assessment of the relative phosphorylation state of nucleotides revealed that while the ratio of triphosphorylated to diphosphorylated species did not substantially differ between D2, D8A, and D8C cells, the ratio of nucleosides to monophosphorylated species was substantially elevated in D8C cells across 3 nucleotide species (Data S3-2). These results are consistent with catabolism of the adenylate pool in order to stabilize the energy charge under conditions of energy depletion, which has been described in skeletal muscle as well as primary human lymphocytes^32, 33^, and suggest that the observed reactivity changes in D8C cells are a result of a collapse of the nucleotide pool, rather than the energy charge of the residual adenylate pool, in the setting of increased bioenergetic demand.

**Fig. 2.**
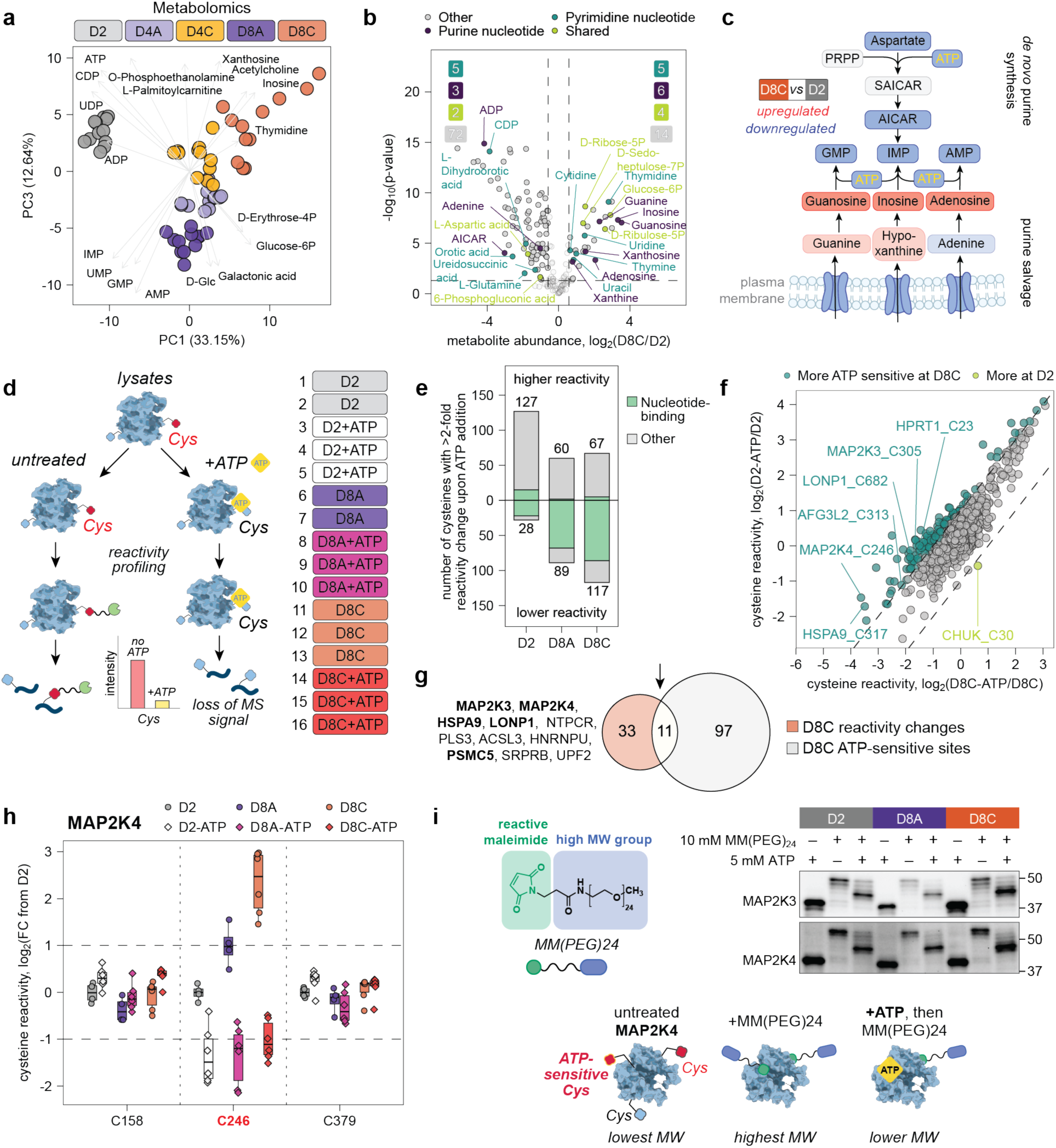
Cysteine reactivity chemical proteomics and metabolic profiling reveal dysfunction of *de novo* purine synthesis in chronically stimulated T cells. (**a**) PCA of metabolomics data from n = 4 donors. Signal intensity values were log_2_ transformed and only metabolites quantified in all replicates were used. The top and bottom five loadings from PC1 and PC3 are shown with grey arrows. (**b**) Volcano plot showing log_2_ fold changes of metabolite abundances between D8C and D2 T cells. Metabolites are colored based on manually curated pathway lists retrieved from KEGG. Dashed lines represent cutoffs of p-value < 0.05 and fold-change > 1.5. Data are from n = 4 donors. (**c**) Schematic of purine *de novo* synthesis and salvage pathways. Metabolites are colored according to fold change between D8C and D2 T cells. (**d**) Schematic of ATP add-back experiment and diagram of TMT 16-plex channels (Data S2-5). See Materials and Methods for more details. (**e**) Quantification of cysteines with two-fold difference in cysteine reactivity between control and ATP add-back conditions in D2, D8A, and D8C T cells. Nucleotide-binding proteins (green) are labeled based on Gene Ontology annotation. (**f**) Scatter plot of log_2_ fold changes between control and ATP add-back conditions in D8C (x axis) and D2 (y axis) T cells. Dashed lines indicate a cutoff for ratio of fold change between D8C and D2 T cells of > 2. (**g**) Venn diagram of reactivity changes and ATP-sensitive sites in D8C cells. Only cysteines quantified in both datasets are shown. (**h**) Boxplot of log_2_ fold changes in cysteine reactivity of MAP2K4 in control (circle) and ATP add-back (diamond) conditions in D2, D8A, and D8C T cells compared to D2 control condition. Boxes mark the lower and upper quartiles, horizontal black lines mark the median value, and whiskers extend to the furthest point within 1.5x the interquartile range. (**i**) Validation of ATP-sensitivity of reactive cysteines in MAP2K3 and MAP2K4 using activity-based protein profiling with a high-molecular weight cysteine-reactive maleimide probe and Western blot visualization (WB-ABPP).

Next, to directly test whether the observed insufficiency in mitochondrial ATP synthesis was directly responsible for the observed state-dependent differences in cysteine reactivity, we conducted a “function-first” ATP add-back experiment in which T cell lysates from activated (D2), acute (D8A), and chronic (D8C) cells were treated with 5 mM ATP for 10 min before exposure to a broadly reactive iodoacetamide probe, followed by the standard TMT-ABPP workflow (Fig. 2d; Data S2-5). We anticipated that cysteines directly or allosterically affected by ATP binding would exhibit changes in reactivity. As expected, most decreased reactivity changes were observed in nucleotide-binding proteins (Fig. 2e). Reactive cysteines in chronically stimulated T cells were more sensitive to ATP add-back than in activated or acute conditions (Fig. 2e,f). Several proteins with reactivity changes were sensitive to ATP add-back, including MAP2K4_C246 and MAP2K3_C305, both of which showed decreased cysteine reactivity following ATP add-back (Fig. 2g,h). We further validated T cell state-dependent ATP sensitivity of MAP2K3 and MAP2K4 cysteine reactivity by performing an in-gel ABPP experiment with a high-molecular weight maleimide-based cysteine-reactive probe, followed by Western blot visualization of MAP2K3 and MAP2K4, both of which showed probe- and ATP-dependent band shifts (Fig. 2i). Notably, higher ATP concentrations were required to prevent probe-dependent MAP2K3 and MAP2K4 band shifts in D8C than in D2 T cells (apparent EC_50_ for MAP2K4: 0.077 mM in D2 vs 0.300 mM in D8C; apparent EC_50_ for MAP2K3: 0.134 mM in D2 vs 0.328 mM in D8C), consistent with reduced availability of ATP in chronic T cells (Extended Data Fig. 2i,j).

Beyond stress kinase activation, five out of six ATPase subunits of the 26S proteasome complex (PSMC proteins) exhibited ATP-sensitive reactivity changes, whereas non-ATPase subunits (PSMA, PSMB, and PSMD proteins) remained unaffected (Extended Data Fig. 3b). We confirmed the ATP dependence of these reactivity changes using an existing activity-based probe targeting proteasome catalytic subunits (Extended Data Fig. 3c)^34^. Proteasome activity was higher in chronically stimulated cells, in line with previous literature showing non-linear relationship between ATP concentrations and proteasome activity^35^, and decreased across all conditions following ATP add-back (Extended Data Fig. 3d). These results are consistent with prior literature implicating proteasome activation in T cell exhaustion^15^, while suggesting that nucleotide-dependent regulation of proteasome activity is mechanistically distinct from changes in the abundance of individual proteasome subunits.

We also identified ATP-sensitive cysteine reactivity changes in two members of the mitochondrial protein import complex, HSPA9 and LONP1, both of which play key roles in mitochondrial protein quality control (Extended Data Fig. 3e-i)^36, 37, 38, 39, 40, 41^. To determine whether the observed cysteine reactivity changes in LONP1 were associated with functional alterations, we separated cells into detergent-soluble and insoluble fractions (Extended Data Fig. 3j,k). This method enables evaluation of improperly folded or aggregated proteins, which accumulate in the insoluble pellet, such as the HSPA9 co-chaperone DNAJA3^40^. DNAJA3, but not other LONP1-sensitive proteins HSPA9 and NDUFA9, accumulated in the detergent-insoluble fraction following chronic stimulation, consistent with partial impairment of LONP1 ATPase function (Extended Data Fig. 3k-m).

To more broadly assess protein aggregation^13, 14^ in human T cells and determine their contribution to the observed reactivity changes, we performed quantitative proteomics of detergent-soluble and insoluble fractions (Extended Data Fig. 4a and Data S2-6). Few proteins showed T cell state-dependent accumulation in the insoluble fraction (Extended Data Fig. 4b,c). Proteins harboring differentially reactive cysteines, including MAP2K3 and MAP2K4, were more enriched in detergent-soluble fraction, suggesting that the global cysteine reactivity changes were primarily driven by biochemical and structural changes associated with biological cues but not protein misfolding (Extended Data Fig. 4b-e). However, consistent with our Western blot analysis, DNAJA3 and LONP1 were substantially enriched in the detergent-insoluble fraction, suggesting partial disruption of protein quality control system in human T cells (Extended Data Fig. 4f).

### Metabolic rewiring to limit ROS accumulation leads to integrated, ER, and proteotoxic stress responses during chronic TCR stimulation

Chronically stimulated T cells showed cysteine reactivity changes in protein classes related to protein homeostasis and stress responses. Notably, we also observed ATP-sensitive reactivity changes in chronically stimulated T cells in three cysteine residues within the scaffold protein GCN1, which binds and promotes the activation of GCN2 and ISR in response to tRNA uncharging as well as ribosomal collisions (Fig. 3a and Extended Data Fig. 5a)^42^. Of the ATP-sensitive reactive cysteines, two (C55, C648) are located in the polysome-binding region, while the third (C2179) resides within the GCN2-binding domain^42^, which is known to be ATP-regulated. We therefore hypothesized that chronic antigen-driven impairment in mitochondrial ATP production could lead to activation of multiple stress response pathways due to impaired ATP-dependent protein folding and GCN1 activation (Fig. 3b)^43, 44, 45^. Western blot analysis showed that chronic TCR stimulation induced both the ISR (p-eIF2α, ATF4, CHOP)^46^ and the ER stress response (ATF6, p-PERK, BiP, and the spliced form of XBP1) (Fig. 3c,d and Extended Data Fig. 5b-d), which we confirmed by gene set enrichment analysis of genes containing ATF4-or ATF6-binding motifs (Fig. 3e,f) and upregulation of ISR-regulated genes (Extended Data Fig. 5e).

**Fig. 3.**
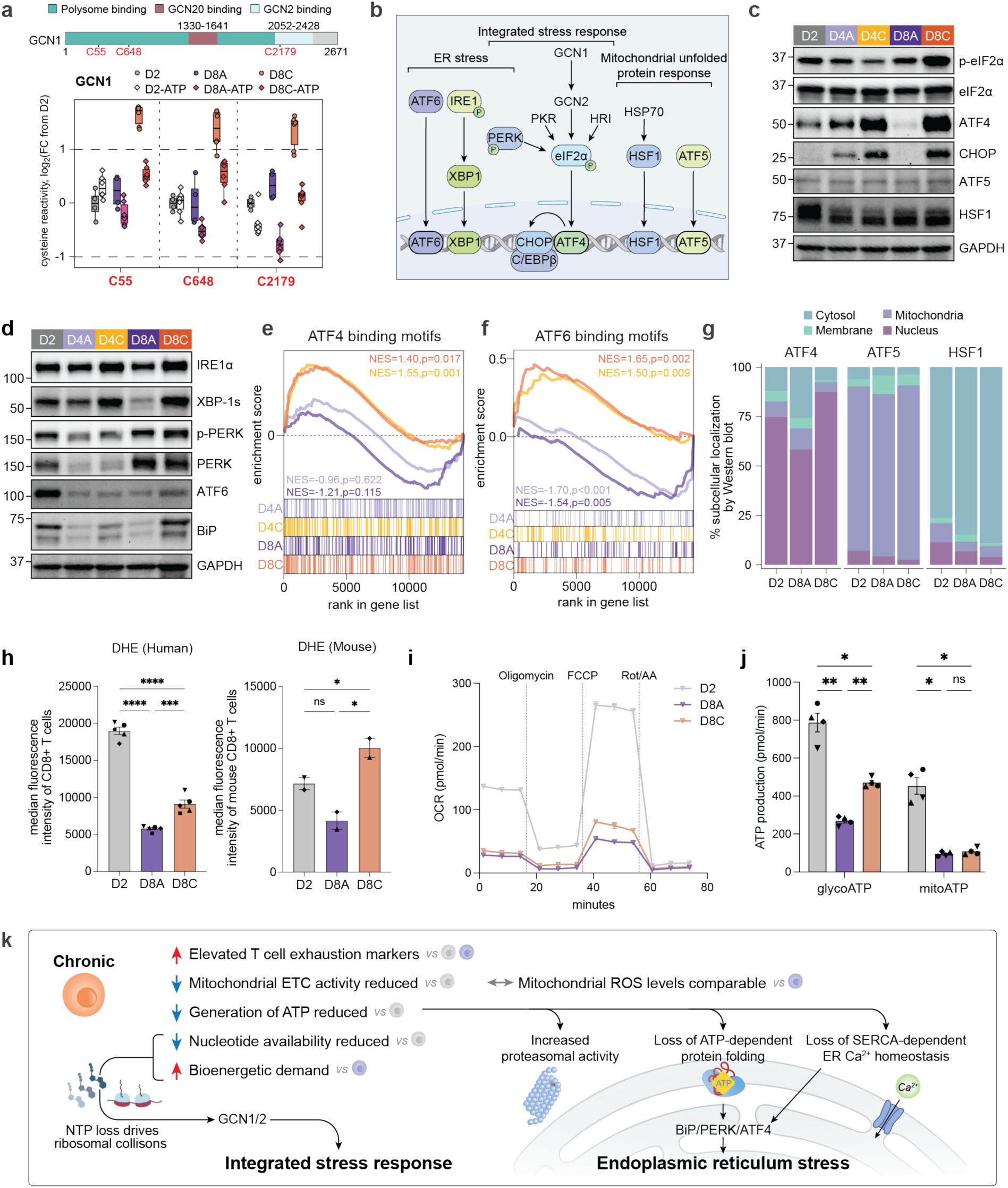
Differential activation of integrated, mitochondrial, and ER stress responses during chronic TCR stimulation. (**a**) Boxplot of log_2_ fold changes in cysteine reactivity of GCN1 in control (circle) and ATP add-back (diamond) conditions in D2, D8A, and D8C T cells compared to D2 control condition. Boxes mark the lower and upper quartiles, horizontal black lines mark the median value, and whiskers extend to the furthest point within 1.5x the interquartile range. Data are from n = 2 independent cysteine reactivity profiling experiments. Sequence of GCN1 with annotated polysome-, GCN20-, and GCN2-binding domains is shown^42, 50^. (**b**) Schematic of mtUPR, ISR, and ER stress-response pathways. (**c**) Western blot analysis of protein expression in ISR (ATF4, CHOP, p-eIF2α) and mtUPR (ATF5, HSF1) pathways. GAPDH included as loading control for ATF4 membrane only. (**d**) Western blot analysis of protein expression of ER stress signaling pathway members. GAPDH included as loading control for IRE1α membrane only. (**e**) GSEA of bulk RNA-sequencing data on the Hallmark gene set for ATF4_BINDING_MOTIFS. Each condition was compared to D2 T cells. Gene expression data are from n = 3 donors. (**f**) GSEA of bulk RNA-sequencing data on Hallmark gene sets for ATF6_BINDING_MOTIFS. Each condition was compared to D2 T cells. Gene expression data are from n = 3 donors. (**g**) Western blot quantification of subcellular localization of ATF4, ATF5, and HSF1. Data are shown as average of n = 2 donors. (**h**) Quantification of oxidative stress in human and mouse CD8^+^ T cells by flow cytometry using dihydroethidium (DHE; pan-cellular superoxides) ROS-reactive dye ^4^. For human samples, statistical comparison by repeated measures one-way ANOVA with Šídák’s multiple comparisons test (***, p < 0.001; ****, p < 0.0001). Donor-matched conditions are indicated by differently shaped data points. Data are from n = 5 donors. For mouse samples, statistical comparison by repeated measures mixed-effects analysis with Šídák’s multiple comparisons test (ns, p > 0.05; *, p < 0.05). Subject-matched conditions are indicated by differently shaped data points. Data are from n = 2 mice. (**i**) Representative plot of oxygen consumption rate (OCR) as measured by extracellular flux analysis in D2, D8A, and D8C cells. Injections of oligomycin (ATP synthase inhibitor), FCCP (mitochondrial uncoupler), and rotenone/antimycin A (electron transport chain inhibitors) are indicated by vertical lines. (**j**) Calculated mitochondrial and glycolytic ATP production based on extracellular flux analysis of D2 (gray), D8A (purple), and D8C (orange) human T cells. Statistical comparison by one-way repeated measures ANOVA with Tukey’s multiple comparisons test (ns, p > 0.05; *, p < 0.05; **, p < 0.01). Donor-matched conditions are indicated by differently shaped data points; data are presented as mean ± SEM. Data are from n = 4 donors. (**k**) Schematic showing molecular differences between activated, acutely, and chronically stimulated primary human T cells.

In contrast, ATF5^47^ and HSF1^48^, both of which are hallmarks of mitochondrial unfolded protein response (mtUPR) activation, were not induced or translocated into the nucleus following chronic TCR stimulation (Fig. 3g and Extended Data Fig. 5f). The mtUPR is known to be activated by mitochondrial ROS^49^, which has been established as a key driver of terminal T cell exhaustion in mouse models^4, 16, 17^. However, the ratio of oxidized to reduced glutathione (GSSG:GSH) did not significantly differ between D2, D8A, and D8C cells (Data S3-2). Moreover, while chronic TCR stimulation increased accumulation of total cellular as well as mitochondrial ROS in mouse T cells, chronically stimulated human T cells exhibited only a mild increase in cellular ROS and no increase in mitochondrial ROS (Fig. 3h and Extended Data Fig. 6a-c).

Finally, supplementation of free radical scavengers such as β-mercaptoethanol and *N-*acetylcysteine, which have been shown to improve the proliferation of mouse T cells during chronic TCR stimulation^4^, had no effect on human T cells, suggesting that chronic stimulation does not result in sufficient ROS accumulation to limit human T cell growth (Extended Data Fig. 6d). As mitochondrial ROS is generated during electron transport, the lack of mitochondrial ROS increase during chronic TCR stimulation suggested that oxidative phosphorylation might be actively suppressed in chronically stimulated human T cells, which we confirmed using extracellular flux analysis (Fig. 3i and Extended Data Fig. 6e,f). As a result, chronically stimulated T cells showed substantially less mitochondrial ATP production compared with activated T cells and similar to acute T cells despite the increased bioenergetic demand of chronic stimulation (Fig. 3j).

Chronic stimulation of human T cells did not decrease nutrient uptake overall, as extracellular acidification rates were higher in chronic T cells (Extended Data Fig. 6f). Rather, proteomic data showed that chronically stimulated cells decreased expression of components of complexes I-IV of the electron transport chain (ETC) (Extended Data Fig. 6g), suggesting that chronic T cells actively limit oxidative metabolism of carbon within the TCA cycle. To test this hypothesis, we performed stable isotope tracing of D2, D8A, and D8C cells cultured in the presence of universally ^13^C-labeled glucose or glutamine for 4 hours (Extended Data Fig. 6h,i; Data S3-3, S3-4). Analysis of glucose tracing showed that steady-state labeling of glycolytic intermediates by glucose was unimpaired in D8C cells, indicating that glucose uptake and glycolytic metabolism was unimpeded (Extended Data Fig. 6j). Steady-state labeling of glutamine as well as proximal glutamine metabolites glutamate and alpha-ketoglutarate (α-KG) was increased in D8C cells, indicating that both uptake and initial metabolism of glucose and glutamine was unimpaired by chronic TCR stimulation (Extended Data Fig. 6k). However, we observed a substantial decrease in downstream oxidative metabolism of either glucose or glutamine within the TCA cycle, as demonstrated by a significant decrease in m+2 labeling of fumarate, malate, and aspartate by ^13^C-glucose and a significant decrease in m+4 labeling of aspartate by ^13^C-glutamine. The mitochondrial TCA cycle consists of both oxidative (NAD^+^-dependent) and non-oxidative enzymatic reactions; the decrease in TCA cycle labeling by both ^13^C-glucose and ^13^C-glutamine was most potent for NAD^+^-dependent reactions catalyzed by IDH3, OGDH, and MDH2 (Extended Data Fig. 6h,i). We also observed a substantial increase in non-oxidative metabolism of glutamine in D8C cells to generate proline, which has been shown to limit mitochondrial reductive stress by diverting electrons for mitochondrial NADPH synthesis (Extended Data Fig. 6k)^51, 52, 53^. In addition to proline, lipid species, including triglycerides and ether phospholipids, act as electron acceptors and play established roles in antioxidant defense^54, 55^. Consistent with active diversion of mitochondrial NADH from the ETC, lipidomic profiling revealed chronically stimulated T cells accumulated both triglycerides and ether phospholipids compared with D2 cells (Extended Data Fig. 6l, Data S3-5). These findings are consistent with our proteomic data showing an increase in accumulation of proteins from peroxisomes, which are required for ether phospholipid synthesis, and indicate that chronically stimulated T cells limit donation of electrons to the mitochondrial ETC by shunting NADH towards alternative electron sinks, including proline and lipid synthesis (Extended Data Fig. 6m). In combination with activation of the ISR and ER stress response pathways, but not mtUPR, these results are consistent with decreased ATP availability rather than mitochondrial ROS being a fundamental driver of early dysfunction in chronically stimulated human T cells (Fig. 3k)^44, 56^.

### Stress kinase activation in *in vivo* exhausted T cells and primary human TILs

The dysfunctional T cell transcriptomic and epigenomic program, often referred to as T cell ‘exhaustion’, has classically been established in T cells from mice with chronic viral infections or from either mouse or human tumors. The proteomic landscapes of mouse or human T cells undergoing chronic antigen-driven dysfunction *in vivo* have not been well described. We therefore isolated CD8^+^ T cells from the spleens of mice bearing either acute (LCMV-Armstrong) or chronic (LCMV-Clone 13) viral infections as well as from the tumors and spleens of KPC tumor-bearing mice, which we directly compared with D2, D8A, or D8C mouse T cells *in vitro* using low-input unenriched proteomics (Fig. 4a and Extended Data Fig. 7a; Data S2-7). In parallel, we compared human CD3^+^PD-1^+^ TILs from primary renal cell carcinoma tumors together with D2, D8A, or D8C human T cells *in vitro* (Fig. 4b and Extended Data Fig. 7b; Data S2-8). PCA of mouse T cells revealed a first principal component (PC1) which predominantly separated *in vitro* from *in vivo* conditions and a second component (PC2) which predominantly separated T cells experiencing chronic antigen stimulation *in vitro* (D8C) or antigen stimulation *in vivo* (LCMV-Arm, LCMV-Clone 13, TIL), suggesting both antigen and environmentally driven proteomic alterations *in vivo* (Fig. 4c and Extended Data Fig. 7c). PCA of human T cells similarly separated human T cells cultured *in vitro* from primary TILs along PC1, with a smaller contribution of PC2 separating TILs and D8C cells from day 0 (D0), D2, and D8A cells (Fig. 4d). These results suggest that certain proteomic changes seen in dysfunctional T cells *in vivo* may not be fully recapitulated *in vitro*. However, key exhaustion markers were similarly regulated in both human and mouse across *in vitro* and *in vivo* models (Fig. 4e and Extended Data Fig. 7d,e). Importantly, we observed increased phosphorylation of MAP2K4 and JNK in dysfunctional T cells from LCMV-Clone 13 infected mice, KPC tumors, and primary human TILs from renal cell carcinoma tumors, recapitulating our findings from the *in vitro* system (Fig. 4f,g).

**Fig. 4.**
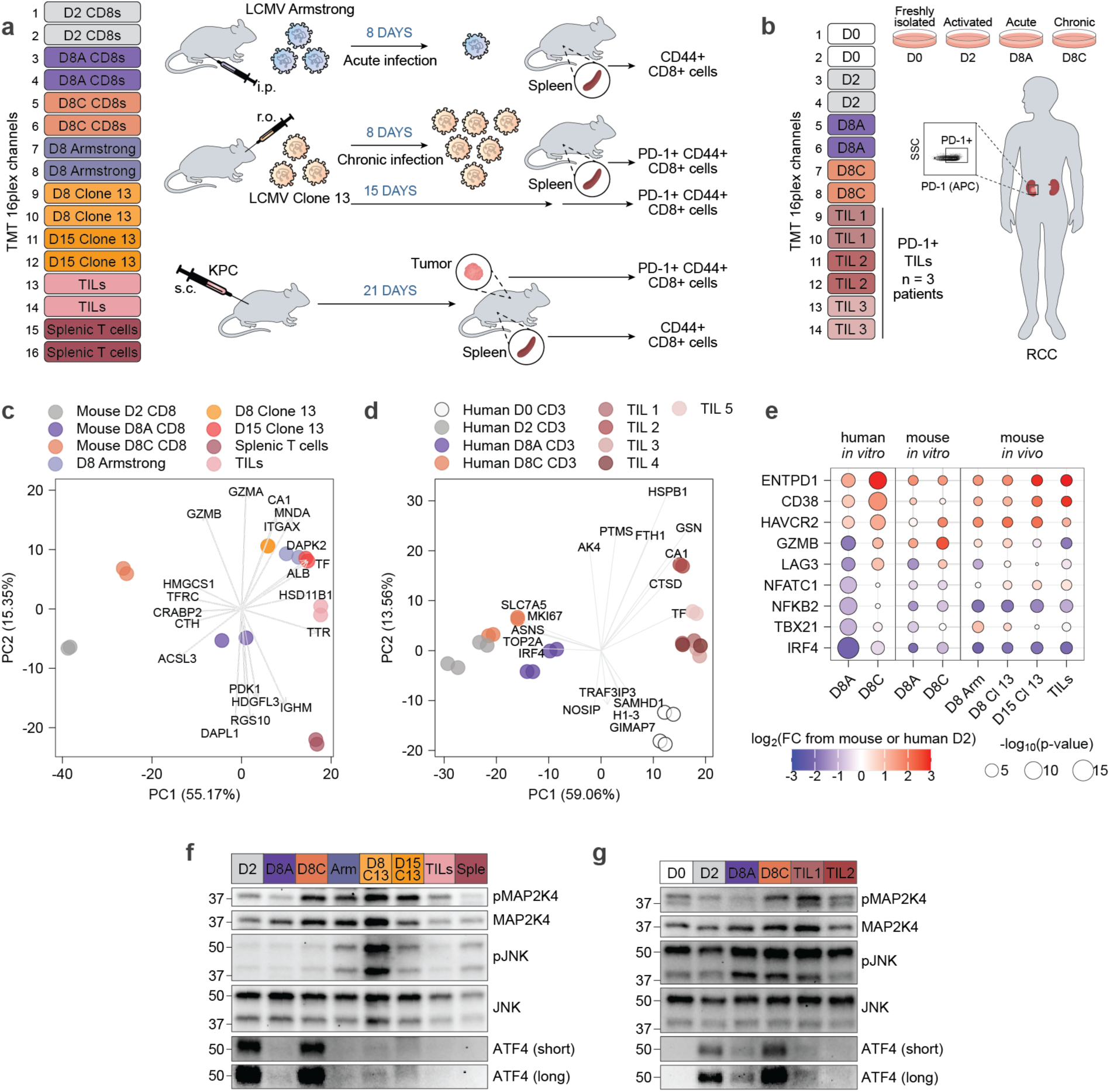
Complementary *in vivo* models of murine and human T cell exhaustion validate activation of MAP2K4-JNK pathway *in vivo*. (**a**) Schematic depicting experimental strategy to generate activated or exhausted CD8^+^ T cells from mice infected with acute or chronic strains of lymphocytic choriomeningitis virus (LCMV) infection as well as mice bearing subcutaneous KPC tumors in comparison with mouse T cells undergoing acute or chronic stimulation *in vitro*. Viably sorted CD8^+^ T cells with the indicated phenotypes were analyzed by low-input unenriched proteomics and Western blot analysis. (**b**) Schematic depicting the strategy to isolate human CD3^+^ T cells generated *in vitro* as described previously, as well as PD-1^+^ cells from renal cell carcinoma (RCC) patient tumors. Viable CD3^+^ T cells were sorted as shown. (**c**) PCA of unenriched proteomic data of *in vitro*-generated mouse D2, D8A, D8C T cells, and T cells isolated from mice bearing either LCMV infections or KPC tumors. Protein ratio values were log_2_ transformed and only proteins quantified in all experiments (4,343) were used. The five highest and five lowest loadings from PC1 and PC2 were plotted. (**d**) PCA of unenriched proteomic data of *in vitro*-generated human D0, D2, D8A, D8C T cells, and TILs isolated from five independent RCC patients. Protein ratio values were log_2_ transformed and only proteins quantified in all experiments (1,904) were used. The five highest and five lowest loadings from PC1 and PC2 were plotted. (**e**) Dot plot of log_2_FC and p-values of selected proteins detected in human and mouse T cells *in vitro* and *in vivo*. Values are relative to human or mouse D2 T cells. (**f**) Western blot analysis of *in vitro*-generated mouse T cells and T cells from mice bearing either LCMV infections or KPC tumors. (**g**) Western blot analysis of *in vitro*-generated human T cells and TILs from two independent RCC patients.

### Stress pathway activation limits the cytotoxic capacity of chronically stimulated T cells

Finally, we asked whether ER stress or stress kinase activation contributed substantially to the development of dysfunctional T cell state. To address this question, we treated activated T cells (D2) with either the reported MKK4/7 inhibitor BSJ-04-122^57^ or the ISR antagonist ISRIB^58^ for 6 days following primary T cell activation (Fig. 5a,b). We also activated ER stress by treating acutely or chronically stimulated T cells for 18 hours with thapsigargin^59^, which inhibits SERCA pumps and ER calcium stores leading to ER stress induction. Western blot analysis confirmed that thapsigargin was sufficient to induce ATF4 expression in acutely stimulated T cells, while ISRIB only minimally impacted eIF2α phosphorylation in chronically stimulated T cells (Extended Data Fig. 8a). BSJ-04-122 significantly reduced phosphorylation and activation of both MAP2K3 and MAP2K4 as well as their downstream targets p38 and JNK (Fig. 5c).

**Fig. 5.**
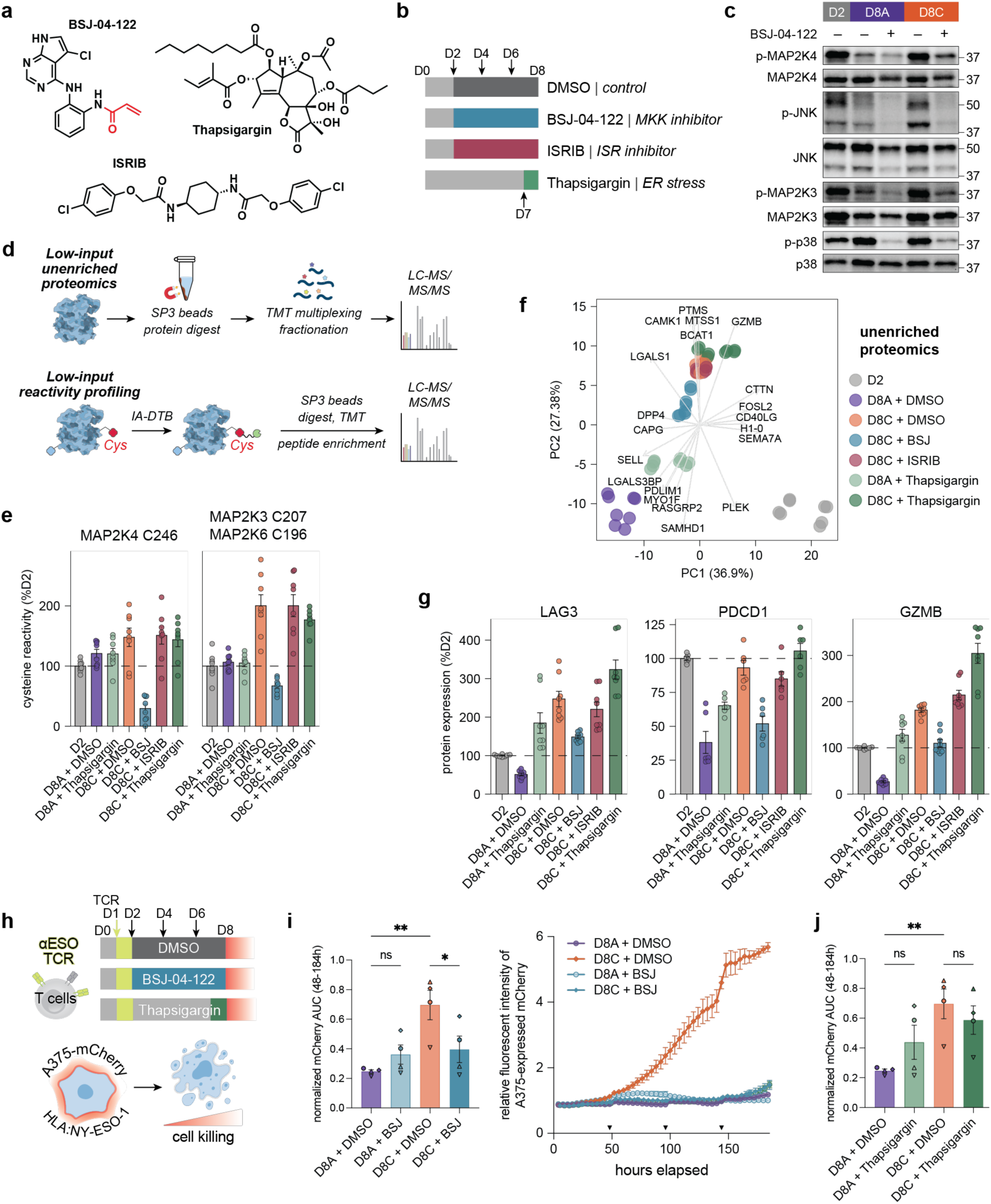
Pharmacologic targeting of stress kinase and ER stress responses modulates T cell dysfunction. (**a**) Chemical structures of ISRIB, BSJ-04-122, and thapsigargin. The covalent electrophilic group of BSJ-04-122 is highlighted in red. (**b**) Schematic of cytokine-mediated T cell expansion with or without chronic TCR stimulation and addition of small-molecule modulators of MAPK (10 µM BSJ-04-122), ISR (200 nM ISRIB), and ER stress (25 nM thapsigargin) pathways. (**c**) Western blot analysis of phosphorylation levels of MAP2K4 (Ser257), JNK (Thr183/Tyr185), MAP2K3 (Ser189), and p38 (Thr180/Tyr182) relative to protein expression in D2, D8A, and D8C T cells with or without 10 µM BSJ-04-122 treatment. (**d**) Schematic of low-input unenriched proteomics and cysteine reactivity profiling. See Materials and Methods for more details (Data S2-9, S2-10). (**e**) Cysteine reactivity profiling of MAP2K4 and MAP2K3/6 in D8A and D8C T cells with or without inhibitor treatments compared to D2 T cells (% of median D2 level). Data are presented as mean ± SEM; n = 4 donors (2 technical replicates per experimental condition). (**f**) PCA of unenriched proteomic data with or without inhibitor treatment from n = 4 donors (Data S2-9). Protein ratio values were log_2_ transformed and only proteins quantified in all experiments (4,198) were used. The five highest and five lowest loadings from PC1 and PC2 were plotted. (**g**) Bar graphs showing LAG3, PDCD1, and GZMB protein expression levels in D2, D8A, and D8C T cells with or without inhibitor treatments compared to D2 T cells (% of median D2 expression). Data are presented as mean ± SEM; n = 4 donors (2 technical replicates per experimental condition). (**h**) Schematic of antigen-specific cancer cell killing by D8A and D8C T cells transduced with NY-ESO-1-specific TCR and treated with DMSO, BSJ-04-122, or thapsigargin. (**i**) Left: Normalized AUC of mCherry intensity for 48-184 hours normalized to untreated D8A condition. Statistical comparison by one-way repeated measures ANOVA with Šídák’s multiple comparisons test (ns, p > 0.05; *, p < 0.05; **, p < 0.01). Donor-matched conditions are indicated by differently shaped data points. Right: A representative cell viability curve of NY-ESO-1-expressing A375 melanoma cells engineered to express mCherry following 184 hours of co-culture with T cells pre-treated with DMSO or BSJ-04-122, normalized to 0 h condition. Inverted triangles on x axis indicate timepoints when A375 melanoma cells were added to the co-culture. (**j**) Quantification of antigen-specific cancer cell killing by D8A and D8C T cells transduced with NY-ESO-1-specific TCR and treated with DMSO or thapsigargin. Normalized AUC of mCherry intensity for 48-184 hours normalized to untreated D8A condition. Statistical comparison by one-way repeated measures ANOVA with Šídák’s multiple comparisons test (ns, p > 0.05; **, p < 0.01). Donor-matched conditions are indicated by differently shaped data points. A representative cell viability curve is shown on the right.

We profiled these cells using a newly developed low-input proteomics platform, which leverages SP3 bead enrichment, enabling >20-fold reduction in protein input requirements for both unenriched proteomics and reactivity profiling experiments (Fig. 5d; Data S2-9, S2-10)^60^. We confirmed engagement of both MAP2K4_C246 and MAP2K3_C207 following BSJ-04-122 treatment, as observed by competition for their reactivity with this covalent electrophile (Fig. 5e and Extended Data Fig. 8b). Consistent with roles for both ER stress and MAP2K4, but not ISR, in driving T cell exhaustion, thapsigargin-treated D8A cells had increased proteomic similarity to D8C cells compared with DMSO-treated D8A cells, whereas BSJ-04-122-treated D8C cells were more similar to D8A cells compared with DMSO-treated D8C cells (Fig. 5f). Interrogation of specific proteins associated with exhaustion, including checkpoint receptors (PDCD1, LAG3, CTLA4), transcription factors (NFATC1, NR4A3, EGR2), and GZMB, showed an increase with thapsigargin treatment and a decrease with BSJ-04-122, which we validated by flow cytometry (Fig. 5g and Extended Data Fig. 8c,d). Finally, thapsigargin was sufficient to induce expression of proteins containing both ATF4 and ATF6 binding motifs in the absence of chronic TCR stimulation, whereas BSJ-04-122 attenuated chronic TCR stimulation-driven expression of the same proteins (Extended Data Fig. 8e). Taken together, these results are consistent with effective pharmacologic induction and reversal of T cell dysfunction by targeting ER stress and stress kinases, respectively.

Finally, we asked whether pharmacologic targeting of stress pathway or stress kinase activity would impact T cell effector function during serial tumor encounter. T cells transduced with the high-affinity NY-ESO-1-specific TCR were acutely or chronically stimulated for 6 days in the presence of either vehicle, BSJ-04-122, or thapsigargin (18 hours) and then co-cultured with NY-ESO-1 expressing A375 melanoma cancer cell line for an additional eight days, with replenishment of cancer cells every 2 days (Fig. 5h). BSJ-04-122 dramatically restored the cytotoxic capacity of chronically stimulated T cells as reflected by a significant reduction in the tumor cell abundance over 8 days of co-culture (Fig. 5h,i). In contrast, thapsigargin pre-treatment led to reduced tumor cell killing by acutely activated T cells (Fig. 5j). However, we cannot rule out the contribution of calcium signaling to thapsigargin-mediated T cell dysfunction because sustained intracellular calcium signaling induces T cell exhaustion by activating the calcineurin-NFAT signaling axis in the absence of cooperative AP-1 transcription factors^61^.

## DISCUSSION

In the present study, we leverage an *in vitro* platform of chronic antigen stimulation in peripheral blood human T cells from healthy donors to identify unique metabolic, proteomic, and post-translational alterations that are directly attributed to persistent TCR stimulation. To increase the accessibility of our *in vitro* platform, we developed an open-access portal, which displays the transcriptional, proteomic, and cysteine reactivity data for any queried gene across the activated, acutely, and chronically stimulated conditions. While this platform is unlikely to comprehensively capture all features of dysfunctional T cells within tumors, and particularly those features driven primarily by the tumor microenvironment, it offers a powerful and scalable experimental paradigm to identify specific proteomic and metabolomic changes driven by chronic TCR stimulation that we validated across orthogonal models of *in vivo* T cell exhaustion.

Our work identified unique metabolic features of human T cells during chronic TCR stimulation. Key amongst these features is the accumulation of mitochondrial NADH. In response to chronic stimulation, human T cells induce several adaptive mechanisms to limit mitochondrial ROS accumulation, diverting both carbon and reducing equivalents towards triglyceride, ether phospholipid, and proline biosynthesis at the expense of mitochondrial ATP synthesis, ultimately resulting in collapse of the nucleotide pool. Our reactivity profiling analyses identified functionally relevant changes during chronic stimulation that are directly connected to these metabolic defects. Notably, we identified and validated the MAP2K4-JNK axis as a targetable regulatory node capable of reducing the accumulation of exhaustion markers and restoring cytotoxic T cell function. While MAP2K4 has been previously reported to be activated in chronically stimulated human T cells, it had not been tested whether MAP2K4 inhibition could improve T cell function^27^.

Our results also point to broader functional consequences related to nucleotide imbalance, including mitochondrial protein homeostasis and proteasome activation. As a result of these and likely other changes, chronically stimulated T cells exhibit activation of both the integrated and ER stress responses, but not the mitochondrial UPR. Prior studies have established important roles for ER and proteotoxic stress in T cell dysfunction and exhaustion^13, 14, 27, 61, 62, 63, 64, 65^. Our findings extend this work by connecting some of these stress responses to nucleotide imbalance and identifying specific nucleotide-sensitive proteins and pathways that may contribute to their activation.

However, we do not believe that the global cysteine reactivity changes observed in our dataset are primarily driven by widespread protein misfolding. Several observations support this conclusion. First, the overall number and magnitude of cysteine-reactivity changes are relatively comparable between acutely and chronically stimulated T cells, despite protein misfolding and aggregation having been reported predominantly in chronically stimulated cells. If protein misfolding were the dominant driver of the observed reactivity changes, we would expect substantially more widespread alterations specifically in chronically stimulated T cells. Second, many proteins exhibit site-specific reactivity changes, with one cysteine displaying altered reactivity while additional quantified cysteines within the same protein remain unchanged, including MAP2K4 and HSPA9. This pattern is more consistent with localized conformational or regulatory changes than with global protein unfolding or aggregation. Third, protein aggregation would generally be expected to decrease probe accessibility and thus reduce cysteine labeling. In contrast, several of our key findings, including the increased reactivity of MAP2K3 and MAP2K4, represent gains in probe accessibility that are more consistent with activation-associated conformational remodeling than nonspecific protein misfolding. Finally, mass spectrometry-based detergent fractionation confirmed that chronically stimulated cells did not exhibit widespread protein accumulation in detergent-insoluble fractions.

Our work also reveals potential differences between human and mouse systems^13, 14^. Further mechanistic studies are necessary to elucidate the extent of mitochondrial protein homeostasis dysfunction and to determine why mtUPR activation is absent in our system. These investigations could provide deeper insights into the ability of human T cell mitochondria to maintain protein homeostasis under chronic stress conditions. Our results do not exclude a potential role for mitochondrial ROS in human T cell dysfunction; indeed, depolarized mitochondria have been identified in primary human TILs and linked to terminal T cell exhaustion. Mitochondrial ROS may emerge under continued chronic TCR stimulation over weeks or even months of tumor residence; furthermore, ROS may occur under *in vivo* environmental conditions that favor ROS generation, such as intratumoral hypoxia.

Our study validated cysteine reactivity profiling as an unbiased approach to investigate structural and functional proteomic changes across different immune cell states. While we identified nucleotide binding as a key biochemical feature that can be read out using reactivity profiling platforms, it is important to recognize that the enrichment and identification of cysteine-containing peptides can also be influenced by other direct or indirect post-translational modifications on the same tryptic peptides, including phosphorylation, ubiquitination, and fatty acylation, among others. We demonstrated that function-first approaches, including ATP add-back TMT-ABPP experiments with cysteine-reactive probes, serve as powerful strategies for annotating biochemical cues associated with specific reactivity changes. ATP add-back revealed differential ATP sensitivity of MAP2K3 and MAP2K4 between activated and chronically stimulated T cells, consistent with differences in nucleotide occupancy, as well as nucleotide-dependent regulation of proteasome activity, which correlated with intracellular ATP levels. However, it is worth noting that, because these experiments were conducted in cell lysates, where subcellular organization and physiological metabolite gradients are lost, they may not fully recapitulate the physiological roles of ATP in intact cells. Expanding function-first chemical proteomic platforms will deepen our understanding of the factors driving these reactivity changes.

We acknowledge several potential limitations to our experimental paradigm. Our proteomic analysis of *in vivo* exhausted T cells from mice and primary human TILs clearly indicates additional proteomic alterations that are environmentally encoded. Tumor-infiltrating T cells are subject to several cellular and environmental inputs *in vivo* that may contribute to their dysfunction, including antigens of varying TCR avidity, immunosuppressive stromal cells, paracrine factors including cytokines, and altered nutrient availability. We anticipate building upon this platform to understand how additional tumor-associated features synergize with chronic TCR stimulation to activate dysfunctional programs that are tissue-and/or tumor-specific. Finally, expanding reactivity profiling platforms to other nucleophilic amino acid residues, including lysine, arginine, and tyrosine, will provide further insights into proteins involved in T cell dysfunction following chronic TCR stimulation, considering their distinct abundances, physicochemical properties, and functional roles.

## Acknowledgements

We thank Taylor Chen, Sanraj Mittal, and Dr. Bryan Ngo for preliminary investigations. O.A.-W. is supported by the Neil S. Hirsch Foundation, Edward P. Evans Foundation, Break Through Cancer, NIH/NCI (R01 CA251138, R01 CA242020, R01 CA283364, and P50 CA254838), and NIH/NHLBI (R01 HL128239), and the Leukemia & Lymphoma Society. S.A.V was supported by an NCI K08 Career Development Award (NCI K08 CA237731). E.V.V. and S.A.V. were supported by a Damon Runyon-Rachleff Innovation Award. S.A.V., L.F.S.P, Y.-H.L., J.D.S., B.L, O.A.W and J.R. were additionally supported by a Cancer Center Support Grant (P30 CA008748). E.V.V. is also supported by Rockefeller University start-up funds, Robertson Foundation, The Achelis and Bodman Foundation, Herbert and Nell Singer Foundation, The Alexandrine and Alexander L. Sinsheimer Fund, Weill Cancer Hub East, Searle Scholar Program, The Stavros Niarchos Foundation Institute for Global Infectious Disease Research at Rockefeller, and CZ Biohub New York Investigator Program. J.L. and K.M.M. are supported by the SNF Institute for Global Infectious Disease Research, and J.L. is a Searle Scholar. H.K is supported by Pels Family Center Fellowship. C.R.W. is supported by the Marlene Hess Center for Research on Women’s Health and Biomedicine. L.F.S.P is supported by the MERIT Mandel Postdoctoral Fellow (MSKCC) and MSK Society Scholars Prize (MSKCC). Y.C. was supported by an F99/K00 transition award (F99CA294267). T.L.Z. was supported by NIH (T32GM136640-Tan to T.L.Z). K.N.K. was supported by the German Research Foundation, the Melanoma Research Foundation, and MSKCC’s Center for Experimental Immuno-Oncology. C.A.K. was supported by NIH (R37 CA259177, R01CA286507, P50CA217694, P30CA008748), The Parker Institute for Cancer Immunotherapy, Cycle for Survival, the Metropoulos Family Foundation, and the Samuel I. Newhouse Foundation. A.A.H. was supported in part by The Castellano Family and the Weiss Family Kidney Research Fund. Flow cytometry assays were optimized with guidance from Dr. Svetlana Mazel and the Rockefeller Flow Cytometry Resource Center (RRID:SCR_017694). Some analyses were performed on computational resources from the Rockefeller University High Performance Computing Resource Center (RRID: SCR_025889). We acknowledge the use of the Integrated Genomics Operation Core, funded by the NCI Cancer Center Support Grant (CCSG, P30 CA008748), Cycle for Survival, and the Marie-Josée and Henry R. Kravis Center for Molecular Oncology, the Weill Cornell Medicine Proteomics and Metabolomics Core Facility, and the Donald B. and Catherine C. Marron Cancer Metabolism Center at MSKCC.

## Author contributions

Conceptualization: H.K., C.R.W., S.A.V., E.V.V.

Data curation: H.S., B.Z., J.R., J.L., S.A.V., E.V.V.

Formal analysis: H.K., C.R.W., H.S., B.Z.

Investigation: H.K., C.R.W., T.-J.C., L.F.S.P., Y.-H.L., J.D.S., T.L.Z., C.R., Y.A., Y.C., L.M., A.A.H., S.A.V., E.V.V.

Methodology: H.K., C.R.W., T.-J.C., L.F.S.P., Y.-H.L., J.D.S., C.R., Y.C., K.N.K., A.A.H., S.A.V., E.V.V.

Validation: H.K., C.R.W., T.-J.C., L.F.S.P., J.D.S., S.A.V., E.V.V.

Resources: A.A.H., J.L., C.A.K., O.A.-W., S.A.V., E.V.V.

Software: H.S., B.Z., N.R., J.R., K.M.M.

Visualization: H.K., C.R.W., T.-J.C., H.S., B.Z., S.A.V., E.V.V.

Supervision: A.A.H., J.L., C.A.K., O.A.-W., S.A.V., E.V.V.

Writing – original draft, H.K., C.R.W., T.-J.C., L.F.S.P., H.S., S.A.V., E.V.V.

Writing – review & editing: all authors read, edited, and approved the final manuscript

## Competing interests

O.A.-W. is a founder and scientific advisor of Codify Therapeutics, holds equity and receives research funding from this company. O.A.-W. has served as a consultant for Amphista Therapeutics, and MagnetBio, and is on scientific advisory boards of Envisagenics Inc. and Harmonic Discovery Inc.; O.A.-W. received research funding from Astra Zeneca, Nurix Therapeutics, and Minovia Therapeutics, unrelated to this study. S.A.V. has provided consulting services for Generate:Biomedicines and has received research funding from Bristol Myers Squibb unrelated to the current work. E.V.V. is listed as a co-inventor on patents with Vividion Therapeutics. C.A.K. has consulted for or is on the scientific advisory boards for Achilles Therapeutics, Affini-T Therapeutics, Aleta BioTherapeutics, Bellicum Pharmaceuticals, Bristol Myers Squibb, Catamaran Bio, Cell Design Labs, Chronara Biosciences, Decheng Capital, G1 Therapeutics, Klus Pharma, Merck, Obsidian Therapeutics, PACT Pharma, Roche/Genentech, Royalty Pharma, Stereo Biotherapeutics, and T-knife, and is a scientific co-founder and equity holder in Affini-T Therapeutics. A.A.H. is on the advisory board for Merck. The remaining authors declare no competing interests.

## Data, code, and materials availability

Raw proteomic data and RNA-sequencing data have been deposited to PRIDE and NCBI GEO. Code for data processing and visualization and for the accompanying web portal will be deposited to GitHub. Any additional information required to reanalyze the data reported in this paper is available from the lead contact upon request. Questions and requests for chemical probes and plasmids generated in this study should be directed to and will be fulfilled by the lead contact with a completed materials transfer agreement.

## Extended data figures

**Extended Data Fig. 1.**
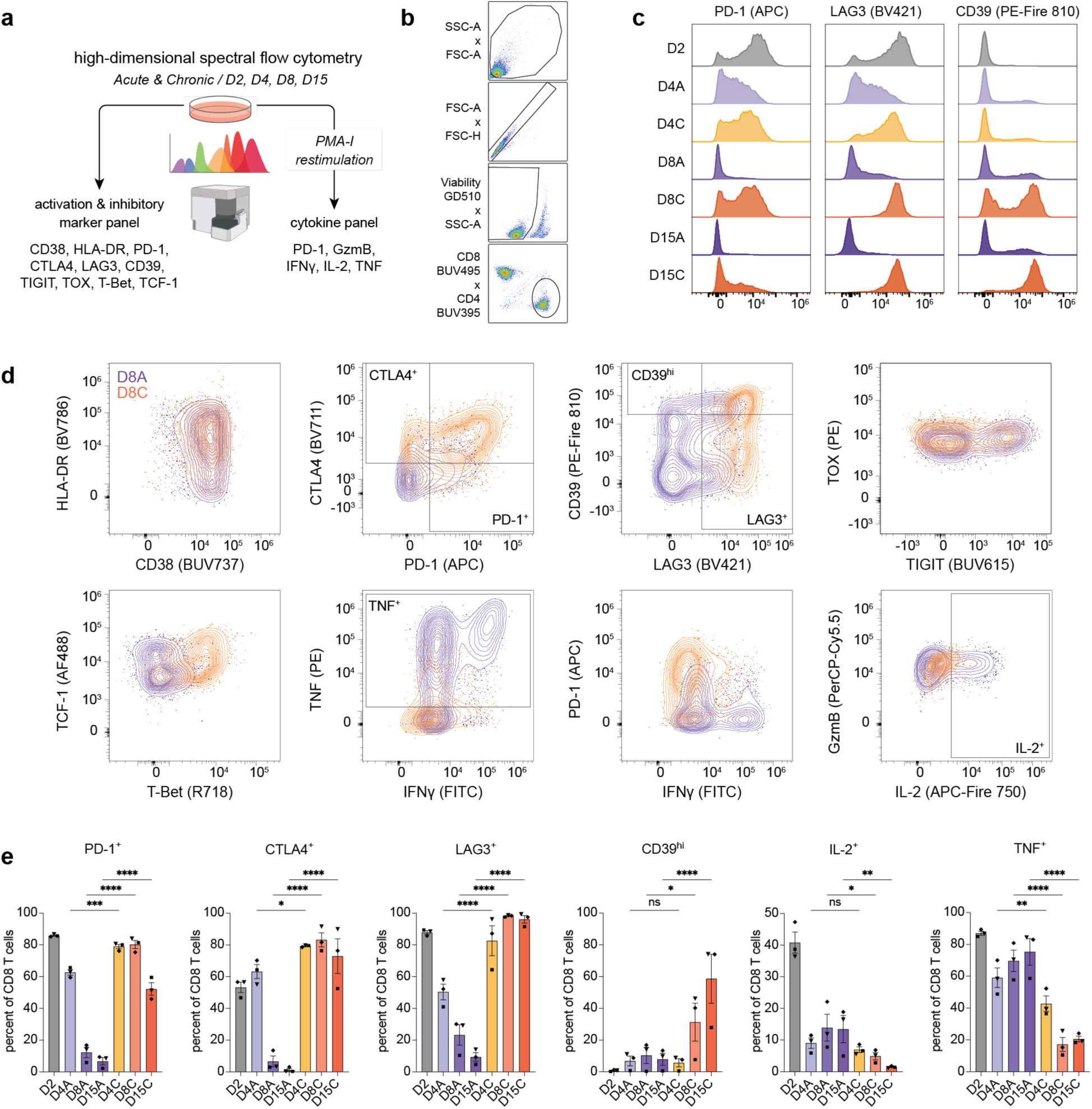
Flow cytometry-based validation of the *in vitro* T cell culturing system (Related to Figure 1). (**a**) Schematic of flow cytometry experiments used to profile markers of activation and T cell dysfunction over time. (**b-d**) Representative gating strategy (**b**), histograms (**c**), and contour plots (**d**) of activation/inhibitory marker and cytokine flow cytometry panels for CD8^+^ T cells from D8 acute (purple) and chronic (orange) conditions. Histograms are normalized to mode. Contour plots represent >3,000 cells per condition. (**e**) Quantification of T cell exhaustion markers and effector cytokines from flow cytometry time course experiments. Statistical comparison by one-way repeated measures ANOVA with Holm-Šídák’s multiple comparisons test of D4A vs D4C, D8A vs D8C and D15A vs D15C conditions (ns, p > 0.05; *, p < 0.05; **, p < 0.01; ***, p < 0.001; ****, p < 0.0001). Donor-matched conditions are indicated by differently shaped data points. Data are presented as mean ± SEM; n = 3.

**Extended Data Fig. 2.**
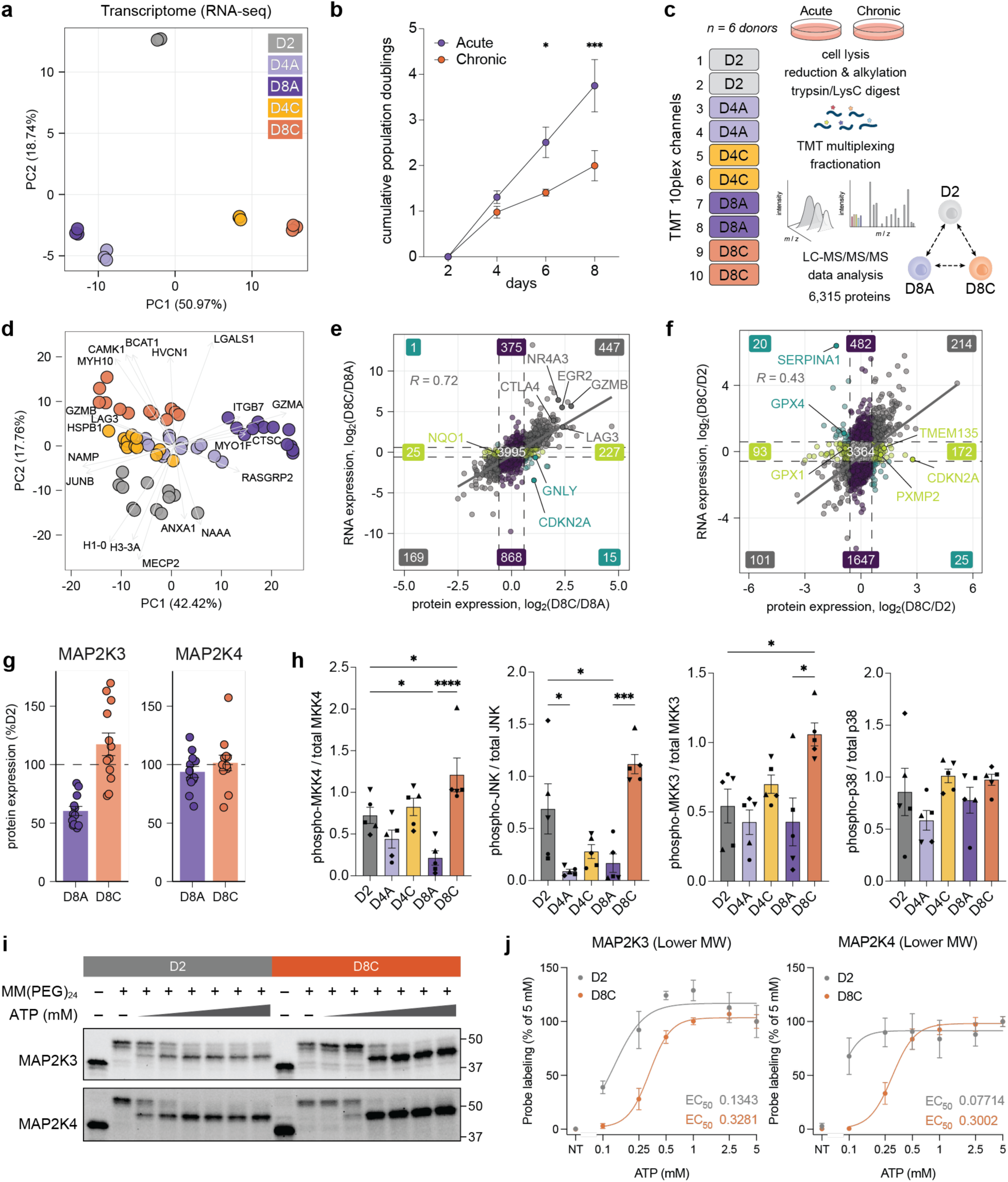
Transcriptomic and proteomic analyses of the *in vitro* T cell culturing system (Related to Figures 1 and 2). (**a**) Principal component analysis (PCA) of bulk RNA-sequencing from n = 3 donors (Data S1). The 5,000 protein-coding genes with the highest row-wise variance were used for PCA, and normalized count values were log_2_ transformed before PCA. (**b**) Population doublings of T cells cultured with or without chronic TCR stimulation from n = 3 independent donors compared by two-way ANOVA with Šídák’s multiple comparisons test (*, p < 0.05; ***, p < 0.001). (**c**) Experimental schematic of TMT-multiplexed unenriched proteomics of primary human T cells cultured with and without chronic TCR stimulation at three time points (Data S2-1). See Materials and Methods for more details. (**d**) PCA of unenriched proteomic data from n = 6 donors. Protein ratio values were log_2_ transformed and only proteins quantified in all experiments (3,615) were used. The five highest and five lowest loadings from PC1 and PC2 were plotted. (**e-f**) Log_2_ fold change of protein expression (x axis) and RNA expression (y axis) between D8C and D8A (**e**) and D8C and D2 (**f**) T cells. Comparison restricted to genes quantified in both proteomic and transcriptomic datasets (Data S2-2). Pearson’s correlation coefficient and regression line are shown. Dashed lines represent cutoffs of fold-change > 1.5. Transcriptomic data from n = 3 donors and proteomic data from n = 6 donors. (**g**) Bar graphs showing MAP2K3 and MAP2K4 protein expression levels in D8A and D8C T cells compared to D2 T cells (% of median D2 expression). Data are presented as mean ± SEM; n = 6 donors, 2 technical replicates per experimental condition. (**h**) Western blot quantifications of T cell lysates at D2, D4, and D8 timepoints. Statistical comparisons by one-way repeated measures ANOVA with Holm-Šídák’s multiple comparisons test of D4A vs D4C, D8A vs D8C, and D2 vs D8C (ns, p > 0.05; *, p < 0.05; ***, p < 0.001; ****, p < 0.0001). Donor-matched conditions are indicated by differently shaped data points. Phosphorylation levels of MAP2K4 at Ser257 normalized by total MAP2K4; of JNK at Thr183 and Tyr185 normalized by total JNK; of MAP2K3 at Ser189 normalized to total MAP2K3, and of p38 at Thr180 and Tyr182 normalized to total p38. Data are presented as mean ± SEM; n = 4-5. (**i**) ATP dose-dependent response for reactive cysteines in MAP2K3 and MAP2K4 using activity-based protein profiling with a high-molecular weight cysteine-reactive maleimide probe. ATP concentrations of 0, 0.1, 0.25, 0.5, 1, 2.5, and 5 mM were tested using Western blot visualization (WB-ABPP; representative of three independent donors). (**j**) Quantification of lower molecular weight bands of MAP2K3 and MAP2K4 across ATP concentrations, displayed with best-fit nonlinear curve and apparent EC_50_ values, n = 3 donors. ATP concentration of 0 mM is indicated by NT (not treated).

**Extended Data Fig. 3.**
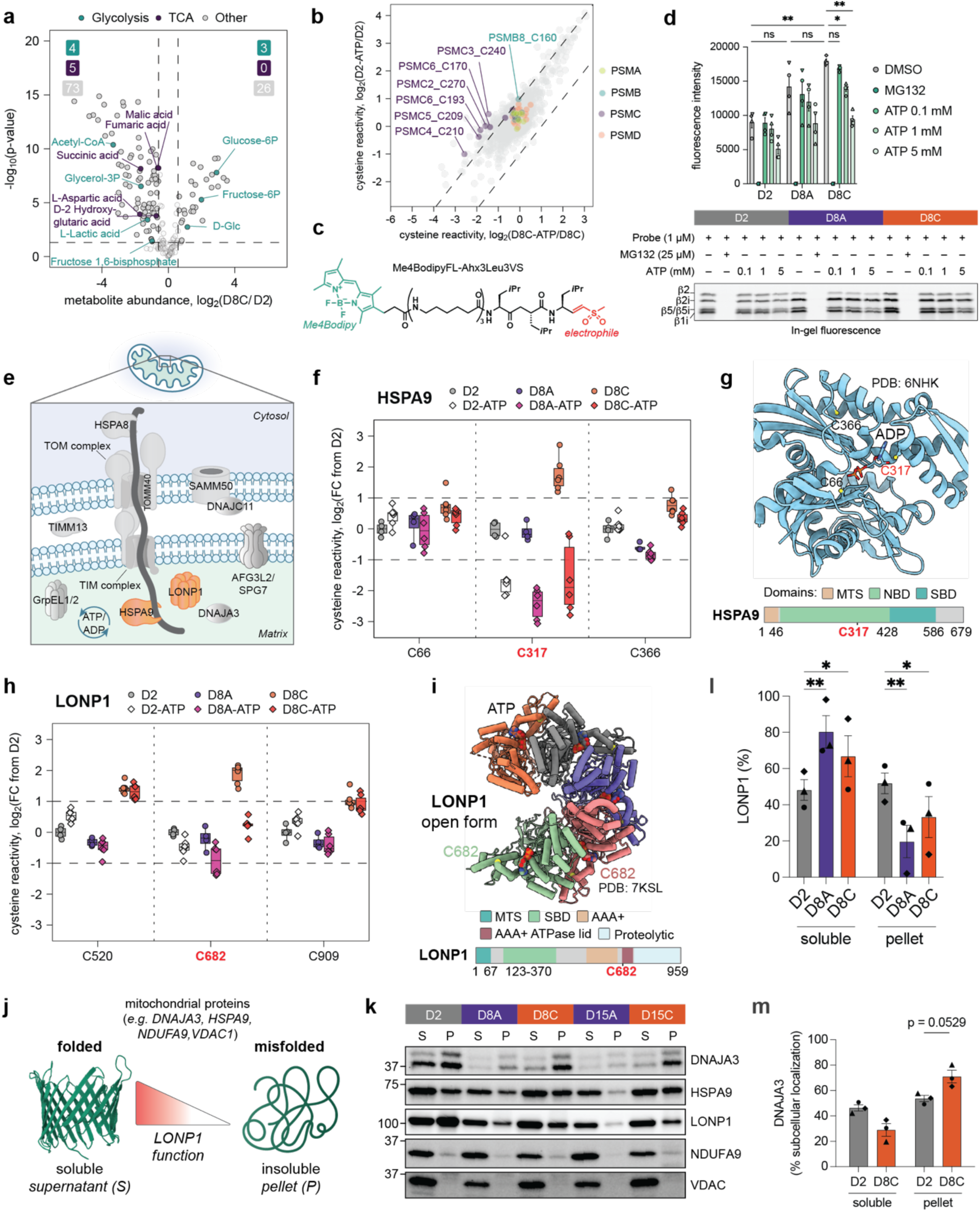
Identification of ATP-sensitive cysteine reactivity changes during chronic TCR stimulation (Related to Figure 2). (**a**) Volcano plots showing log_2_ fold changes of metabolite abundances between D8C and D2 T cells. Metabolites are colored based on manually curated pathway lists retrieved from KEGG. Dashed lines represent cutoffs of p-value < 0.05 and fold-change > 1.5. Data are from n = 4 donors. (**b**) Scatter plot of log_2_ fold changes in cysteine reactivity between D8C cells with ATP add-back compared to D8C cells without ATP add-back (x axis) and log_2_ fold change in cysteine reactivity between D2 cells with ATP add-back compared to D2 cells without ATP add-back (y axis). Colors show proteasome subunit families PSMA, PSMB, PSMC, and PSMD. Data are from n = 2 donors analyzed across two independent cysteine reactivity profiling experiments. (**c**) Chemical structure of a fluorescent proteasome activity probe, Me_4_BodipyFL-Ahx_3_Leu_3_VS. (**d**) Top: Quantification of active proteasome subunit labeling in cell lysates of D2, D8A, and D8C T cells. Statistical comparisons by two-way repeated measures ANOVA with Šídák’s multiple comparisons test of D2 vs D8A, D2 vs D8C, and D8A vs D8C, as well as D8C + DMSO vs D8C + 0.1, 1 or 5 mM ATP (ns, p>0.05; *, p < 0.05; **, p < 0.01). Donor-matched conditions are indicated by differently shaped data points. Bottom: gel-based proteasome activity profiling of D2, D8A, and D8C T cells. Cell lysates were preincubated with 25 µM MG132 or ATP (0.1, 1 or 5 mM), followed by incubation with 1 µM Me_4_BodipyFL-Ahx_3_Leu_3_VS. (**e**) Schematic overview of mitochondrial protein import and degradation pathways, including HSPA9 and LONP1. (**f**) Boxplot of log_2_ fold changes in cysteine reactivity of HSPA9 in control (circle) and ATP add-back (diamond) conditions in D2, D8A, and D8C T cells compared to D2 control condition. Boxes mark the lower and upper quartiles, horizontal black lines mark the median value, and whiskers extend to the furthest point within 1.5x the interquartile range. (**g**) Crystal structure of HSPA9 in ADP-bound state (PDB: 6NHK^1^) with quantified cysteines C66 and C366 labeled in black, and differentially reactive C317 labeled in red. (**h**) Boxplot of log_2_ fold changes in cysteine reactivity of LONP1 in control (circle) and ATP add-back (diamond) conditions in D2, D8A, and D8C T cells compared to D2 control condition. Boxes mark the lower and upper quartiles, horizontal black lines mark the median value, and whiskers extend to the furthest point within 1.5x the interquartile range. (**i**) Electron microscopy structure of the open form of LONP1 (PDB: 7KSL _2_) with labeling of differentially reactive C682. (**j**) Schematic of detergent-soluble and insoluble fractions for identification of misfolded or aggregated proteins in activated, acutely, and chronically stimulated T cells ^3^. (**k**) Comparison of solubility of mitochondrial proteins in D2, D8A, D8C, D15A, and D15C T cells. Cell lysate was separated into 1% Triton X-100-soluble fraction (S) and insoluble pellet (P), and protein levels were analyzed by Western blot. (**l**) Quantification of the distribution of LONP1 between soluble and insoluble fractions in activated, acutely, and chronically stimulated T cells. Statistical comparison by one-way repeated measures ANOVA with Šídák’s multiple comparisons test (*, p < 0.05; **, p < 0.01; non-significant comparisons not displayed). Donor-matched conditions are indicated by differently shaped data points. Data are presented as mean ± SEM; n = 3. (**m**) Quantification of the distribution of DNAJA3 between soluble and insoluble fractions in activated and chronically stimulated T cells. Statistical comparison by two-tailed paired t test (ns, p > 0.05). Donor-matched conditions are indicated by differently shaped data points. Data are from n = 3 donors.

**Extended Data Fig. 4.**
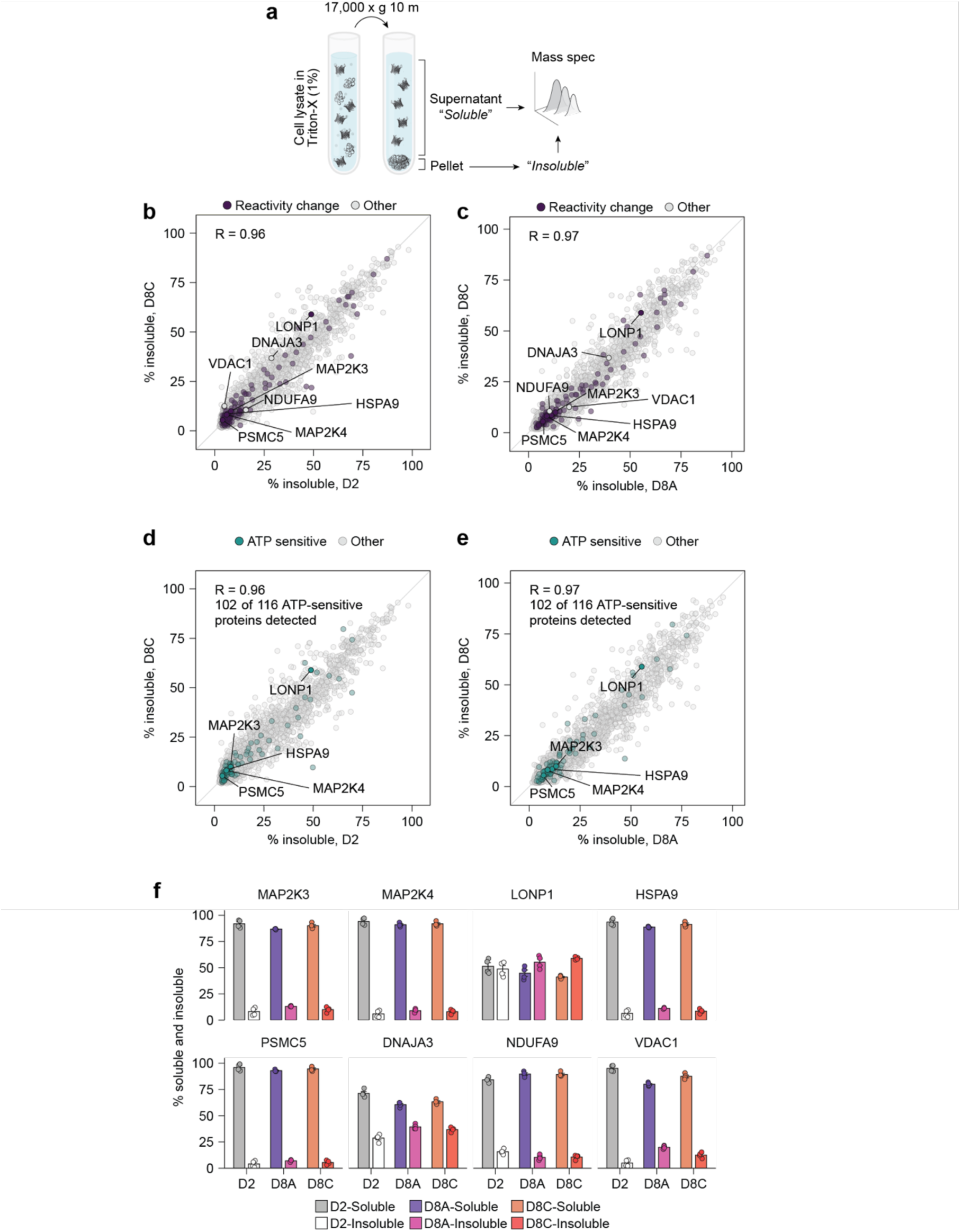
Cysteine reactivity changes are not attributable to loss of proteostasis in chronically stimulated cells (Related to Figure 2). (**a**) Experimental schematic showing separation of soluble and insoluble fractions by centrifugation, followed by mass spectrometry analysis. (**b-e**) Percentage of proteins in insoluble fractions (100 × I/(I + S); calculated from mean channel ratios) in D8C versus D2 (b,d) and D8C versus D8A (c,e) T cells. Only proteins quantified in both donors of every state × fraction group are shown. (b, c) Proteins with identified cysteine reactivity changes are shown in purple, without – in grey. Diagonal line is the line of identity. (d,e) Proteins with ATP-sensitive cysteines are shown in teal, without – in grey. (**f**) Mean soluble and insoluble percentage for each protein in D2, D8A, and D8C T cells from unenriched proteomics experiments of detergent-fractionated cells. Error bars are SEM and points are TMT channels.

**Extended Data Fig. 5.**
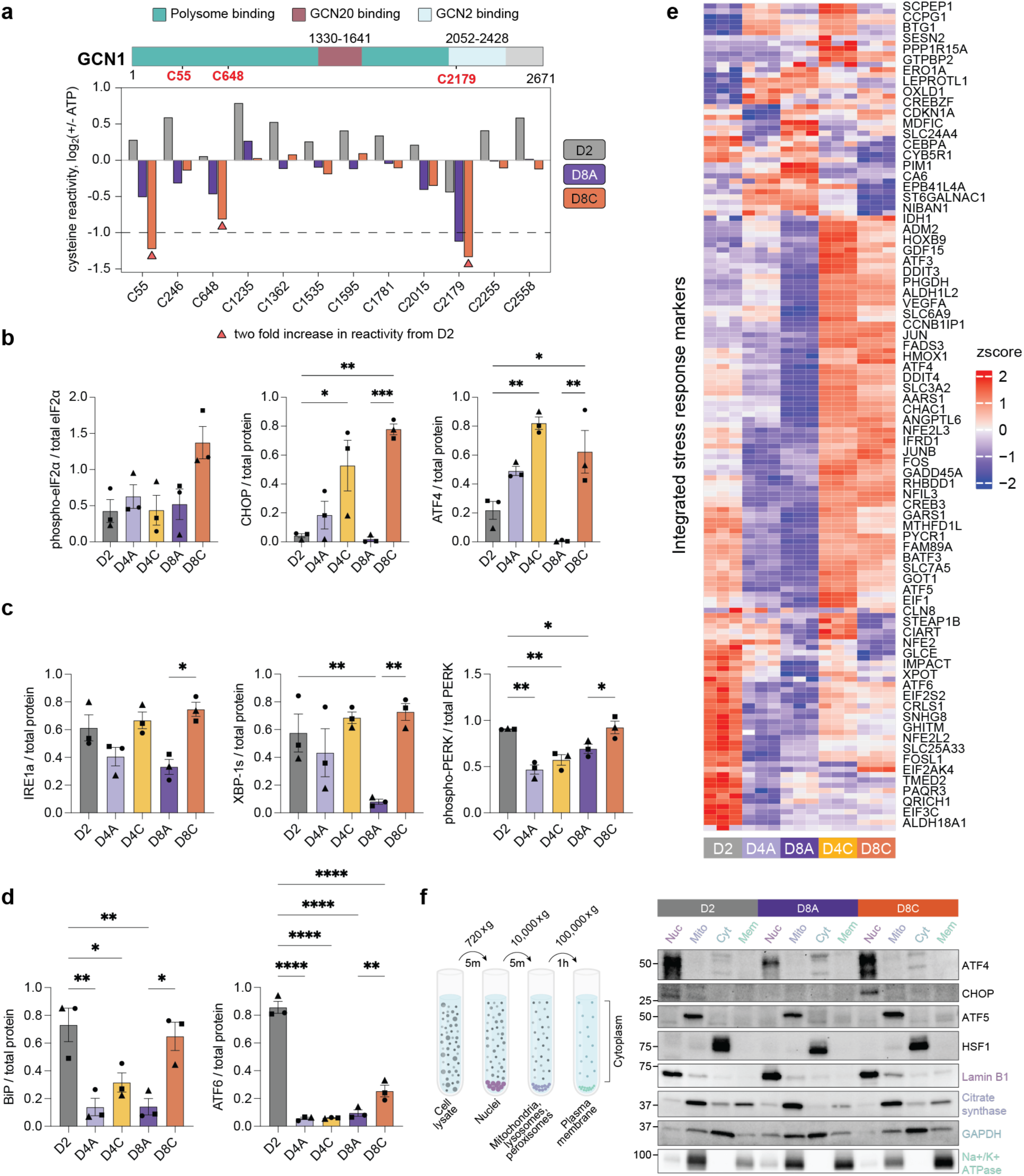
Differential activation of integrated, mitochondrial, and ER stress responses during chronic TCR stimulation (Related to Figure 3). (**a**) Bar graph of log_2_ fold changes in cysteine reactivity of GCN1 in D2, D8A, and D8C cells between control and ATP add-back conditions. Red triangles below bars denote a cysteine showing a general reactivity change (FC > 2) compared to D2 cells. Data are from n = 2 independent cysteine reactivity profiling experiments. Sequence of GCN1 with annotated polysome-, GCN20-, and GCN2-binding domains ^4, 5^. (**b**) Western blot quantifications of ISR proteins in T cell lysates at D2, D4, and D8 timepoints. Statistical comparisons by one-way repeated measures ANOVA with Holm-Šídák’s multiple comparisons test of D4A vs D4C, D8A vs D8C, and D2 vs D8C (*, p < 0.05; **, p < 0.01, ***, p < 0.001). Donor-matched conditions are indicated by differently shaped data points. ATF4 and CHOP were normalized to total protein level (n = 3). Phosphorylation levels of eIF2α at Ser51 were normalized by total eIF2α. (**c-d**) Western blot quantifications of ER-stress proteins in T cell lysates at D2, D4, and D8 timepoints. Statistical comparisons by one-way repeated measures ANOVA with Holm-Šídák’s multiple comparisons test of D2 versus each condition, D4A vs D4C, and D8A vs D8C (*, p < 0.05; **, p < 0.01, ****, p < 0.0001; non-significant comparisons not displayed). Donor-matched conditions are indicated by differently shaped data points. Data are presented as mean ± SEM; n = 3. (**e**) Heatmap of bulk RNA-Seq expression of ISR genes ^6^. Every second gene is labeled. (**f**) Schematic overview of subcellular fractionation of primary human T cells using differential centrifugation (left) and Western blot analysis of subcellular localization of transcription factors involved in ISR or mtUPR (right).

**Extended Data Fig. 6.**
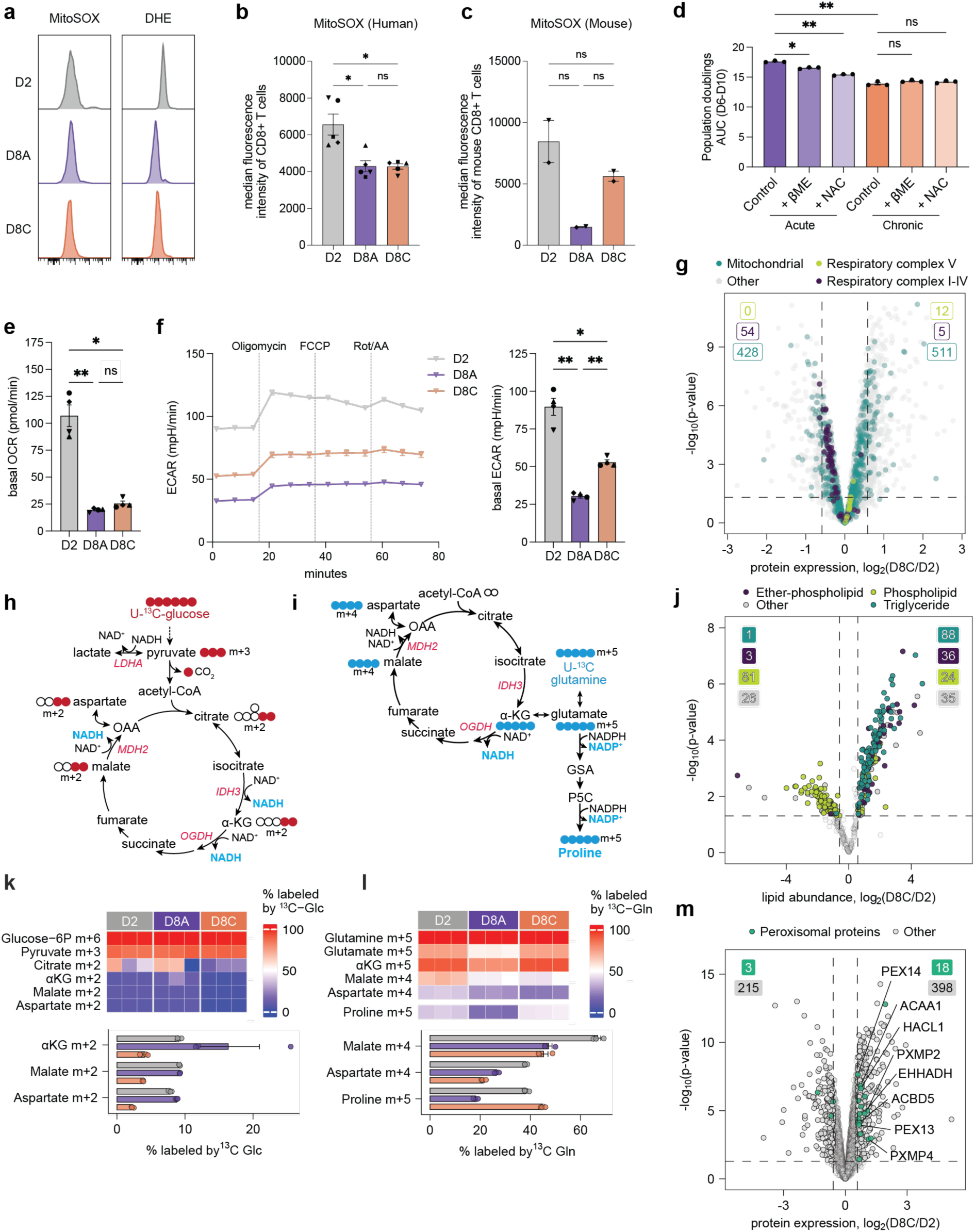
Metabolic and proteomic profiling highlights reduced mitochondrial NAD^+^ regeneration, increased non-oxidative TCA cycle metabolism and activation of nucleotide salvage in chronically stimulated T cells (Related to Figure 3). (**a**) Representative histograms of oxidative stress markers in human CD8^+^ T cells by flow cytometry using two ROS-reactive dyes: mitochondrial superoxide indicator (mitoSOX) and dihydroethidium (DHE; pan-cellular superoxides). (**b-c**) Quantification of oxidative stress in human (**b**) and mouse (**c**) CD8^+^ T cells by flow cytometry using mitochondrial superoxide indicator (mitoSOX). For human samples: Statistical comparison by repeated measures one-way ANOVA with Šídák’s multiple comparisons test (ns, p > 0.05; *, p < 0.05; ***, p < 0.001; ****, p < 0.0001). Donor-matched conditions are indicated by differently shaped data points. Data are from n = 5 donors. For mouse samples: statistical comparison by one-way ANOVA (MitoSOX) with Šídák’s multiple comparisons test (ns, p > 0.05). Subject-matched conditions are indicated by differently shaped data points. Data are from n = 2 mice. (**d**) Area under the curve (AUC) of population doubling in human T cells after treatment with ROS scavenger compounds β-mercaptoethanol (βME, 50 µM) or *N*-acetylcysteine (NAC, 5 mM). Statistical comparison by repeated measures one-way ANOVA (ns, p > 0.05, *, p < 0.05, **, p < 0.01). (**e**) Quantification of oxygen consumption rate (OCR) across n = 4 independent donors. Statistical comparison by one-way repeated measures ANOVA with Tukey’s multiple comparisons test (ns, p > 0.05; *, p < 0.05; **, p < 0.01). Donor-matched conditions are indicated by differently shaped data points; data are presented as mean ± SEM. (**f**) Representative plot of extracellular acidification rate (ECAR) as measured by extracellular flux analysis in D2, D8A, and D8C cells. Injections of oligomycin (ATP synthase inhibitor), FCCP (mitochondrial uncoupler), and rotenone/antimycin A (electron transport chain inhibitors) are indicated by vertical lines. Quantification across n = 4 independent donors is shown on the right. Statistical comparison by one-way repeated measures ANOVA with Tukey’s multiple comparisons test (ns, p > 0.05; *, p < 0.05; **, p < 0.01). Donor-matched conditions are indicated by differently shaped data points; data are presented as mean ± SEM. (**g**) Volcano plot showing log_2_ fold change of proteins between D8C and D2 T cells. Mitochondrial proteins and respiratory complexes I-V are labeled based on Gene Ontology Cellular Component annotation. (**h**) Schematic depicting how oxidative metabolism of [U-^13^C] glucose generates labeled metabolites associated with the TCA cycle. Red circles represent ^13^C-labeled carbons. (**i**) Schematic depicting how oxidative and non-oxidative metabolism of [U-^13^C] glutamine generates labeled metabolites associated with the TCA cycle. Blue circles represent ^13^C-labeled carbons. (**j**) Heatmap (top) and bar graph (bottom) showing fractional labeling by [U-^13^C] glucose of indicated metabolites in D2, D8A, and D8C T cells. Representative data from one of two independent donors (n = 3 replicates/donor) showing similar results. Data are presented as mean ± SEM. (**k**) Heatmap (top) and bar graph (bottom) showing fractional labeling by [U-^13^C] glutamine of indicated metabolites in D2, D8A, and D8C T cells. Representative data from one of two independent donors (n = 3 replicates/donor) showing similar results. Data are presented as mean ± SEM. (**l**) Volcano plot showing log_2_ fold changes in lipid abundances between D8C and D2 T cells. Dashed lines represent cutoffs of p-value < 0.05 and fold-change > 1.5. Data are from n = 2 donors. (**m**) Volcano plot showing log_2_ fold change of proteins between D8C and D2 T cells. Peroxisomal proteins are labeled based on Gene Ontology Cellular Component annotation.

**Extended Data Fig. 7.**
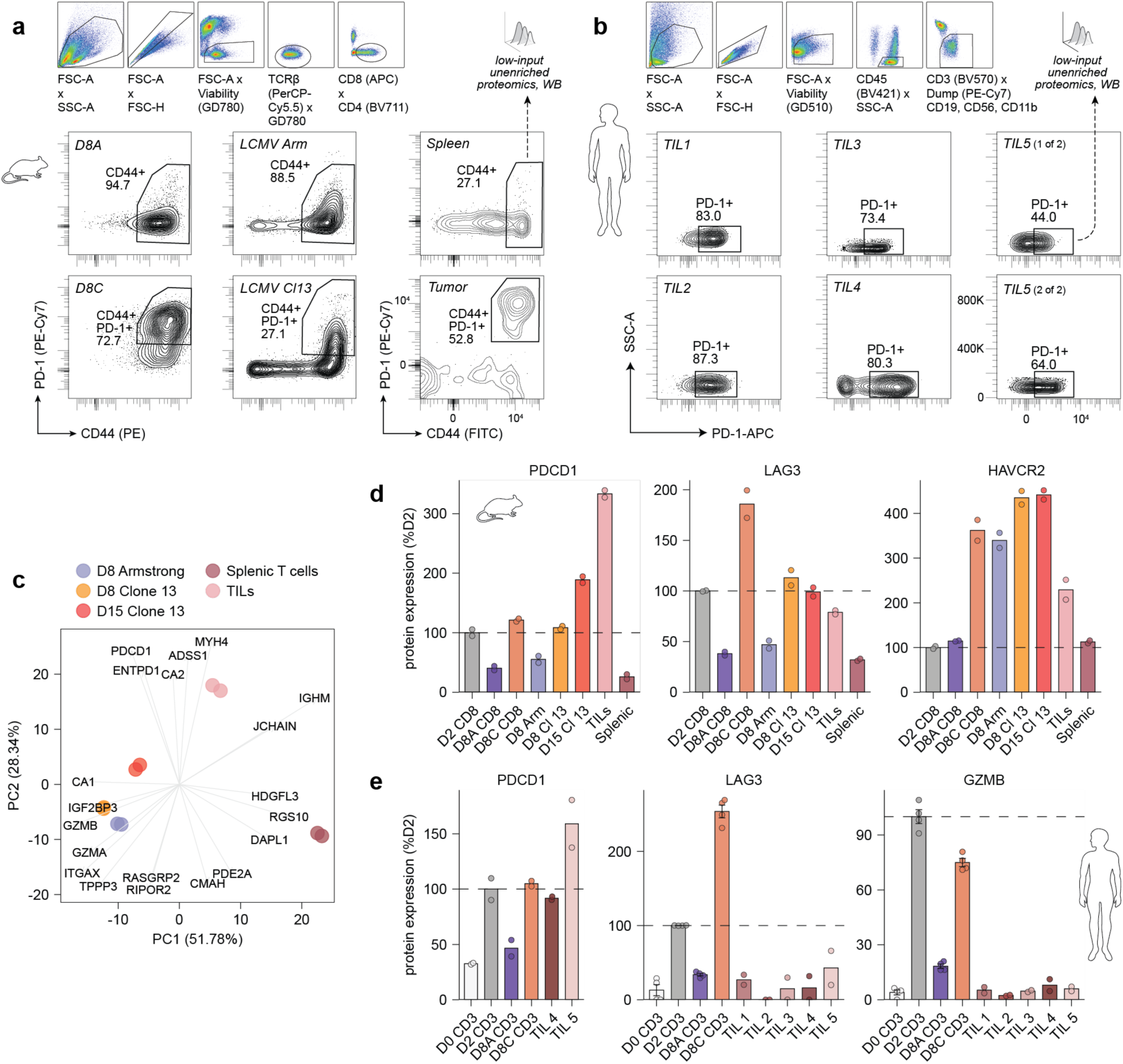
Phenotypic validation of T cell dysfunction in exhausted T cells from mouse and human samples (Related to Figure 4). (**a**) Gating strategy for mouse T cells sorted as shown in Fig. 4a. Contour plots below show representative CD44 and PD-1 expression for individual conditions. (**b**) Gating strategy for human T cells sorted as shown in Fig. 4b. Contour plots below show representative PD-1 expression for individual conditions. (**c**) PCA of unenriched proteomic data of T cells isolated from mice bearing either LCMV infections or KPC tumors. Protein ratio values were log_2_ transformed and only proteins quantified in all experiments (4,343) were used. The five highest and five lowest loadings from PC1 and PC2 were plotted. (**d**) Bar graphs showing the protein expression levels of T cell exhaustion markers, including PDCD1, LAG3, and HAVCR2 in *in vitro* and *in vivo* mouse T cells compared to *in vitro*-generated D2 T cells (% of mean D2 expression). Data are presented as mean; n = 1 donor (2 technical replicates per experimental condition). (**e**) Bar graphs showing the protein expression levels of T cell exhaustion markers, including PDCD1, LAG3, and GZMB in TILs from RCC patients compared to *in vitro*-generated D2 T cells (% of mean D2 expression). Data are presented as mean ± SEM; n = 5 donors (2 technical replicates per experimental condition).

**Extended Data Fig. 8.**
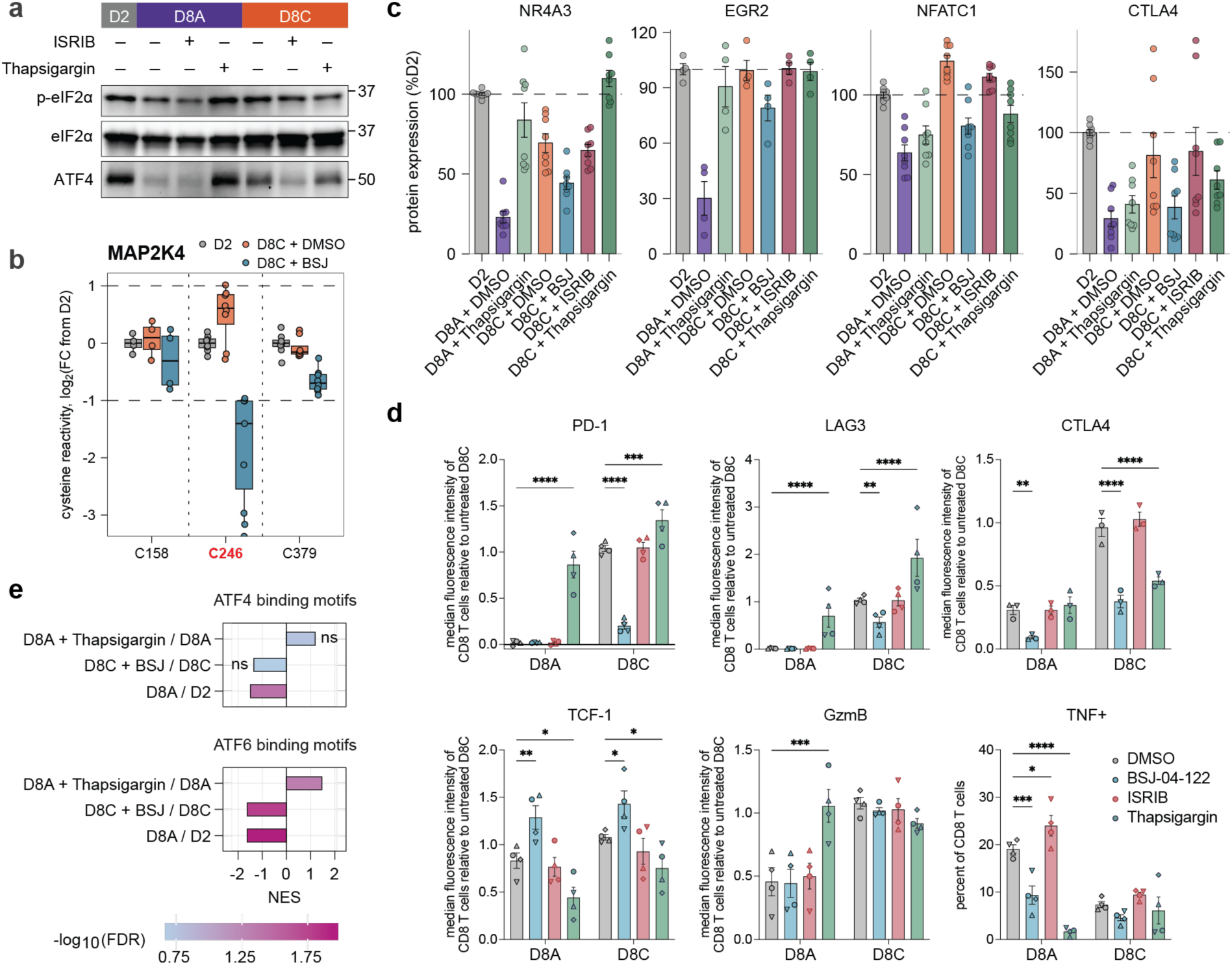
Characterization of T cell phenotypes following pharmacologic targeting of stress kinases, ISR, and ER stress response pathway (Related to Figure 5). (**a**) Western blot analysis of phospho-eIF2α and ATF4 in D2, D8A, and D8C T cells with or without the indicated inhibitor treatments. (**b**) Boxplot of log_2_ fold change in cysteine reactivity of MAP2K4 protein between D8C and D2 T cells with or without BSJ treatment. Cysteines with reactivity changes are highlighted in red. Boxes mark the lower and upper quartiles, horizontal black lines mark the median value, and whiskers extend to the furthest point within 1.5x the interquartile range. Data are from n = 4 donors analyzed across four independent cysteine reactivity profiling experiments. (**c**) Bar graphs showing the protein expression levels of T cell exhaustion markers, including NR4A3, EGR2, NFATC1, and CTLA4 in D8A and D8C T cells with or without inhibitor treatments compared to D2 T cells (% of median D2 expression). Data are presented as mean ± SEM; n = 4 donors (2 technical replicates per experimental condition). (**d**) Quantification by flow cytometry of CD8^+^ T cell expression of key markers of T cell exhaustion. Data for PD-1, LAG3, CTLA4, TCF-1, GzmB, and TNF presented as median fluorescent intensity (MFI) relative to the MFI of untreated D8C cells from the matched donor. Comparison by two-way repeated measures ANOVA with Dunnett’s multiple comparisons test (*, p < 0.05; **, p < 0.01; ***, p < 0.001; ****, p < 0.0001). Each condition compared to DMSO-treated control; non-significant comparisons not displayed. Data are presented as mean ± SEM; n = 4 donors. Donor-matched conditions are indicated by differently shaped data points. (**e**) GSEA of unenriched proteomics data from T cells on the hallmark gene sets ATF4 and ATF6 binding motifs. Comparisons not passing an FDR < 0.05 are labelled n.s. Unenriched proteomics data are from n = 4 donors.

## Supplementary Information

### Data S1

**Table S1-1:** Differential gene expression analysis from DESeq2. RNA counts were normalized using DESeq2’s median of ratios method. Genes with zero reads mapping are not displayed. Data are from n = 3 donors.

| # | Title | Description |
| --- | --- | --- |
| 1 | S1-1 Bulk RNA-Sequencing data | Differential gene expression analysis of bulk RNA-Seq data from activated, acutely, and chronically stimulated T cells (related to Fig. 1 and Extended Data Fig. 1). |

## Data S2

**Table S2-1:** TMT-exp data showing protein expression changes in human T cells cultured for 4 or 8 days with or without chronic stimulation compared to activated D2 T cells. Data are from n = 6 donors. **Table S2-2:** TMT-exp and bulk RNA-seq data showing gene and protein expression changes in human T cells cultured for 4 or 8 days with or without chronic stimulation compared to activated D2 human T cells. **Table S2-3:** Proteins with identified cysteine reactivity changes in at least one condition (D4A, D8A, D4C, D8C) compared to D2 human T cells. Data are median from n = 5 donors. **Table S2-4:** Reactivity fold change values versus protein expression fold change values for human T cells cultured for 4 or 8 days with or without chronic stimulation compared to activated D2 human T cells. **Table S2-5:** Cysteine reactivity values for human T cells in D2, D8A, and D8C stimulation conditions with and without the addition of ATP. Data are from n = 2 donors. **Table S2-6:** Unenriched proteomics of the soluble and insoluble fractions of activated T cells and T cells cultured for 8 days with or without chronic stimulation. Data are normalized such that the averages of the soluble percentage and insoluble percentage sum to 100%. Data are from n = 2 donors. **Table S2-7:** Low-input TMT-exp data showing protein expression changes in CD8^+^ T cells from mice infected with acute or chronic strains of lymphocytic choriomeningitis virus (LCMV) infection as well as mice bearing subcutaneous KPC tumors in comparison with mouse T cells undergoing acute or chronic stimulation *in vitro*. Data is shown normalized to *in vitro*-generated D2 cells. **Table S2-8:** Low-input TMT-exp data showing protein expression changes in *in vitro*-generated acutely or chronically stimulated human CD3^+^ T cells and PD-1^+^ cells isolated from five independent renal cell carcinoma (RCC) patients compared to *in vitro*-generated D2 cells. Data are from n = 5 patients. **Table S2-9:** Low-input TMT-ABPP data for human T cells cultured for 8 days under chronic stimulation with or without inhibitor treatment compared to activated D2 cells. Data are from n = 4 donors. **Table S2-10:** Low-input TMT-exp data showing protein expression changes in human T cells cultured for 8 days under chronic stimulation with or without inhibitor treatment compared to activated D2 cells. Data are from n = 4 donors.

| # | Title | Description |
| --- | --- | --- |
| 1 | S2-1 Unenriched proteomics | Whole proteome (TMT-exp) data from activated, acutely, and chronically stimulated human T cells (related to Fig. 1 and Extended Data Fig. 2, 6) |
| 2 | S2-2 Bulk RNA vs WP | Whole proteome (TMT-exp) and transcriptome (bulk RNA-Seq) data (related to Extended Data Fig. 2) |
| 3 | S2-3 Reactivity changes | Cysteine reactivity changes identified from IA-DTB reactivity TMT-ABPP data (related to Fig. 1, 2) |
| 4 | S2-4 Cysteine reactivity vs WP | Cysteine reactivity values (TMT-ABPP) versus whole proteome (TMT-exp) values for D4A, D8A, D4C, and D8C human T cells (related to Fig. 1) |
| 5 | S2-5 ATP add-back | Cysteine reactivity values (TMT-ABPP) for activated, acutely, and chronically stimulated human T cells with and without addition of ATP (related to Fig. 2, 3 and Extended Data Fig. 3, 5) |
| 6 | S2-6 Detergent solubility | Unenriched proteomics data from the soluble and insoluble fractions of activated, acutely, and chronically stimulated T cells (related to Extended Data Fig. 4) |
| 7 | S2-7 Low-input WP, mouse | Low-input whole proteome (TMT-exp) data from *in vitro*-generated mouse D2, D8A, D8C T cells, and T cells isolated from mice bearing either LCMV infections or KPC tumors (related to Fig. 4 and Extended Data Fig. 7) |
| 8 | S2-8 Low-input WP, TILs | Low-input whole proteome (TMT-exp) data from *in vitro*-generated human D0, D2, D8A, D8C T cells, and TILs isolated from five independent RCC patients (related to Fig. 4 and Extended Data Fig. 7) |
| 9 | S2-9 Low-input RC, inhibitors | Low-input cysteine reactivity (TMT-ABPP) for activated, acutely, and chronically stimulated human T cells with and without inhibitor treatment (related to Fig. 5 and Extended Data Fig. 8) |
| 10 | S2-10 Low-input WP, inhibitors | Low-input whole proteome (TMT-exp) data from acutely and chronically stimulated human T cells with and without inhibitor treatment (related to Fig. 5 and Extended Data Fig. 8) |

## Data S3

**Table S3-1:** Polar metabolite abundance values in human T cells at day 2, day 4, or day 8 of culturing with or without chronic stimulation as determined by LC-MS/MS analysis. The channel ratio values after quantile regression imputation of left censored data (QRILC) and cell volume normalization are shown alongside raw signal intensity values and differential expression calculations. Data are from n = 4 donors. **Table S3-2:** Ratios of nucleoside to nucleotide, nucleoside triphosphate to diphosphate, and oxidized to reduced glutathione in human T cells activated for 2 days (D2) or cultured for 8 days with (D8C) or without (D8A) chronic stimulation. Each ratio is normalized to its D2 value within a donor, so every D2 value is 1 by construction. P-values are from a Friedman test with Dunn’s multiple comparisons test (adjusted p-values), computed in GraphPad Prism 10.3.0. GTP:GDP is from n = 3 donors, every other ratio from n = 4. **Table S3-3:** Abundance of stable metabolite isotopologues in D2, D8A, or D8C human T cells cultured in the presence of universally ^13^C-labeled glucose or glutamine. **Table S3-4:** Abundance of stable metabolite isotopologues in D8A, or D8C human T cells cultured in the presence of universally ^13^C-labeled glucose or glutamine, used for processivity analysis. **Table S3-5:** Lipid abundance values in human T cells at day 2, day 4, or day 8 of culturing with or without chronic stimulation. The channel ratio values after quantile regression imputation of left censored data (QRILC) and cell volume normalization are shown alongside raw signal intensity values and differential expression calculations. Data are from n = 2 donors.

| # | Title | Description |
| --- | --- | --- |
| 1 | S3-1 Polar metabolites | Abundance of polar metabolites in activated, acutely, and chronically stimulated human T cells (related to Figures 2 and Extended Data Fig. 3) |
| 2 | S3-2 Metabolite ratios | Nucleotide and glutathione ratios in activated, acutely, and chronically stimulated T cells |
| 3 | S3-3 Isotope tracing | Abundance of stable metabolite isotopologues in D2, D8A, or D8C human T cells cultured in the presence of universally <sup>13</sup>C-labeled glucose or glutamine (related to Extended Data Fig. 6) |
| 4 | S3-4 Additional isotope tracing | Abundance of stable metabolite isotopologues in D8A or D8C human T cells cultured in the presence of universally <sup>13</sup>C-labeled glucose or glutamine, used for processivity analysis (related to Extended Data Fig. 6) |
| 5 | S3-5 Whole cell lipidomics | Abundance of lipids in activated, acutely, and chronically stimulated human T cells (related to Extended Data Fig. 6) |

## (B) Biological Methods

## EXPERIMENTAL MODEL AND SUBJECT DETAILS

### Human samples

All studies involving primary human T cells used in *in vitro* experiments were conducted using buffy coats obtained from the New York Blood Center, and leukopaks or LRS cones from STEMCELL Technologies. The primary human T cells were isolated from blood samples collected from randomly selected, de-identified healthy donors. Donor sex was not recorded. Human tumor tissue samples were processed as previously described ^1^. Briefly, samples were obtained from patients with pathologically confirmed kidney cancer who had consented to MSKCC’s institutional biobanking protocol (MSKCC IRB #06-107). Utilization of biospecimens was approved under IRB #12-237 and IRB #20-025. Samples were obtained from patients undergoing clinically indicated operative resections of their primary kidney tumor. Samples were directly obtained from the operating room during nephrectomy. Tumor tissue was transported from the operating room to the laboratory in sterile saline on ice and kept at 4 °C until processed.

### Isolation of human peripheral blood mononuclear cells (PBMCs) and T cells

For isolation of human PBMCs, 12.5 mL of Lymphoprep density gradient medium was added to 50 mL Falcon tubes, followed by careful overlaying of 30 mL of whole blood diluted 2-fold in phosphate-buffered saline (PBS). Tubes were centrifuged for 20 min at 524 x *g* at room temperature (18-22 °C) with the brake off and acceleration set to 5. Following centrifugation, buffy coats containing white PBMC layer were collected, pooled into sterile Falcon tubes, washed with 15 mL PBS, and centrifuged for 8 min at 520 x *g* (4 °C) with the brake on. Pellets were subsequently combined and resuspended in 5 mL hypotonic red blood cell (RBC) lysis buffer (cat. sc-296258) for 5 min. After this time, tubes were topped up to 50 mL with PBS, centrifuged again, and cells were resuspended in 20 mL resuspension buffer (PBS without calcium or magnesium, supplemented with 2% FBS and 2 mM EDTA). Cell count and viability were assessed using a hemocytometer with Trypan blue staining. T cells were isolated using negative selection EasySep human T cell isolation kits (STEMCELL Technologies) according to slightly modified manufacturer’s instructions. Briefly, PBMCs were diluted to 50 million cells per mL in resuspension buffer. The resuspended cells were placed into 14 mL separation tubes (maximum 8 mL/tube), followed by the addition of isolation cocktail (25 µL/mL) and incubation for 5 min at room temperature. A suspension of RapidSpheres (25 µL/mL) was added, mixed by pipetting up and down, and incubated for another 5 min. The cell suspension was then topped up to 10 mL with resuspension buffer and incubated for 5 min on a magnetic rack to remove bead-bound cells. The supernatant containing T cell-enriched cell suspension was transferred into a new 14 mL separation tube on the magnetic rack, followed by incubation for 5 min to completely remove bead-bound cells. The supernatant was then transferred into a 50 mL conical tube, after which cells were centrifuged and resuspended in T cell media (RPMI-1640 supplemented with 10% heat-inactivated FBS, 2 mM *L*-glutamine, and 100 U/mL Penicillin-Streptomycin).

### Isolation of Tumor infiltrating lymphocytes (TILs) from human renal cell carcinoma tumors

Tumor infiltrating lymphocytes (TILs) from human tumors were isolated as described before ^1^. Briefly, single-cell suspensions from human renal cell carcinoma tumors were generated using Human Tumor Dissociation Kit (Miltenyi Biotec, 130-095-929), following manufacturer’s instructions. In brief, tumors were cut into small pieces (∼2-4 mm³) and transferred to gentleMACS C tubes (Miltenyi Biotec, 130-093-237) with R-10 medium (RPMI-1640 medium supplemented with 10% heat inactivated FBS, 4 mM *L-*glutamine, 100 U/mL Penicillin-Streptomycin and 10 mM HEPES). Based on the tumor weight, volume of R-10 medium was adjusted and enzyme mix was added. Samples were dissociated with a gentleMACS Octo Dissociator with Heaters (Miltenyi Biotec, 130-134-029) using the 37C_h_TDK3 program for 30 min. Dissociated tissue was then mashed with the flat end of a syringe plunger, passed through a 100 μm strainer, washed with 5 mL of R-10 medium and centrifuged at 500 x *g* for 5 min at 4 °C. Single-cell suspension was resuspended in 1 mL ACK buffer and incubated at RT for 3 min to lyse erythrocytes. Cells were then washed with 9 volumes of R-10 medium, centrifuged at 500 x g for 5 min at 4 °C, and either cryopreserved in Bambanker freezing medium (Fisher Scientific, NC3072290) or stained for sorting. Cells were stained for viability with 1:400 Ghost Dye Violet (Cell Signaling Technology, 59863S) and 1:20 Human TruStain FcX Fc Receptor Blocking Solution (BioLegend, 422302) in 1X PBS for 10 min on ice. Thereafter, cells were washed with 1X PBS, centrifuged at 350 x *g* for 5 min at 4 °C and stained for surface markers described below with 1X BD Horizon Brilliant Stain Buffer (BD Biosciences, 566349) in FACS buffer (2% FBS in 1 X PBS without calcium and magnesium) for 30 min at 4 °C. Cells were then washed with FACS buffer, spun down at 350 x *g* for 5 min at 4 °C, resuspended in FACS buffer, filtered through a 35 µm nylon mesh (Corning, 352235) and sorted on a SH800S Sony Cell Sorter into tubes containing FACS buffer. Sorted TILs (CD3^+^ T cells) cells were centrifuged at 520 x *g* for 5 min at 4 °C, washed once with ice cold 1X PBS, and cell pellets were snap frozen on dry ice until further processing for proteomic and Western blot analyses.

#### Human TILs sort panel

| Staining type | Marker | Fluorophore | Catalog # | Clone | Dilution |
| --- | --- | --- | --- | --- | --- |
| Surface | CD3 | BV570 | 300436 | UCHT1 | 1:25 |
| Surface | CD56 | PE-Cy7 | 362510 | 5.1H11 | 1:50 |
| Surface | CD11b | PE-Cy7 | 101216 | M1/70 | 1:400 |
| Surface | CD19 | PE-Cy7 | 363012 | SJ25C1 | 1:100 |
| Surface | CD45 | BV421 | 368522 | 2D1 | 1:200 |
| Surface | PD1 | APC | 329908 | EH12.2H7 | 1:100 |

### Mouse T cell isolation

Experiments involving primary mouse cells were performed according to Memorial Sloan Kettering Cancer Center Institutional Animal Care and Use Committee (IACUC) guidelines (protocol number 20-10-012). Mouse T cells were isolated from spleens of 6-12-week-old C57BL/6J mice (Jackson Laboratory 000664). Briefly, mice were euthanized with CO_2_ followed by cervical dislocation. Spleens were collected in ice-cold mouse T cell media - RPMI-1640 medium (Media Preparation Core, MSKCC) supplemented with 10% heat inactivated FBS (GeminiBio, 100-106), 4 mM *L-*glutamine, 50 μM β-Mercaptoethanol (ThermoFisher, 21985023), 100 U/mL Penicillin-Streptomycin (ThermoFisher, 15140163). Single cell suspensions of splenocytes were generated by mashing spleens in 5 mL RPMI-1640 medium (Media Preparation Core, MSKCC) with the flat end of a syringe plunger and passing through a 40 µm strainer. Splenocytes were centrifuged at 1,200 rpm for 5 min at 4 °C, resuspended in 5 mL ACK buffer (150 mM NH_4_Cl, 10 mM KHCO_3_, 0.1 mM EDTA-Na_2_) and incubated at RT for 90 s to lyse erythrocytes. Cells were filtered through a 40 µm strainer and washed with 45 mL PBS to remove ACK buffer. Splenocytes were resuspended at 1 x 10^8^ cells/mL in isolation buffer (1 X PBS without calcium and magnesium containing 2% FBS and 2 mM EDTA) and used for T cell isolation using Dynabeads Untouched Mouse T cell kit (ThermoFisher, 11413D) according to the manufacturer’s protocol. Briefly, 500 µL of splenocyte suspension (5 x 10^7^ cells) was mixed with 100 µL heat inactivated FBS and 100 µL Antibody Mix and incubated for 20 min at 4 °C. Cells were washed with 10 mL isolation buffer to remove excess of antibody and centrifuged at 1,200 rpm for 5 min at 4 °C. Cells were resuspended in 4 mL isolation buffer, mixed with 1 mL pre-washed Mouse Depletion Dynabeads and incubated for 15 min at RT with gentle tilting and rotation. Thereafter, 5 mL of isolation buffer were added to the cell suspension and tubes were placed in magnets for 2 min. Untouched T cells present in the supernatant were transferred to new tubes, centrifuged at 1,200 rpm for 5 min at 4 °C, resuspended at 1 x 10^6^ cells/mL in mouse T cell media supplemented with 10 ng/mL mIL-2 (Peprotech, 212-12), and activated as described in “Generation of acutely or chronically stimulated mouse T cells”.

### Mouse tumor model and isolation of tumor infiltrating lymphocytes (TILs)

*In vivo* experiments involving mouse models were performed according to Memorial Sloan Kettering Cancer Center IACUC guidelines (protocol number 20-10-012). Parental 2838c3 KPCY (Kras^LSL-G12D/+^;Trp53^LSL-R172H/+^;Pdx1-Cre;Rosa26^YFP/YFP^) cell line was a kind gift of Dr. Katelyn Byrne ^2^. For all experiments presented here, the KPC cell line was generated by CRISPR Cas9-mediated deletion of YFP (sgRNA: 5’-GTAGCCGAAGGTGGTCACGA-3’) using the Lenti-CRISPRv2 plasmid ^3^ (Addgene, 52961) and further selected by fluorescence-activated cell sorting (FACS). KPC cells were implanted subcutaneously (2 × 10^5^ cells/mouse in 100 μL 1 X PBS) into the right flank of 8-week-old female C57BL/6J mice (Jackson Laboratory, 000664). When tumors reached an average of 1,500 mm^3^, mice were euthanized with CO_2_ followed by cervical dislocation, and tumors were harvested to isolate tumor infiltrating lymphocytes (TILs). Additionally, spleens were harvested to isolate T cells not exposed to persistent tumor antigen stimulation. When collected, spleens and tumors were placed in cold RPMI+++ media - RPMI-1640 medium (Media Preparation Core, MSKCC) supplemented with 10% heat inactivated FBS (GeminiBio, 100-106), 4 mM *L-*glutamine, and 100 U/mL Penicillin-Streptomycin (ThermoFisher, 15140163) - until further processing. Spleens and tumors were then cut in small pieces and incubated for 45 min at 37 °C, 250 rpm shaking, in 1-2 mL of digestion buffer containing RPMI-1640 medium supplemented with 100 X DNase I (Millipore Sigma, 10104159001), 350 units/mL hyaluronidase (Worthington Biochemical Corporation, LS002592) and 50 μg/mL Liberase (Millipore Sigma, 05401020001). Single cell suspensions of tumors and spleens were generated by mashing tissues in RPMI+++ media with the flat end of a syringe plunger and passing through a 100 µm strainer (Greiner Bio-One, 542000). Cells were centrifuged at 500 x *g* for 5 min at 4 °C, resuspended in 1-2 mL ACK buffer and incubated at RT for 5 min to lyse erythrocytes. Cells were then washed with 9 volumes of 1 X PBS to quench ACK buffer, centrifuged at 500 x *g* for 5 min at 4 °C, and proceeded with Dynabeads FlowComp Mouse Pan T (CD90.2) Kit (FisherScientific, 11465D) following manufacturer’s instruction to enrich for T cell population. After enrichment, cells were stained on ice for viability with 1:2000 Ghost Dye Red 780 (Cytek Biosciences, 13-0865-T100) and 1:100 TruStain FcX (BioLegend, 101320) in 1X PBS for 10 min. Following centrifugation at 300 x *g* for 5 min at 4 °C, cells were stained for surface markers described below with 1X BD Horizon Brilliant Stain Buffer (BD Biosciences, 566349) in FACS buffer for 30 min at 4 °C, then washed with FACS buffer and spun down at 300 x *g* for 5 min at 4 °C. Cells were then resuspended in FACS buffer and sorted on a BD FACSymphony S6 Cell Sorter into tubes containing FACS buffer. Sorted cells were centrifuged at 520 x *g* for 5 min at 4 °C, washed once with ice cold 1X PBS, and cell pellets were snap frozen on dry ice until further processing for proteomic and Western blot analyses.

#### Spleen and TILs sort panel

| Staining type | Marker | Fluorophore | Catalog # | Clone | Dilution |
| --- | --- | --- | --- | --- | --- |
| Surface | CD45 | BV570 | 103136 | 30-F11 | 1:800 |
| Surface | CD11b | BUV395 | 563553 | M1/70 | 1:800 |
| Surface | NK1.1 | BUV395 | 564144 | PK136 | 1:200 |
| Surface | CD45R/<br>B220 | BUV395 | 563793 | RA3-6B2 | 1:800 |
| Surface | TCRb | BUV805 | 748405 | H57-597 | 1:100 |
| Surface | CD8 | BUV615 | 613004 | 53-6.7 | 1:800 |
| Surface | CD4 | PE | 100408 | GK1.5 | 1:800 |
| Surface | CD44 | FITC | 11-0441-85 | IM7 | 1:400 |
| Surface | PD1 | PE-Cy7 | 109110 | RMP1-30 | 1:200 |

### Lymphocytic choriomeningitis virus (LCMV) infection and isolation of virus reactive mouse T cells

Lymphocytic choriomeningitis virus (LCMV) was obtained from the Gene Transfer, Targeting, and Therapeutics Core, Salk Institute for Biological Studies, La Jolla, CA, USA. C57BL/6J female mice (10-week-old) were infected with LCMV Armstrong (2 × 10^5^ PFU/mouse in 100 μL 1 X PBS, via intraperitoneal injection) or Clone 13 (2 × 10^6^ PFU/mouse in 100 μL 1 X PBS, via retro-orbital injection). Spleens from LCMV-infected mice were harvested on day 8 (Armstrong and Clone 13) and day 15 (Clone 13) post-infection. At experiment endpoint, mice were euthanized with CO_2_ followed by cervical dislocation, and spleens were collected to isolate virus-reactive T cells. When collected, spleens were placed in cold RPMI+++ media until further processing. Single cell suspensions of spleens were generated by mashing tissues in RPMI+++ media with the flat end of a syringe plunger and passing through a 100 μm strainer. Cells were centrifuged at 500 x *g* for 5 min at 4 °C, resuspended in 1 mL ACK buffer and incubated at RT for 3 min to lyse erythrocytes. Cells were then washed with 9 volumes of 1 X PBS to remove ACK buffer, centrifuged at 500 x *g* for 5 min at 4 °C, and proceeded with Dynabeads Untouched Mouse CD8 Cells Kit (ThermoFisher, 11417D) following manufacturer’s instruction to enrich for CD8^+^ T cells. After enrichment, cells were stained on ice for viability staining with 1:2000 Ghost Dye Red 780 (Cytek Biosciences, 13-0865-T100) and 1:100 TruStain FcX (BioLegend, 101320) in 1X PBS for 10 min. Following centrifugation at 300 x *g* for 5 min at 4 °C, cells were stained for surface markers described below with 1X BD Horizon Brilliant Stain Buffer (BD Biosciences, 566349) in FACS buffer for 30 min at 4 °C, then washed with FACS buffer and spun down at 300 x *g* for 5 min at 4 °C. Cells were then resuspended in FACS buffer and sorted on a BD FACSAria III Cell Sorter into tubes containing FACS buffer. Sorted cells were centrifuged at 520 x *g* for 5 min at 4 °C, washed once with ice cold 1X PBS, and cell pellets were snap frozen on dry ice until further processing for proteomic and Western blot analyses.

#### LCMV T cell sort panel

| Staining type | Marker | Fluorophore | Catalog # | Clone | Dilution |
| --- | --- | --- | --- | --- | --- |
| Surface | TCRb | PerCP/Cy5.5 | 109228 | H57-597 | 1:100 |
| Surface | CD8 | APC | 100712 | 53-6.7 | 1:800 |
| Surface | CD4 | BV711 | 100550 | GK1.5 | 1:800 |
| Surface | CD44 | PE | 12-0441-83 | IM7 | 1:400 |
| Surface | PD1 | PE-Cy7 | 109110 | RMP1-30 | 1:200 |

## METHODS DETAILS

### T cell activation (D2 activated T cells)

For T cell activation, untreated 15 cm culture dishes were pre-coated with 17 mL of a solution of anti-CD3 antibody (5 µg/mL, diluted 1:200 from a 1 mg/mL stock; original stock concentration: 6.66 mg/mL) and anti-CD28 antibody (2 µg/mL; stock concentration: 7.3 mg/mL) in PBS. Plates were incubated overnight at 4 °C or, alternatively, for 2 h at 37 °C. Prior to cell plating, the coating solution was aspirated, and plates were washed twice with 5 mL of PBS. Primary human T cells, isolated by negative selection as described above, were resuspended in T cell media supplemented with IL-7 and IL-15 at final concentrations of 5 ng/mL each (5 µL of each stock per 100 mL of media). On day 0, T cells were plated at a density of 1 × 10⁶ cells/mL (20 × 10⁶ cells per 15 cm plate) and incubated for 48 h at 37 °C and 5% CO_2_ for activation before being allocated to conditions with and without chronic stimulation.To prepare cytokines for T cell culture *in vitro*, human recombinant IL-7 and IL-15 (STEMCELL Technologies) were reconstituted at 100 µg/mL by dissolving 100 µg of lyophilized powder in 1 mL of 0.1% BSA in molecular-grade water. Aliquoted stock solutions were stored at −80 °C until use.

### Generation of acutely and chronically stimulated primary human T cells (D4A, D8A, D15A, D4C, D8C, D15C)

On day 2, activated T cells were harvested by pipetting up and down multiple times to ensure complete detachment from the plate. Cells were spun down (520 x *g*, 5 min, 4 °C), counted using trypan blue exclusion, and resuspended in fresh T cell media containing IL-7 and IL-15 at a density of 1 × 10⁶ cells/mL. Cells (20 × 10⁶) were plated on 15 cm dishes, which have either been pre-coated with anti-CD3 antibody (chronic stimulation; 5 µg/mL in PBS, 200x stock) or used without pre-coating (acute stimulation). Every 48 hours, all cells from each respective condition were pooled, counted, and resuspended in fresh media containing IL-7 and IL-15 at 1 × 10⁶ cells/mL. Cells were replated at 20 × 10⁶ cells per 15 cm dish, either on anti-CD3 pre-coated plates or uncoated plates, following the same conditions as day 2. This subculture procedure was repeated every two days to maintain consistent experimental conditions with the exception of the last timepoint, which was collected one day after re-plating (D15A/D15C).

### Generation of acutely or chronically stimulated mouse T cells

Generation of acutely or chronically stimulated mouse T cells was performed as previously described ^4^. Briefly, freshly isolated T cells were seeded at 1 x 10^6^ cells/mL and activated with plate-bound anti-CD3 (3 μg/mL, ThermoFisher, 16-0031-38) and anti-CD28 (1 μg/mL, ThermoFisher, 16-0281-85) for 48 h at 37 °C and 5% CO_2_ in mouse T cell media supplemented with 10 ng/mL mIL-2 (Peprotech, 212-12), henceforth referred to as ‘complete mouse T cell media’. On day 2, following 48 h of activation, activated T cells were harvested by pipetting up and down multiple times to ensure complete detachment from the plate. Cells were spun down (300 x *g*, 5 min, 4 °C), counted using a Beckmann Coulter Multisizer 3 Particle Size Analyzer with a gate of between 200 and 1000 femtoliters (fL), and resuspended in fresh complete mouse T cell media at a density of 1 × 10⁶ cells/mL. Cells (2 × 10⁶) were plated in 6-well plates, which had either been pre-coated with anti-CD3 antibody (3 μg/mL in PBS, ThermoFisher, 16-0031-38) or used without pre-coating (acute stimulation). Every 48 h, all cells from each respective condition were pooled, counted, and resuspended in fresh complete mouse T cell media at 1 × 10⁶ cells/mL. Cells were replated at 2 × 10⁶ cells per well of a 6-well plate, either on anti-CD3 pre-coated plates or uncoated plates, following the same conditions as day 2. This subculture procedure was repeated every two days to maintain consistent experimental conditions. Cell density was kept below 5 × 10^6^ cells/mL on each passage.

### Flow cytometry analysis

#### Inhibitory receptor panel

Cells were plated at 200,000 live cells per well in a U-bottom 96-well plate. Cells were spun at 520 x *g* for 3 min at 4 °C and washed with 1% BSA in 1X PBS. Cells were incubated with Fc block (cat. 422302) at a 1:20 dilution for 30 min at 4 °C. Following centrifugation, cells were stained for surface markers with 1X Brilliant Buffer (cat. 566349) in 1% BSA in PBS for 30 min at 4 °C, then washed with 1% BSA in PBS and spun down. Viability staining was performed for 30 min at 4 °C using Ghost Dye 510 Violet (cat. 59863) diluted 1:800 in PBS. Following washing and centrifugation, cells were fixed for 30 min at room temperature using the eBioscience Intracellular Fixation & Permeabilization Buffer Set (cat. 88-8824-00). Fixed cells were centrifuged at 700 x *g* for 5 min and washed twice with permeabilization buffer. Cells were stained overnight at 4 °C for two of the intracellular markers, T-Bet and TCF-1, diluted in permeabilization buffer. The following day, cells were spun down and resuspended in stains for two additional markers (CTLA4 and TOX) for a further 30 min at 4 °C. Cells were washed twice in permeabilization buffer and resuspended in 1% BSA in PBS. Data was acquired using a Cytek Aurora spectral cytometer, with a minimum of 5,000 live CD8 cells measured per condition. Gating for CD4 and CD8 was determined by eye, while all other gates were set using fluorescence minus one (FMO) controls.

| Staining type | Marker | Fluorophore | Catalog # | Clone | Dilution |
| --- | --- | --- | --- | --- | --- |
| Surface | CD4 | BUV395 | 363-0047-42 | SK3 | 1:200 |
| Surface | CD8 | BUV496 | 612942 | RPA-T8 | 1:800 |
| Surface | CD38 | BUV737 | 612825 | HB7 | 1:400 |
| Surface | CD39 | PE-Fire 810 | 328245 | A1 | 1:200 |
| Surface | HLA-DR | BV786 | 564041 | G46-6 | 1:200 |
| Surface | PD-1 | APC | 329908 | EH12.2H7 | 1:100 |
| Surface | LAG3 | BV421 | 369314 | 11C3C65 | 1:100 |
| Surface | TIGIT | BUV615 | 570444 | TgMab-2 | 1:50 |
| Intracellular (O/N) | T-Bet | R718 | 568175 | 4B10 | 1:200 |
| Intracellular (O/N) | TCF-1 | Alexa 488 | 6444S | C63D9 | 1:3,200 |
| Intracellular (30 m) | CTLA4 | BV711 | 369632 | BNI3 | 1:400 |
| Intracellular (30 m) | TOX | PE | 130-120-716 | REA473 | 1:50 |

#### Cytokine panel

Experiments were performed as above with the following differences: Upon cell counting and plating, cells were resuspended in T cell media containing phorbol 12-myristate 13-acetate (PMA, cat. P1585, 50 ng/mL) and ionomycin (cat. I0634, 1 µg/mL). After 1 h, Brefeldin A (cat. 00-4506-51, 1:1,000 dilution) was added to the media for an additional 3 h incubation at 37 °C. Blocking, surface and viability staining was performed as for the inhibitory receptor panel. Fixation was performed using the BD Biosciences Cytofix/Cytoperm kit (cat. 554714) for 30 min at 4 °C. All intracellular staining was performed overnight.

| Staining type | Marker | Fluorophore | Catalog # | Clone | Dilution |
| --- | --- | --- | --- | --- | --- |
| Surface | CD4 | BUV395 | 363-0047-42 | SK3 | 1:200 |
| Surface | CD8 | BUV496 | 612942 | RPA-T8 | 1:800 |
| Surface | PD-1 | APC | 329908 | EH12.2H7 | 1:100 |
| Intracellular | GzmB | PerCP-Cy5.5 | 372212 | QA16A02 | 1:200 |
| Intracellular | IFNγ | FITC | 502506 | 4S.B3 | 1:200 |
| Intracellular | IL-2 | APC-Fire 750 | 500352 | MQ1-17H12 | 1:400 |
| Intracellular | TNF | PE | 502909 | MAb11 | 1:1,600 |

#### Restimulation flow on inhibitor-treated samples

To evaluate function under more physiologically relevant restimulation conditions, we used ImmunoCult Human CD3/CD28/CD2 T Cell Activator beads (cat. 10970) at a concentration of 12.5 µL/mL. After 1 hour, Brefeldin A (cat. 00-4506-51, 1:1,000 dilution) was added to the media for an additional 5 h incubation at 37 °C. All other steps were performed as for the “Cytokine panel” experiments.

#### TCR transduction confirmation

To confirm successful transduction of primary human T cells with the 1G4 TCR, single-cell suspensions were first stained with Ghost Dye 510 (see Key Resources Table) and Human FcBlock (Human TruStain FcX) in a volume of 50 µL of PBS for 10 min in the dark at room temperature (RT). Cells were then washed with 200 µL of FACS buffer and centrifuged at 700 x *g* for 2 min at 4 °C. Cells were then resuspended in 50 µL of surface stain containing 1x Brilliant Buffer in FACS buffer and the below panel for 20 min at RT in the dark. Following surface staining, cells were washed once with FACS buffer and resuspended in a final volume of 200 µL. Immediately prior to acquisition, cell suspensions were passed through a 100 µm nylon mesh filter to ensure a single-cell suspension. Data were acquired on a Cytek Aurora. An untransduced culture from the same donor was run simultaneously to gate for the true 1G4 TCR (identified by Mouse TCR β Chain) positive population and this percentage was used as the transduction efficiency to plate the Incucyte-based tumor cell killing assays.

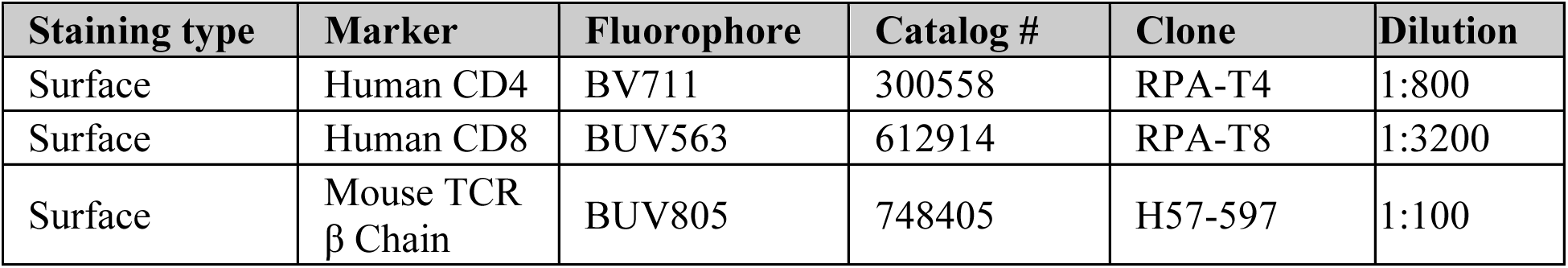

### Oxidative stress assays

To assess oxidative stress in primary human and mouse T cells, we used dihydroethidium (DHE) and MitoSOX Red (MSR) as indicators of superoxide accumulation. Stock solutions were freshly prepared as follows: DHE was reconstituted at 5 mM in DMSO, Antimycin A (AA) at 10 mM in ethanol, and MSR at 1 mM in DMSO. Working solutions were prepared in PBS containing 5 mM pyruvate, with final concentrations of 0.625 µM DHE, 1.64 µM MSR, and 10 µM AA.

Cells were seeded at 2 × 10⁵ cells per well in a 96-well U-bottom plate and centrifuged at 520 × g for 5 min. After washing with PBS, cells were incubated with a blocking buffer containing an Fc receptor blocker (cat. 422302, 1:20) and Ghost Dye Violet 510 (cat. 59863S, 1:800) for 10 min at room temperature. Following another PBS wash, surface staining was performed by incubating cells with BUV395-conjugated CD8 antibody (1:200) along with superoxide detection dyes (DHE or MSR) in the presence or absence of AA for 30 min at 37 °C in a humidified incubator, protected from light. Staining conditions included unstained controls, single-color controls, fluorescence-minus-one (FMO) controls, and a positive control with AA treatment. After surface staining and oxidative stress dye incubation, cells were washed with ice-cold PBS, resuspended in 200 µL ice-cold FACS buffer, kept on ice, preventing cells from light until analysis. Flow cytometry was performed using a Cytek Aurora, with acquisition gated on CD8⁺ T cells and a stopping gate of 1,500 events per sample. Data were analyzed using FlowJo.

### Extracellular flux analysis

Oxygen consumption rate (OCR), extracellular acidification rate (ECAR), and ATP production rates were measured using a Seahorse XFe96 Extracellular Flux Analyzer (Agilent). Primary human T cells were isolated, activated, and cultured as described above. Activated (D2), acutely, and chronically stimulated T cells (D8A, D8C) from the same donor were analyzed on the same plate. To enable this, freshly isolated T cells were cryopreserved in Bambanker freezing medium at −80 °C until day 6 of cell culture. At that time, the cells were thawed, spun down to remove the freezing medium (92 x *g*, 10 min), re-suspended in T cell culture medium, and activated for 2 days, allowing D2 cells to be analyzed in parallel with D8A and D8C cells from the same donor. One day before the analysis, XFe96/XF Pro cell culture microplates (Agilent, 103794-100) were coated with 20 µL of 50 µg/mL poly-*L-*lysine (Millipore Sigma, P4707) in 1X PBS and incubated for 30 min at RT. Plates were washed twice with 100 µL of 1X PBS, air dried for 1-2 h and stored overnight at 4 °C. On the day of the analysis, T cells were spun down at 300 x *g* for 5 min at 4 °C and washed once with Seahorse medium (XF RPMI medium, 10 mM XF glucose, 1 mM XF pyruvate, and 2 mM *L-*glutamine). T cells were then plated at 1 x 10^5^ per well in Seahorse medium on XFe96/XF Pro poly-*L-*lysine-coated plates and centrifuged at 300 x *g* for 1 min at 4 °C. Cells were incubated in a non-CO_2_ incubator at 37 °C for 45-60 min before the analysis. OCR and ECAR were measured at basal level and after treatment with oligomycin (1 μM), FCCP (1 μM), and rotenone/antimycin mix (0.5 μM), contained in the Seahorse XF Cell Mito Stress Test Kit (Agilent, 103015-100). OCR, ECAR, and ATP production rates were determined using Agilent Seahorse Analytics XF software.

### Incucyte-based tumor cell killing assay^5^

#### *1G4-LY* TCR transfer plasmids

The HLA-A*02:01-restricted NY-ESO-1_157-165_-specific TCR 1G4 has been described previously^6^. Human constant regions of the 1G4 TCR were replaced with codon optimized murine constant regions^7^ harboring a second disulfide bond^8^ as previously described. Codon optimized sequences encoding a bicistronic 1G4 TCR construct were synthesized by Genscript and subcloned into the lentiviral transfer plasmid pZR071^9^ (Addgene, #180264) via restriction enzyme cloning using MluI-HF (NEB, #R3198S) and SbfI-HF (NEB, #R3642S). The affinity enhanced 1G4-LY ^6^ was generated via site directed mutagenesis (NEB, #E0554S). Final sequences were verified by whole plasmid sequencing (Plasmidsaurus).

#### Tumor cell lines

For repetitive cytotoxicity assays, A375 (ATCC, CRL-1619) and SK-MEL-37 (Antibody and Bioresource Core Facility at MSKCC and RU, CVCL_3878) melanoma cells, were retrovirally transduced with nuclear localized mCherry (pMSGV1_mCherry-NLS_P2A_blastR), selected with blasticidin (10 µg/mL) and either used as bulk population (SK-MEL-37) or single cell cloned by limiting dilution (A375). The NLS is derived from *c-*MYC (PAAKRVKLD) and attached to the *C-*terminus of mCherry. The P2A sequence is preceded by a furin cleavage site (RAKR). The change from puroR to blastR was done by Genscript.

#### Lentivirus production and concentration

Lentiviral production was performed according to previously published protocols, with minor modifications.^9, 10^ Lentiviral vectors were generated by co-transfecting Lenti-X 293T cells (Takara, cat #632180) with a third-generation packaging system. Briefly, Lenti-X 293T cells were maintained in DMEM supplemented with 10% FBS and seeded at a density of 3-3.5 x 10^6^ cells per 10 cm dish in cOPTIMEM (OPTIMEM supplemented with 5% FBS, 2 mM *L-*glutamine, 1x MEM non-essential amino acids (Gibco #11140-050), and 1 mM sodium pyruvate (Gibco #11360-070) 16-24 h prior to transfection to reach 70-90% confluency. Transfection was performed using Lipofectamine 3000 (Thermo Fisher Scientific) according to the manufacturer’s instructions. For a 10 cm dish, a total of 21.25 µg of DNA, comprising the transfer plasmid, psPAX.2 (Addgene #12260), and pMD2.G (Addgene #12259) at a 1:1:1 molar ratio, was mixed with P3000 reagent in Opti-MEM and added to the Lenti-X 293T cells. When other culture formats were used, reagent volumes and amounts were scaled proportionally by surface area. Three to six hours post-transfection, 1x Viral Boost (Alstem #VB100) was added.

Viral supernatants were harvested at 24 and 48 h post-transfection. The harvests were combined and cleared of debris by centrifugation (500 x *g* for 5 min) and filtered through a 0.45 μm PES filter. The virus was concentrated using Lentivirus Precipitation Solution (Alstem #VC100) at a 1:4 (v/v) ratio by incubating for a minimum of 3 h to overnight at 4 °C and centrifuged at 1,500 x *g* for 30 min. The resulting viral pellets were resuspended in cold RPMI at a 1/100 of the original volume and either used immediately or aliquoted and stored at −80 °C.

#### Primary human T cell transduction

Primary human bulk T cells (CD3^+^) were isolated from healthy donor PBMCs by negative selection and activated as described above. Twenty-four hours after initiating activation, T cells were transduced with concentrated lentivirus at a final volume of 2% (v/v) by pre-diluting the 100x viral stocks at 1:5 in T cell medium before addition to the culture. After 24 h of viral exposure, cells were centrifuged and resuspended in fresh RPMI+++ media supplemented with 5 ng/mL hIL-7 (Peprotech, 200-07) and hIL-15 (Peprotech, 200-15). Successful transduction was confirmed by flow cytometry.

#### Cell killing assay^5^

Tumor cells stably expressing a nuclear-localized fluorescent reporter (A375 mCherry-NLS or SK-MEL37 mCherry-NLS) were seeded into flat-bottom 96-well plates (Corning 3599) at 1 × 10⁴ cells per well in 100 µL RPMI supplemented with 10% fetal bovine serum and 1% penicillin–streptomycin (RPMIc). Cells were allowed to adhere for ≥3 h before baseline imaging. Plates were loaded onto an Incucyte live-cell imaging system and scanned using a 10× objective, acquiring phase contrast and red fluorescence signal with preset acquisition parameters. Four images per well were collected at 3-4 h intervals. Plates were equilibrated for at least 30 min prior to imaging to minimize condensation artifacts.

After baseline acquisition, plates were removed from the Incucyte and T cells were added in 100 µL RPMIc at the indicated effector-to-target (E:T) ratios, accounting for TCR transduction efficiency as determined by flow cytometry as described above on the day of assay setup. Conditions were plated in technical triplicates. Control wells included tumor cells alone and tumor cells co-cultured with untransduced T cells. Plates were reloaded into the Incucyte at least 30 min prior to imaging to minimize condensation artifacts.

Every 48 h thereafter, 75 µL of medium was carefully removed from each well without disturbing the cell layer, and fresh tumor cells (1 × 10⁴ per well in 100 µL RPMIc) were added. Longitudinal imaging was continued throughout the assay under identical acquisition settings.

### RNA sequencing of activated, acutely, and chronically stimulated human T cells

#### Sample preparation

Primary human T cells were activated and expanded with or without chronic stimulation as described above. Following 2, 4, and 8 days of activation, biological triplicate wells were harvested and RNA was extracted using a TRIzol-chloroform-based method^11^. After RiboGreen quantification and quality control by Agilent BioAnalyzer, 500 ng of total RNA with RIN values of 9.4-10 underwent polyA selection and TruSeq library preparation according to instructions provided by Illumina (TruSeq Stranded mRNA LT Kit, catalog # RS-122-2102), with 8 cycles of PCR. Samples were barcoded and run on a NovaSeq 6000 generating 100-bp paired-end reads, using the NovaSeq S4 Reagent Kit (200 Cycles) (Illumina). An average of 127 million paired reads was generated per sample. Ribosomal reads represented 0.1-1.2% of the total reads generated and the percent of mRNA bases averaged 88%. RNA sequencing was performed at the Integrated Genomics Operation Core, MSKCC.

#### Data processing

FASTQ files were quality-checked using FastQC v0.12.0 and trimmed with Trim Galore v0.6.10 to remove adapter sequences and read ends with Phred scores below 15. STAR v2.7.0a was used to align the FASTQ files to the hg38 reference genome. SAM files outputted by STAR were sorted using the sort function from Samtools v1.19.2 and indexed using the index function to generate BAM index files. A gene expression count matrix was generated using featureCounts v2.0.6, quantifying only read pairs that overlapped exons. Downstream differential gene expression analysis was performed using DESeq2 v1.42.

#### Principal component analysis

Principal component analysis (PCA) was performed using *sci-kit learn*^12^ library for Python. The 5,000 protein-coding genes with the highest row-wise variance were used and normalized count values were log_2_ transformed before PCA. The first two principal components (PC1 and PC2) were plotted.

#### Gene set enrichment analysis

The bulk RNA-Seq dataset was filtered to genes with a sum of least 1,000 reads across 15 samples (14,391 genes) for GSEA. Genes were ranked based on log_2_(FC) and the permutation type was set to gene-set.

### Single-cell RNA sequencing of acutely and chronically stimulated primary human T cells (D15A, D15C)

#### Sample preparation

Primary human T cells were isolated and cultured as described above (‘Generation of acutely and chronically stimulated primary human T cells’). On day 15 following initial activation, acute and chronic T cells were collected and stained with a viability dye along with anti-CD4 (RPA-T4, BioLegend 300534, 1:100) and anti-CD8 (RPA-T8, BD Biosciences 612914, 1:1,200). Live CD4^+^ and CD8^+^ T cells from acute and chronic conditions were flow sorted and underwent single cell gene expression profiling using the 10x Genomics Next GEM Single Cell 3’ Gene Expression Kit, v3.1. Library preparation from cDNA was performed using the MAS-Seq Kit (PacBio) followed by Illumina sequencing.

#### MAS-ISO-seq data preprocessing

PacBio HiFi reads were first segmented using the *split* function in Skera (v1.3.0). Primers were then removed using lima v2.12.0. With Iso-Seq (v4.0.0), tags (UMI, cell barcode) were clipped from reads using the *tag* function with the design parameter set as *‘T-12U-16B’*, reads were refined using the *refine* function with the *‘--require-polya’* parameter, single cell barcodes were corrected with the *correct* function using the ‘*3M-februrary-2018-REVERSE_COMPLEMENTED.txt’* barcode inclusion list, and reads were deduplicated with the *groupdedup* function. Following this, reads were aligned to GrCh38/hg38 with the pbmm2 (v1.16.99) *align* function with the *‘--preset ISOSEQ*’ parameter. Transcripts were quantified for each sample utilizing IsoQuant (v3.3.1) with transcript annotations from GENCODE GrCh38v39 and the parameter *‘--data_type pacbio_ccs’*. Custom scripts were then used to remove reads with mapq < 5 and reads that were labeled *‘noninformative’* or *‘intergenic’* by IsoQuant. The remaining reads were assigned to genes, and read assignments were then converted into a cell barcode to gene counts matrix. Additionally, a library size for each cell was calculated as the number of mapped reads in each cell.

The counts matrix was loaded into a Seurat object using the *CreateSeuratObject* function in Seurat (v5.1.0). The counts was normalized by multiplying by 10,000/(cell library size) and then log transformed with the *log1p* function. Cells were initially filtered such that cells with ≤100 expressed genes and cells with ≤100 total counts were removed. Additionally, genes were filtered such that genes expressed in ≤10 cells were removed. Next, cells were filtered based on total counts excluding ribosomal and mitochondrial genes. Cells with less than 2^10.3^ to 2^10.75^ non-ribosomal/mitochondrial counts were removed, with the exact cutoff determined on a per sample basis by visualization of count density. Cells were then filtered such that those with greater than 9 to 12 percent mitochondrial content were removed on a per sample basis. Doublet Detection v4.2 was used to annotate doublets with the parameter *‘voter_thresh=0.5’* for the *predict* function. After running *FindVariableFeatures* and *ScaleData* (default parameters), PCA was conducted using *RunPCA* with variable features as the *‘features’* parameter, and a neighborhood graph was constructed with *FindNeighbors* with ‘*dims = 1:15’*. Leiden clusters were computed for each sample with *FindClusters* (1.5 resolution). Leiden clusters with ≥25% cells assigned as doublets were removed. The samples were then combined into a single object, and visualized with a UMAP created by running *FindVariableFeatures*, *ScaleData* (default parameters)*, RunPCA* (using variable features), *FindNeighbors* with ‘*dims = 1:15’, FindClusters* (1.0 resolution), and *RunUMAP* with ‘*dims = 1:15’*.

CD8^+^ T cell subtype annotations from a Pan-cancer T cell atlas^13^ were transferred onto the *in vitro* CD8^+^ T cell scRNA-seq samples by first determining anchors between the two datasets with the *FindTransferAnchors* function with parameters *‘reference = [atlas data], query = [in vitro data], dims=1:30, reference.reduction=”pca’’. TransferData* function was called to generate predicted cell types for the *in vitro* data from the anchors using cell type annotations from the atlas as the reference data.

### Metabolomic analysis

#### Sample preparation

For lipid and polar metabolomic analysis, activated, acutely, and chronically stimulated human T cells were harvested at the specified days following initial activation. Cells were washed with cold PBS twice, pelleted, and supernatant was removed. Cell pellets were snap frozen and stored at −80 °C until analysis. Lipid and polar metabolite analysis were performed at Weill Cornell Medicine Proteomics and Metabolomics Core Facility.

#### Lipid analysis

Lipids were extracted from samples as previously described^14^. The extract was dried down using a SpeedVac and then reconstituted using acetonitrile/isopropanol/water 65:30:5 prior to LC-MS/MS analysis. Chromatographic separation was performed on a Vanquish UHPLC system with a Cadenza CD-C18 3 µm packing column (Imtakt, 2.1 mm id x 150 mm) coupled to a Q Exactive Orbitrap mass spectrometer (Thermo Scientific) via an Ion Max ion source with a HESI II probe (Thermo Scientific). The mobile phase consisted of buffer A: 60% acetonitrile, 40% water, 10 mM ammonium formate with 0.1% formic acid and buffer B: 90% isopropanol, 10% acetonitrile, 10 mM ammonium formate with 0.1% formic acid. The LC gradient was as follows: 0-1.5 min, 32% buffer B; 1.5-4 min, 32-45% buffer B; 4-5 min, 45-52% buffer B; 5-8 min, 52-58% buffer B; 8-11 min, 58-66% buffer B; 11-14 min, 66-70% buffer B; 14-18 min, 70-75% buffer B; 21-25 min, isocratic 97% buffer B, 25-25.1 min 97-32% buffer B; followed by 5 min of re-equilibration of the column before the next run. The flow rate was 200 μL/min. A data-dependent mass spectrometric acquisition method was used for lipid identification. In this method, each MS survey scan was followed by up to 10 MS/MS scans performed on the most abundant ions. Data was acquired in both positive mode and negative mode. The following electrospray parameters were used: spray voltage 3.0 kV, heated capillary temperature 350 °C, HESI probe temperature 350 °C, sheath gas, 35 units; auxiliary gas, 10 units. For MS scans: resolution, 70,000 (at m/z 200); automatic gain control target, 3e6; maximum injection time, 200 ms; scan range, 250-1800 m/z. For MS/MS scans: resolution, 17,500 (at 200 m/z); automatic gain control target, 1e5 ions; maximum injection time, 75 ms; isolation window, 1 m/z; NCE, stepped 20, 30, and 40. LC-MS files were processed using MS-DIAL software^15^ for lipid identification and relative quantitation.

#### Polar metabolite analysis

The sample was extracted using pre-chilled 80% methanol (−80 °C). The extract was dried with a SpeedVac, and redissolved in HPLC grade water before it was applied to the hydrophilic interaction chromatography LC-MS. Metabolites were measured on a Q Exactive Orbitrap mass spectrometer (Thermo Scientific), which was coupled to a Vanquish UPLC system (Thermo Scientific) via an Ion Max ion source with a HESI II probe (Thermo Scientific). A Sequant ZIC-pHILIC column (2.1 mm i.d. × 150 mm, particle size of 5 µm, Millipore Sigma) was used for separation of metabolites. A 2.1 × 20 mm guard column with the same packing material was used for protection of the analytical column. Flow rate was set at 150 μL/min. Buffers consisted of 100% acetonitrile for mobile phase A, and 0.1% NH_4_OH/20 mM CH_3_COONH_4_ in water for mobile phase B. The chromatographic gradient ran from 85% to 30% A in 20 min followed by a wash with 30% A and re-equilibration at 85% A. The Q Exactive was operated in full scan, polarity-switching mode with the following parameters: the spray voltage 3.0 kV, the heated capillary temperature 300 °C, the HESI probe temperature 350 °C, the sheath gas flow 40 units, the auxiliary gas flow 15 units. MS data acquisition was performed in the m/z range of 70–1,000, with 70,000 resolution (at 200 m/z). The AGC target was 1e6 and the maximum injection time was 250 ms. The MS data was processed using XCalibur 4.1 (Thermo Scientific) to obtain the metabolite signal intensities. Identification required exact mass (within 5 ppm) and standard retention times.

#### Data processing and imputation

Within each biological replicate, metabolites were required to have at least one condition with an average signal intensity above 5,000 (174/203 metabolites passed). After filtering and log transformation, missing data was imputed through quantile regression imputation of left-censored data (QRILC) performed using imputeLCMD version 2.1. Channel ratios (signal intensity / sum of signal intensities per metabolite) were calculated and used for all analyses. Metabolite class annotations were curated through literature review.

#### Principal component analysis

Metabolite ratio values were log_2_ transformed and only metabolites with a channel ratio above 0 in all channels before imputation (156 metabolites) were used. The first and third principal components (PC1 and PC3) were plotted for polar metabolite data. Principal component analysis was performed with *scikit-learn* ^12^ version 1.4.2 for Python version 3.1.1. Loadings for plotting were selected by taking the 5 metabolites with highest and lowest loadings for both principal components.

### Isotope tracing experiments

#### Sample preparation

T cells that had either been activated for 2 days (D2) or were cultured with and without chronic stimulation until day 8 post-activation as described in “Generation of acutely and chronically stimulated primary human T cells” (D8A, D8C) were washed with PBS and re-suspended in RPMI-1640 without glucose or glutamine (Media Preparation Core, MSKCC) containing 10% dialyzed FBS (GeminiBio, 100-108), 100 U/mL Penicillin-Streptomycin (ThermoFisher, 15140163), 5 μM BME (ThermoFisher, 21985023), 5 ng/mL hIL-7 (Peprotech, 200-07), and 5 ng/mL hIL-15 (Peprotech, 200-15), to which either universal ^12^C-glucose (Millipore Sigma, G7021) or [U-^13^C] glucose (Cambridge Isotope Laboratories, CLM-1396-PK) to a final concentration of 10 mM and either ^12^C-L-glutamine (ThermoFisher, A2916801) or [U-^13^C] glutamine (Cambridge Isotope Laboratories, CLM-1822-H-PK) to a final concentration of 2 mM were added. The cells were plated at 1.5 x 10^6^ cell/mL in 1 mL of the respective medium and incubated at 37 °C for 4 h. Following this time, plates were placed on ice, cells were rapidly harvested, centrifuged at 300 x *g* for 3 min at 4 °C, and resuspended in 1 mL of ice-cold 80% LC-MS grade methanol (Fisher scientific, A456-4) for metabolite extraction, which was carried out at −80 °C overnight. Subsequently, samples were vortexed and centrifuged at 20,000 x *g* for 20 min to remove proteins. Supernatants were then dried in a vacuum evaporator (Genevac EZ-2 Elite) for 3 h.

#### Isotopologue analysis

Dried extracts were resuspended in 60 μL of 60% acetonitrile in water for hydrophilic interaction liquid chromatography (HILIC) by the Cell Metabolism Core within the Donald B. and Catherine C. Marron Cancer Metabolism Center at MSKCC. Samples were vortexed, incubated on ice for 20 min and clarified by centrifugation at 20,000 x g for 20 min at 4 °C. HILIC LC-MS analysis was performed on a 6545 Q-TOF mass spectrometer (Agilent Technologies) in both positive and negative ionization modes using columns, buffers and LC-MS parameters as described previously^4^. Targeted data analysis, isotopologue extraction, and natural isotope abundance correction were performed using MassHunter Profinder software v.10.0 (Agilent Technologies).

### Proteomic platforms: Abundance-based proteomics with TMT multiplexing

#### Sample preparation

Six 10-plex experiments with primary human T cells from different biological donors were processed. Five experimental conditions were used in duplicate for each donor, including D2, D4A, D4C, D8A, and D8C T cells. Cells were pelleted (600 x *g*, 5 min, 4 °C), washed once with PBS, transferred into 1.5 mL Eppendorf tubes, flash-frozen, and stored at −80 °C until further processing. On the first day of the mass spectrometry workflow, frozen cell pellets were thawed on ice for 10 min. Ice-cold PBS supplemented with cOmplete EDTA-free Protease Inhibitor Cocktail (1 tablet per 10 mL) was added, and cells were lysed by sonication (three cycles of 8 pulses at 60% output). Protein concentrations were then determined using the standard DC Protein Assay (Bio-Rad) and normalized to a final concentration of 1-2 mg/mL.

Abundance-based proteomics with TMT multiplexing was performed following previously published protocols^17^. Briefly, normalized proteome samples (100 µL) were transferred into new 1.5 mL low-binding Eppendorf tubes containing 48 mg of urea, and vortexed to fully dissolve the urea, resulting in a final concentration of 8 M. DTT was then added (5 µL of 200 mM stock; 10 mM final concentration), and the samples were incubated at 65 °C for 15 min. After cooling for 5 min at room temperature, iodoacetamide was added (5 µL of 400 mM stock; 20 mM final concentration), followed by incubation at 37 °C for 30 min. The tubes were then placed on ice, and 400 µL of water was added to each sample. Proteins were precipitated using a methanol/chloroform protocol: 500 µL of cold methanol and 150 µL of chloroform were added to each tube, followed by centrifugation at 10,000 x *g* for 10 min to pellet the proteins. The supernatant was carefully removed without disturbing the protein pellet. If the protein pellet appeared small, the lower chloroform layer could be left in place until after the second methanol wash. An additional 400 µL of cold methanol was added to each sample, and the tubes were briefly sonicated to break up the protein disks. Samples were centrifuged again (10,000 x *g*, 10 min), and the resulting pellets were re-suspended with sonication in 160 µL of 200 mM EPPS buffer (pH 8.0, no urea). Finally, 6 µL of Trypsin/LysC solution (0.42 µg/µL in buffer containing 15 mM CaCl₂) was added to each sample. Digestion was carried out overnight at 37 °C with shaking.

Following the overnight digestion, peptide concentration was determined using the microBCA assay (Thermo Fisher) following manufacturer’s instructions. For each TMT labeling reaction, 25 µg of peptides was used, and the volume was adjusted to 35 µL with 200 mM EPPS buffer. Acetonitrile was then added (9 µL per sample), followed by the addition of the appropriate TMT tag (5 µL per sample; 20 µg/µL in anhydrous acetonitrile), resulting in a final acetonitrile concentration of approximately 25%. Samples were incubated at room temperature for 1 hour to allow labeling. Unreacted tags were quenched by adding 5 µL of 5% hydroxylamine in water and incubating for 15 min. Finally, the samples were acidified with 2.5 µL of formic acid per sample.

A ratio check (RC) sample was prepared by pooling 2 µL from each labeled sample. The combined sample was dried using a SpeedVac vacuum concentrator and desalted using C18 stage tips (10 µL, Thermo Fisher). Briefly, stage tips were activated and equilibrated with two washes of 20 µL acetonitrile, followed by three washes with 20 µL of buffer A (0.1% formic acid, 5% acetonitrile, 95% water). The dried peptide mixture was reconstituted in 20 µL of buffer A with sonication, then loaded onto the stage tip by centrifugation (1,000 x *g*, 1 min). After washing with buffer A (3 × 20 µL, 1,000 x *g*, 1 min), peptides were eluted with 20 µL of buffer B (0.1% formic acid, 20% acetonitrile, 80% water) and dried again using a SpeedVac.

The RC sample was reconstituted in 10 µL of buffer A and analyzed via liquid chromatography-mass spectrometry (LC-MS) using a 2 µL injection on an Orbitrap Eclipse mass spectrometer. The following 70-min LC gradient was used: 0-10 min: 5% buffer B (0.1% FA in acetonitrile) in buffer A (0.1% FA in water); 10-50 min: linear increase from 5% to 20% buffer B; 50-55 min: ramp to 45% buffer B; 55-57 min: ramp to 95% buffer B; 57-59 min: hold at 95% buffer B; 59-61 min: decrease to 5% buffer B; 61-63 min: ramp back to 95% buffer B; 63-70 min: re-equilibration at 5% buffer B. The RC sample was used to calculate normalization factors for each TMT channel. For a 10-plex experiment, a volume equivalent to N × 20 µL (where N is the normalization factor derived from RC) was taken from each channel and combined into a single tube. The pooled sample was dried overnight using a SpeedVac and desalted using a SepPak C18 column (50 mg). The column was first activated with 2 × 1 mL of acetonitrile, then equilibrated with 3 × 1 mL of buffer A. The dried peptide mixture was reconstituted in 1 mL of buffer

A, loaded onto the column, washed with buffer A, and eluted with buffer B into a clean Eppendorf tube. The eluate was dried overnight in a SpeedVac and subjected to high-pH reversed-phase fractionation using an HPLC system.

#### High pH HPLC fractionation (general protocol for all LC-MS/MS/MS workflows)

High pH HPLC fractionation was performed following previously published protocols ^17^. The desalted peptide sample was reconstituted in 500 µL of Buffer A and subjected to high-pH reversed-phase fractionation using HPLC (Agilent) into a 96-well deep-well plate. Separation was performed on a Zorbax Extend-C18 column (3.5 µm, 4.6 × 250 mm) at a flow rate of 0.5 mL/min. The following gradient was used: 0-2 min: 100% Buffer C; 2-3 min: 0% to 13% Buffer D; 3-60 min: 13% to 50% Buffer D; 60-70 min: 50% to 80% Buffer D; 71-75 min: 100% Buffer D; 75-76 min: 100% to 0% Buffer D; 76-85 min: 100% Buffer C; 85-88 min: 0% to 13% Buffer D; 88-90 min: 13% to 80% Buffer D; 90-95 min: hold at 80% Buffer D; 96-101 min: 100% Buffer C; 101-104 min: 0% to 13% Buffer D; 104-106 min: 13% to 80% Buffer D; 106-111 min: hold at 80% Buffer D; 111-112 min: 80% to 0% Buffer D. The following buffers were used: buffer C - 10 mM ammonium bicarbonate (aqueous); buffer D - acetonitrile. To ensure acidification of eluting peptides, each well of the receiving plate was pre-loaded with 10 µL of 20% formic acid. Fractions were collected in a time-dependent mode across the plate in the following order: A1-A12, B1-B12, C1-C12, D1-D12, E1-E12, F1-F12, G1-G12, and H1-H12. After fractionation, samples were dried overnight using a SpeedVac vacuum concentrator. Wells from each column (e.g., A1–H1) were then combined to generate a total of 12 final fractions. These pooled samples were again dried overnight, reconstituted in 10 µL of Buffer A, and prepared for LC-MS/MS/MS analysis.

#### General protocol for liquid chromatography mass spectrometry analysis of TMT multiplexed samples

Peptide samples were analyzed by liquid chromatography-tandem mass spectrometry (LC-MS/MS/MS) using a standard 160-min gradient. The scan sequence consisted of the following steps: (1) MS1 was acquired in the Orbitrap at a resolution of 120,000 over a mass range of 400–1600 m/z, with a 40% RF lens setting. The automatic gain control (AGC) target was set to 250% (normalized), and the maximum injection time was set to automatic. Dynamic exclusion was enabled with a repeat count of 1 and an exclusion duration of 60 s. Data were acquired in profile mode. (2) The top 10 precursor ions from the MS1 scan were selected for MS2/MS3 analysis. Precursor ions were isolated using the quadrupole with a 0.7 m/z isolation window, followed by collision-induced dissociation (CID) in the ion trap. The following parameters were used: standard AGC, 35% normalized collision energy (NCE), and a maximum injection time of 120 ms. (3) Real-time search (RTS) and synchronous precursor selection (SPS) were enabled to select up to 20 MS2 fragment ions for MS3 analysis. MS3 spectra were acquired in the Orbitrap following high-energy collision-induced dissociation (HCD) with 55% NCE, a normalized AGC target of 500%, a maximum injection time of 118 ms, and a resolution of 60,000. The isolation window for MS3 was set at 0.7 m/z.

Raw data files were processed using RAW Converter (v1.1.0.22; available at github.com/proteomicsyates/RawConverter), and the resulting MS1, MS2, and MS3 files were uploaded to the Integrated Proteomics Pipeline (IP2). Peptide identification was performed using the ProLuCID algorithm against a reverse concatenated, non-redundant version of the Human UniProt database (release 2024_03). Cysteine residues were searched with a static modification for carbamidomethylation (+57.02146 Da). A differential modification for iodoacetamide desthiobiotin labeling (+398.25292 Da) was included for site-of-labeling experiments. TMT labeling was considered as a static modification on peptide N-termini and lysine residues (+229.1629 Da for TMT 10-plex, +304.2071 Da for TMT 16-plex). Search results were filtered using DTASelect (v2.0) to maintain a peptide-level false discovery rate (FDR) below 1%. Quantification was performed using MS3 reporter ions with a mass tolerance of 20 ppm via the IP2 platform.

### Data processing and analysis for TMT-exp experiments

#### Data processing

Six 10-plex experiments with primary human T cells from different biological donors were processed. Proteins were required to have at least two unique quantified peptides to pass into the final list. Peptide ratios (signal intensity/sum of signal intensities per peptide) were calculated and ratios of all peptides per protein were averaged. Keratins were not included in analysis. For experiments using the standard protocol, proteins needed to be quantified in at least 2 experiments to be included in the final table. To account for potential variation in expression between donors, proteins detected in two replicates were excluded from analysis if the ratio of the two replicates was > 2 and one replicate had an absolute fold-change < 1.5. For experiments using the low-input protocol, a curated list of proteins (PDCD1, LAG3, TIGIT, HAVCR2, CD39, NR4A1, NR4A3, GZMB, GSR, CD62L, CTLA4, PRKCH, PRKCQ, and SAMHD1) were included in the final table if they were quantified in a single experiment. For the experiment containing human TILs, the requirement that proteins be quantified in two replicates was not applied. For experiments containing *in vivo* samples, hemoglobins were excluded. For experiments using the low-input protocol, normalization factors were calculated by dividing the global median (the median of all channel medians) by each individual channel’s median. Each channel was then multiplied by its corresponding normalization factor.

#### Principal Component Analysis

Principal component analysis was carried out using *scikit-learn*^12^ library in Python. Protein ratio values were log_2_-transformed, and only proteins quantified across all experiments were included. The first two principal components (PC1 and PC2) were plotted. Loadings for plotting were selected by taking 5 proteins with highest and lowest loadings for both principal components.

#### Gene set enrichment analysis of PCA loadings

PCA loadings were used for GSEA using the prerank function in GSEAPy v.1.1.4 with the gene ontology biological process database (“c5.go.bp”) set as the database. The top 15 enriched terms based on NES were plotted.

### Proteomic platforms: Cysteine reactivity profiling, TMT-ABPP

#### Sample preparation

Five 10-plex experiments with primary human T cells from different biological donors were processed. Five experimental conditions were used in duplicate for each donor, including D2, D4A, D4C, D8A, and D8C T cells. Cells were pelleted (600 x *g*, 5 min, 4 °C), washed once with PBS, transferred into 1.5 mL Eppendorf tubes, flash-frozen, and stored at −80 °C until further processing. On the first day of the mass spectrometry workflow, frozen cell pellets were thawed on ice for 10 min. Ice-cold PBS (560 µL/sample) was added and cells were lysed by sonication (three cycles of 8 pulses at 60% output). Protein concentrations were then determined using the standard DC Protein Assay (Bio-Rad) and normalized to a final concentration of 1-2 mg/mL.

Cysteine reactivity profiling experiments with TMT multiplexing were performed following previously published protocols^17^. Briefly, normalized proteome samples (500 µL) were transferred into new 1.5 mL low-binding Eppendorf tubes and iodoacetamide-PEG-desthiobiotin was added (5 µL of 10 mM stock; 100 µM final concentration), and the samples were incubated at room temperature for 1 hour. The tubes were then placed on ice, and proteins were precipitated using a methanol/chloroform protocol described above. The resulting pellets can be stored at −80 °C overnight. The pellets were then re-suspended with sonication in 90 µL of 8M urea TEAB buffer with DTT (2.2 g urea, 6.2 mg DTT, 2 mL H_2_O, and 200 µL 1 M TEAB buffer, pH 8.5), and incubated at 65 °C for 20 min. After cooling for 5 min at room temperature, iodoacetamide was added (10 µL of 500 mM stock; 50 mM final concentration), followed by incubation at 37 °C for 30 min.

#### Trypsin/LysC digestion and streptavidin enrichment

To reduce the urea concentration, samples were diluted with 300 µL of TEAB buffer (50 mM, pH 8.5). Next, 4 µL of Trypsin/LysC (0.5 µg/µL in the appropriate buffer) and 4 µL of 100 mM CaCl₂ (final concentration 1 mM) were added, and the digestion was carried out overnight at 37 °C with shaking. The following day, 25 µL of compact, pre-washed streptavidin-agarose beads resuspended in 300 µL of enrichment buffer (50 mM TEAB, pH 8.5, with 150 mM NaCl and 0.2% NP40) were added to each sample, and the mixture was rotated for 3 h at room temperature. After the enrichment step, the beads were pelleted by centrifugation (2000 × *g*, 1 min), transferred to BioSpin columns, and washed extensively (three washes each with 1 mL of wash buffer, 1 mL of PBS, and 1 mL of water). Finally, peptides were eluted into a new Eppendorf tube with a 50% acetonitrile:water solution containing 0.1% formic acid, and the solvent was removed using a SpeedVac vacuum concentrator.

#### TMT labeling

The resulting protein digests were reconstituted in 100 µL of 30% acetonitrile in water containing 0.1% formic acid, with sonication to ensure complete dissolution, and used for TMT labeling. For each sample (“channel”), 3 µL of TMT reagent (20 µg/µL in dry acetonitrile) was added, followed by vortexing and brief centrifugation. Samples were then incubated for 1 h at room temperature. To quench unreacted TMT tags, 3 µL of 5% hydroxylamine in water was added, followed by vortexing and a 15-min incubation. The samples were acidified with 5 µL of formic acid per sample and pooled into a single Eppendorf tube. Solvent was removed overnight using a SpeedVac vacuum concentrator. The combined sample was then desalted using a SepPak C18 column (50 mg), as previously described. After another round of solvent removal via SpeedVac, the sample underwent high-pH fractionation using HPLC, following the same protocol as outlined above.

### Data processing and analysis for TMT-ABPP

Five 10-plex TMT experiments with primary human T cells from different biological donors were analyzed. The MS3-based peptide quantification was conducted using the Integrated Proteomics Pipeline (IP2), with the reporter ion mass tolerance set to 20 ppm. To ensure data quality at the level of individual TMT experiments, several filtering steps were applied: non-unique peptides, half-tryptic peptides, peptides with more than one internal missed cleavage, and peptides with low total reporter ion intensity (<10,000 across five channels per donor) were excluded. Proteins were required to have at least one unique quantified peptide in each experiment. Peptide channel ratios were calculated (peptide total intensity of each channel was normalized by the sum of signal intensities for all peptides in the same channel) and cysteine aggregation was performed to aggregate signal intensities for multiple peptides for the same cysteine (e.g. missed cleavage sites, charge states). Experiments were combined, requiring all peptides to be quantified in minimum 2 experiments. Averaged values (median) were calculated and used to calculate the R ratios of treatment groups (D4A, D8A, D4C, D8C) to control (D2).

### Reactivity change analysis

To control for potential donor-specific variability in protein expression, proteins were included in the analysis only if at least one peptide R ratio fell within a 1.5-fold range of the corresponding protein expression level observed in the TMT-exp experiments (when available). Proteins were excluded if all peptide R ratios exceeded a 2.0-fold increase or fell below a 0.5-fold decrease. Additionally, peptides detected in two replicates were excluded from analysis if the ratio of the two replicates was > 2 and one replicate had an absolute fold-change < 1.5.

For proteins with two or more quantified peptides, a cysteine was considered for potential reactivity changes if its peptide R ratio deviated by more than two-fold from the corresponding protein expression level determined by TMT-exp data. Additionally, the highest and lowest peptide R ratios within the protein had to differ by more than two-fold. All identified reactivity changes were subsequently reviewed and curated manually.

### Proteomic platforms: ATP-addback, TMT-ABPP

Two 16-plex experiments with primary human T cells from different biological donors were processed. Six experimental conditions were used in duplicate or triplicate for each donor, including D2, D2 + ATP, D8A, D8A + ATP, D8C, D8C + ATP. T cells were pelleted (600 x *g*, 5 min, 4 °C), washed once with PBS, transferred into 1.5 mL Eppendorf tubes, flash-frozen, and stored at −80 °C until further processing. On the first day of the mass spectrometry workflow, frozen cell pellets were thawed on ice for 10 min. Ice-cold PBS containing 1 mM MgCl_2_ and protease inhibitor cocktail (560 µL/sample) was added and cells were lysed by sonication (three cycles of 8 pulses at 60% output). Protein concentrations were then determined using the standard DC Protein Assay (Bio-Rad), normalized to a final concentration of 1-2 mg/mL, and transferred into new 1.5 mL low-binding Eppendorf tubes (500 µL/channel). ATP stock solution (500 mM) was prepared by dissolving powder in molecular biology grade water and adjusting pH to 7.4 with pH strips. ATP (5 µL, 500 mM) was added to ATP-addback groups, molecular biology grade water (5 µL) was added to control groups, and the mixture was incubated at ambient temperature for 10 min. After this pre-incubation step, samples were processed according to cysteine reactivity profiling workflow described above (starting with the iodoacetamide-PEG-desthiobiotin treatment step).

### Proteomic platforms: Abundance-based proteomics of detergent-soluble and insoluble proteome with TMT multiplexing

Two 12-plex experiments with primary human T cells from different biological donors were processed. Six experimental conditions were used in duplicate for each donor, including D2 soluble, D2 insoluble, D8A soluble, D8A insoluble, D8C soluble, and D8C insoluble. T cells were pelleted (600 x *g*, 5 min, 4 °C), washed once with PBS, transferred into 1.5 mL Eppendorf tubes, flash-frozen, and stored at −80 °C until further processing. On the first day of the mass spectrometry workflow, frozen cell pellets were thawed on ice, resuspended in 300 µL ice-cold fractionation buffer (20 mM Tris-HCl, pH 7.4, 150 mM NaCl, 2 mM EDTA, 1% Triton X-100, protease inhibitor cocktail), and lysed by pipetting the solution up and down. Protein concentrations were then determined using the standard DC Protein Assay (Bio-Rad) and normalized to a final concentration of 3 mg/mL. Normalized cell lysates were transferred to new low binding Eppendorf tubes (400 µL/tube) and centrifuged at 17,000 x *g* for 10 min at 4 °C. The supernatant was saved as a soluble fraction and the pellet was washed twice with ice-cold fractionation buffer. The insoluble pellet was reconstituted in 400 µL ice-cold fractionation buffer and lysed by sonication (three cycles of eight 1 second pulses at 60% output). The detergent-soluble and insoluble fractions (300 µL/channel) were processed according to abundance-based proteomics workflow described above (starting with the addition of urea and DTT for reduction).

### Proteomic platforms: Low-input abundance-based proteomics with TMT multiplexing

Cells were pelleted (600 x *g*, 5 min, 4 °C), washed once with PBS, transferred into 1.5 mL Eppendorf tubes, flash-frozen, and stored at −80 °C until further processing. Low-input abundance-based proteomics with TMT multiplexing was performed following a previously published protocol with slight modifications^18, 19^.

#### Trypsin/LysC digestion

On the first day of the mass spectrometry workflow, frozen cell pellets were thawed on ice for 5 min. Ice-cold PBS supplemented with cOmplete EDTA-free Protease Inhibitor Cocktail (1 tablet per 10 mL) was added, and cells were lysed by sonication (three cycles of 8 pulses at 60% output). Protein concentrations were then determined using the standard DC Protein Assay (Bio-Rad) and normalized to a final concentration of 0.5 mg/mL. Normalized proteome samples (20 µL) were transferred into PCR tubes. For each sample (“channel”), 3 µL of SP3 beads (1:1 mixture of hydrophobic and hydrophilic type in lysis buffer) and 30 µL of ethanol containing 20 mM DTT were added, followed by pipetting and brief centrifugation. After a 15 min incubation at room temperature, the PCR tubes were placed on a magnetic stand, and the liquid was aspirated. The beads were washed once with 100 µL of 80% ethanol, resuspended in 25 µL of PBS containing 20 mM iodoacetamide, incubated in the dark for 30 min at room temperature, and then 50 µL of ethanol containing 20 mM DTT was added. After another 15 min incubation at room temperature, the beads were washed twice with 80% ethanol. The remaining beads were resuspended in 30 µL of 200 mM EPPS buffer (pH 8) containing 0.3 µg of Lys-C and 0.3 µg of trypsin and incubated at 37 °C overnight.

#### TMT labeling

The next day, 9 µL of acetonitrile and appropriate TMT tag (3 µL per sample; 20 µg/µL in anhydrous acetonitrile) were added to the mixture of digested peptides and beads, followed by gentle mixing and brief centrifugation. After a 60 min incubation at room temperature, unreacted tags were quenched by adding 7 µL of 5% hydroxylamine in water. All TMT-labeled samples were combined into a single Eppendorf tube, dried using a SpeedVac, and desalted using a 100-mg Sep-Pak column. After another round of solvent removal via SpeedVac, the sample underwent high-pH fractionation using HPLC, following the same protocol as outlined above.

### Proteomic platforms: Low-input cysteine reactivity profiling, TMT-ABPP

Cells were pelleted (600 x *g*, 5 min, 4 °C), washed once with PBS, transferred into 1.5 mL Eppendorf tubes, flash-frozen, and stored at −80 °C until further processing. Low-input cysteine reactivity profiling with TMT multiplexing was performed following a previously published protocol with slight modifications^19^.

#### IA-PEG-DTB labeling and Trypsin/LysC digestion

On the first day of the mass spectrometry workflow, frozen cell pellets were thawed on ice for 5 min. Ice-cold PBS supplemented with cOmplete EDTA-free Protease Inhibitor Cocktail (1 tablet per 10 mL) was added, and cells were lysed by sonication (three cycles of 8 pulses at 60% output). Protein concentrations were then determined using the standard DC Protein Assay (Bio-Rad) and normalized to a final concentration of 1-2 mg/mL. Normalized proteome samples (15 µL) were transferred into PCR tubes. For each sample (“channel”), iodoacetamide-PEG-desthiobiotin was added (5 µL of 2 mM stock; 500 µM final concentration), and the samples were incubated at room temperature for 1 h. After that, 3 µL of SP3 beads (1:1 mixture of hydrophobic and hydrophilic type in lysis buffer) and 30 µL of ethanol containing 20 mM DTT were added, followed by pipetting and brief centrifugation. After a 15 min incubation at room temperature, the PCR tubes were placed on a magnetic stand, and the liquid was aspirated. The beads were washed once with 100 µL of 80% ethanol, resuspended in 25 µL of PBS containing 20 mM iodoacetamide, incubated in the dark for 30 min at room temperature, and then 50 µL of ethanol containing 20 mM DTT was added. After another 15 min incubation, the beads were washed twice with 80% ethanol. The remaining beads were resuspended in 30 µL of 200 mM EPPS buffer (pH 8) containing 0.3 µg of Lys-C and 0.3 µg of trypsin and incubated at 37 °C overnight.

#### TMT labeling

The next day, 9 µL of acetonitrile and appropriate TMT tag (3 µL per sample; 20 µg/µL in anhydrous acetonitrile) were added to the mixture of digested peptides and beads, followed by gentle mixing and brief centrifugation. After a 60 min incubation at room temperature, unreacted tags were quenched by adding 7 µL of 5% hydroxylamine in water. All TMT-labeled samples were combined into a single Eppendorf tube, dried using a SpeedVac, and desalted using a 100-mg Sep-Pak column.

#### Streptavidin enrichment

The desalted TMT-labeled peptides were resuspended in 460 µL of 100 mM HEPES buffer (pH 7.4), 80 µL Pierce High Capacity Streptavidin Agarose (Thermo Fisher Scientific) were added and the mixture was incubated at room temperature for 3 h. The resulting mixture was then loaded on a Ultrafree-MC centrifugal filter (hydrophilic PTFE, 0.22 µm pore size) and centrifugated at 1,000 x *g* for 30 s. Beads were washed sequentially with 300 µL of 100 mM HEPES (pH 7.4) with 0.05% NP-40 twice, 350 µL of 100 mM HEPES (pH 7.4) three times, and 400 µL of H_2_O once. Peptides were eluted sequentially by 1) elution buffer (80% acetonitrile, 0.1% formic acid) with 20 min incubation at room temperature, 2) elution buffer with 20 min incubation at room temperature; 3) elution buffer with 10 min incubation at 72 °C. The combined eluent was dried in a SpeedVac.

#### High pH fractionation

The desalted peptide sample was reconstituted in 300 µL of 0.1% TFA solution and subjected to high-pH reversed-phase fractionation using a fractionation kit (Thermo Fisher Scientific) into Eppendorf tubes. The sample was fractionated into 8 fractions, dried using a SpeedVac, reconstituted in 10 µL of 5% formic acid/5% acetonitrile solution, and prepared for LC-MS/MS/MS analysis.

### Western blot analysis

T cells were pelleted (600 x *g*, 5 min, 4 °C), washed with PBS, transferred to 1.5 mL Eppendorf tubes, flash-frozen, and stored at −80 °C until further analysis. On the day of the analysis, frozen cell pellets were thawed on ice, resuspended in ice-cold PBS containing protease inhibitor cocktail (Roche) and phosphatase inhibitor cocktail (Roche) and lysed by sonication (3 x 8 pulses, 60% duty cycle). Protein concentrations were adjusted to 1-2 mg/mL, 4x Laemmli sample buffer was added, and the samples were heated at 95 °C for 7 min. Proteins were resolved using SDS-PAGE (4-20% Criterion TGX Stain-Free Protein Gel, BioRad) and transferred to 0.45 μm polyvinylidene fluoride membranes (Cytiva). For total protein staining, the membrane was fully dried at room temperature for 1 h and rehydrated using MeOH. Following 30 s incubation, the membrane was washed with PBS and H_2_O, and then stained with Revert 700 (LICORbio) at room temperature for 5 min. The membrane was rinsed 2 times with Revert 700 wash solution (LICORbio) and the images were captured using the 680 nm channel on a ChemiDoc MP Imaging System (Bio-Rad). After capturing the images of total protein staining, the membranes were blocked with 5% milk or 5% BSA in Tris-buffered saline (20 mM Tris-HCl pH 7.6, 150 mM NaCl) with 0.1% tween 20 (TBS-T) buffer at room temperature for 30 min, washed 3 times with TBS-T, and incubated with primary antibodies in 5% BSA in TBS-T at 4 °C overnight. The membranes were washed 3 times with TBS-T, incubated with HRP-conjugated secondary antibodies in 5% BSA in TBS-T at room temperature for 1 h and washed 5 times with TBS-T. The immunoreactive bands were detected using Pierce ECL Western Blotting Substrate (Thermo Fisher Scientific) on the ChemiDoc MP Imaging System (Bio-Rad). Blots were quantified with densitometric analysis using ImageJ software (NIH) and normalized to the total amount of protein in each lane.

### Subcellular fractionation

Activated (D2), acutely (D8A), and chronically stimulated (D8C) T cells were pelleted (600 x *g*, 5 min, 4 °C), washed with PBS, transferred to low binding 1.5 mL Eppendorf tubes, flash-frozen, and stored at −80 °C until further analysis. On the day of the analysis, frozen cell pellets were thawed on ice, resuspended in ice-cold fractionation buffer (20 mM HEPES [pH 7.4], 250 mM sucrose, 10 mM KCl, 2 mM MgCl_2_, 1 mM EGTA, 1 mM EDTA, 1 mM DTT, protease inhibitor cocktail), homogenized by passing cell suspension through a 25 gauge needle 20 times and incubated on ice for 20 min. All samples were kept at 4 °C for the duration of the fractionation. The nuclei were pelleted by centrifugation (720 x *g*, 5 min, 4 °C) and the supernatant, which includes the mitochondria, membrane and cytoplasm, was saved for further processing. The nuclear pellet was washed and resuspended in 500 µL of fractionation buffer, passed through a 25 gauge needle 20 times, and pelleted (720 x *g*, 10 min, 4 °C). The wash step was repeated one more time. The nuclear pellet was then resuspended in TBS with 0.1% SDS with sonication (2 x 8 pulses, 60% duty cycle) to shear genomic DNA and homogenize the suspension. The supernatant containing the mitochondria, membrane and cytoplasm was centrifuged (10,000 x *g*, 5 min, 4 °C) to pellet the mitochondrial fraction. The supernatant was transferred to a new Eppendorf tube. The mitochondrial pellet was then washed 3 times in fractionation buffer and resuspended in 200 µL of TBS with 0.1% SDS with sonication (2 x 8 pulses, 60% duty cycle). The supernatant containing the membrane and cytoplasm was ultracentrifuged (100,000 x *g*, 1 h, 4 °C). The supernatant was re-centrifuged (100,000 x *g*, 45 min, 4 °C) and saved as cytoplasm fraction. The membrane pellet was washed, resuspended in 400 µL of fractionation buffer, passed through a 25 gauge needle, and re-centrifuged (100,000 x *g*, 45 min, 4 °C). The membrane pellet was then resuspended in 500 µL of TBS with 0.1% SDS with sonication (2 x 8 pulses, 60% duty cycle). Protein concentration was adjusted to 2.0 mg/mL and the samples were mixed with 4x Laemmli sample buffer, boiled at 95 °C for 7 min, resolved by SDS-PAGE and immunoblotted.

### Fractionation of Triton X-100 soluble and insoluble proteins

T cells were pelleted (600 x *g*, 5 min, 4 °C), washed with PBS, transferred to low binding 1.5 mL Eppendorf tubes, flash-frozen, and stored at −80 °C until further analysis. On the day of the analysis, cell pellets were thawed on ice, resuspended in 150 µL ice-cold fractionation buffer (20 mM Tris-HCl, pH 7.4, 150 mM NaCl, 2 mM EDTA, 1% Triton X-100, protease inhibitor cocktail), and lysed by pipetting the solution up and down. Protein concentrations were measured using a standard DC assay (Bio-Rad) and normalized to 1-2 mg/mL. Normalized cell lysates were transferred to new low binding Eppendorf tubes (300 µL/tube) and centrifuged at 17,000 x *g* for 10 min at 4 °C to separate into supernatant and pellet fractions. The supernatant was saved as soluble fraction and the pellet was washed with ice-cold fractionation buffer. The supernatant and insoluble pellet were reconstituted in equivalent volumes of Laemmli sample buffer, boiled at 95 °C for 7 min and analyzed by Western blotting.

### Western blot based cysteine reactivity profiling

Frozen cell pellets were thawed on ice for 5 minutes. Cells were resuspended in PBS containing 1 mM MgCl_2_ and cOmplete EDTA-free Protease Inhibitor Cocktail (1 tablet per 10 mL), and lysed by sonication (three cycles of 8 pulses at 60% output). Protein concentrations were then determined using the standard DC Protein Assay (Bio-Rad), normalized to a final concentration of 1 mg/mL, and transferred into low binding 1.5 mL Eppendorf tubes (46 µL/condition). ATP stock solution (125 mM) was prepared by dissolving powder in molecular biology grade water and adjusting pH to 7.4 with pH strips. The stock was then diluted to 62.5, 25, 12.5, 6.25, and 2.5 mM with molecular biology grade water for concentration-dependency experiments. ATP stock solutions (2 µL) were then added to ATP-addback treatment groups, molecular biology grade water (1 µL) was added to control groups, and the mixture was incubated at room temperature for 10 min. After the pre-incubation step, a cysteine reactive probe MM(PEG)_24_ (2 µL of 250 mM stock; 10 mM final concentration) was added to samples, and the mixture was incubated at room temperature for 30 min. The reaction was quenched with 16.6 μL 4x sample buffer and boiled at 95 °C for 7 min. Samples were then analyzed by Western blot. After band intensity quantification with ImageJ software (NIH), EC_50_ curves were generated in GraphPad Prism using nonlinear fit ([Inhibitor] vs. response: Variable slope, four parameters) settings.

### Proteasome activity assay

Frozen cell pellets were thawed on ice for 5 min. Cells were resuspended in PBS containing 1 mM MgCl_2_ and cOmplete EDTA-free Protease Inhibitor Cocktail (1 tablet per 10 mL), and lysed by sonication (three cycles of 8 pulses at 60% output). Protein concentrations were then determined using the standard DC Protein Assay (Bio-Rad), normalized to a final concentration of 1.14 mg/mL, and transferred into PCR tubes (22 µL/condition). MG132 (1 µL of 625 µM stock; 25 µM final concentration) was added to MG132 groups, DMSO (1 µL) was added to control groups, and the mixture was incubated at room temperature for 1 h. ATP stock solutions (2.5, 25, 125 mM) were prepared by dissolving powder in molecular biology grade water and adjusting pH to 7.4 with pH strips. Following the MG132 treatment, ATP (1 µL; 0.1, 1, 5 mM final concentrations) was added to ATP-addback groups, molecular biology grade water (1 µL) was added to control groups, and the mixture was incubated at room temperature for 10 min. After these pre-incubation steps, a pan-reactive fluorescent proteasome activity-based probe Me4BodipyFL-Ahx3Leu3VS (1 µL of 25 µM stock; 1 µM final concentration) was added to samples, and the mixture was incubated at room temperature for 1 h. The reaction was quenched with 8.3 μL 4x sample buffer and boiled at 95 °C for 7 min. Samples were then resolved on SDS-PAGE (4-20% Criterion TGX Stain-Free Protein Gel, BioRad) and analyzed using the appropriate fluorescence channel on a ChemiDoc MP Imaging System (Bio-Rad).

### Inhibitor treatments

BSJ-04-122 (Millipore Sigma) was used to inhibit MAP2K4/7^20^ and was diluted from a 10 mM stock in DMSO to a final concentration of 10 µM. ISRIB (MedChemExpress) was used to inhibit integrated stress response^21^ and was diluted from a 200 µM stock in DMSO to a final concentration of 200 nM. Thapsigargin (Millipore Sigma) was used to induce ER stress^22^ and was diluted from a 25 µM stock in DMSO to a final concentration of 25 nM. T cells were cultured with BSJ-04-122 or ISRIB from day 2 to day 8, or Thapsigargin from day 7 to day 8.

## Statistical Analysis

Statistical analysis of flow cytometry, Western blot, and functional assays was performed using GraphPad Prism 10.3.0. Samples derived from a given blood donor were treated as paired for comparisons by one- and two-way repeated measures ANOVA. Where relevant, Gaussian distribution and sphericity were assumed, and multiple comparisons corrections included. Details of statistical tests are available in figure legends.

## Generation of reference protein tables

Lists of reference proteins were created by retrieving data from the Gene Ontology (go-basic.obo, downloaded on June 20, 2023). Protein lists were created by identifying relevant Gene Ontology terms based on text searches:

| Reference list name | GO terms used | GO term names |
| --- | --- | --- |
| Nucleotide metabolism | GO:0008252, GO:0055086 | nucleotidase activity, nucleobase-containing small molecule metabolic process |
| Cell cycle | GO:0007049, GO:0051301 | cell cycle, cell division |
| Redox-related | GO:0045454, GO:0061692, GO:0098869, GO:0071500, GO:0034599, GO:0034614, GO:0071461, GO:0042744, GO:0042743, GO:0072593, GO:0006979, GO:0000302, GO:0051775 | cell redox homeostasis, cellular detoxification of hydrogen peroxide, cellular oxidant detoxification, cellular response to nitrosative stress, cellular response to oxidative stress, cellular response to reactive oxygen species, cellular response to reactive oxygen species, cellular response to redox state, hydrogen peroxide catabolic process, hydrogen |
|  |  | peroxide metabolic process, reactive oxygen species metabolic process, response to oxidative stress, response to reactive oxygen species, response to reactive oxygen species, response to redox state |
| Mitochondrial proteins | GO:0005739, GO:0031966, GO:0005759, GO:0098800, GO:0005758, GO:0044233, GO:0005740, GO:0042645, GO:0098799, GO:0005746 | mitochondrion, mitochondrial membrane, mitochondrial matrix, inner mitochondrial membrane protein complex, mitochondrial intermembrane space, mitochondria-associated endoplasmic reticulum membrane, mitochondrial envelope, mitochondrial nucleoid, outer mitochondrial membrane protein complex, mitochondrial respirasome |
| Peroxisomal proteins | GO:0005777, GO:0005778, GO:0005782, GO:1990429 | peroxisome, peroxisomal membrane, peroxisomal matrix, peroxisomal importomer complex |
| Nucleotide binding proteins | GO:0000166, GO:0005525, GO:0002135, GO:0005524 | nucleotide binding, GTP binding, CTP binding, ATP binding |
| Respiratory complex I-IV | GO:0005747, GO:0005749, GO:0005750, GO:0005751 | mitochondrial respiratory chain complex I, mitochondrial respiratory chain complex II, succinate dehydrogenase complex (ubiquinone), mitochondrial respiratory chain complex III, mitochondrial respiratory chain complex IV |
| Respiratory complex V | GO:0045259 | proton-transporting ATP synthase complex |

## Generation of reference metabolite tables

Lists of reference metabolites were created by retrieval of reference metabolism pathways from the Kyoto Encyclopedia of Genes and Genomes (KEGG) followed by manual curation:

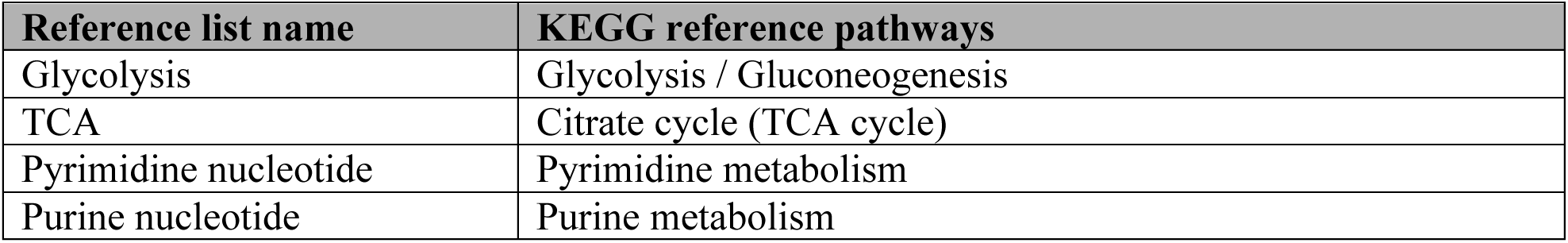

## (C) Key resources

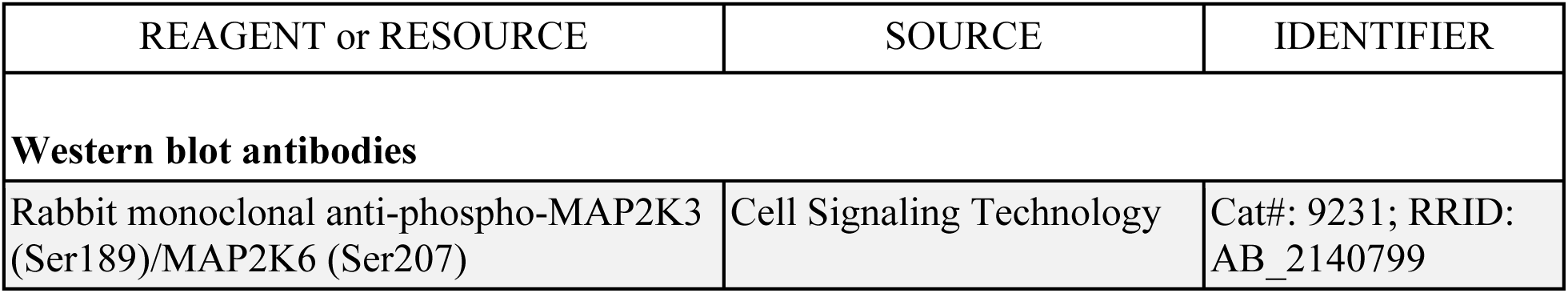
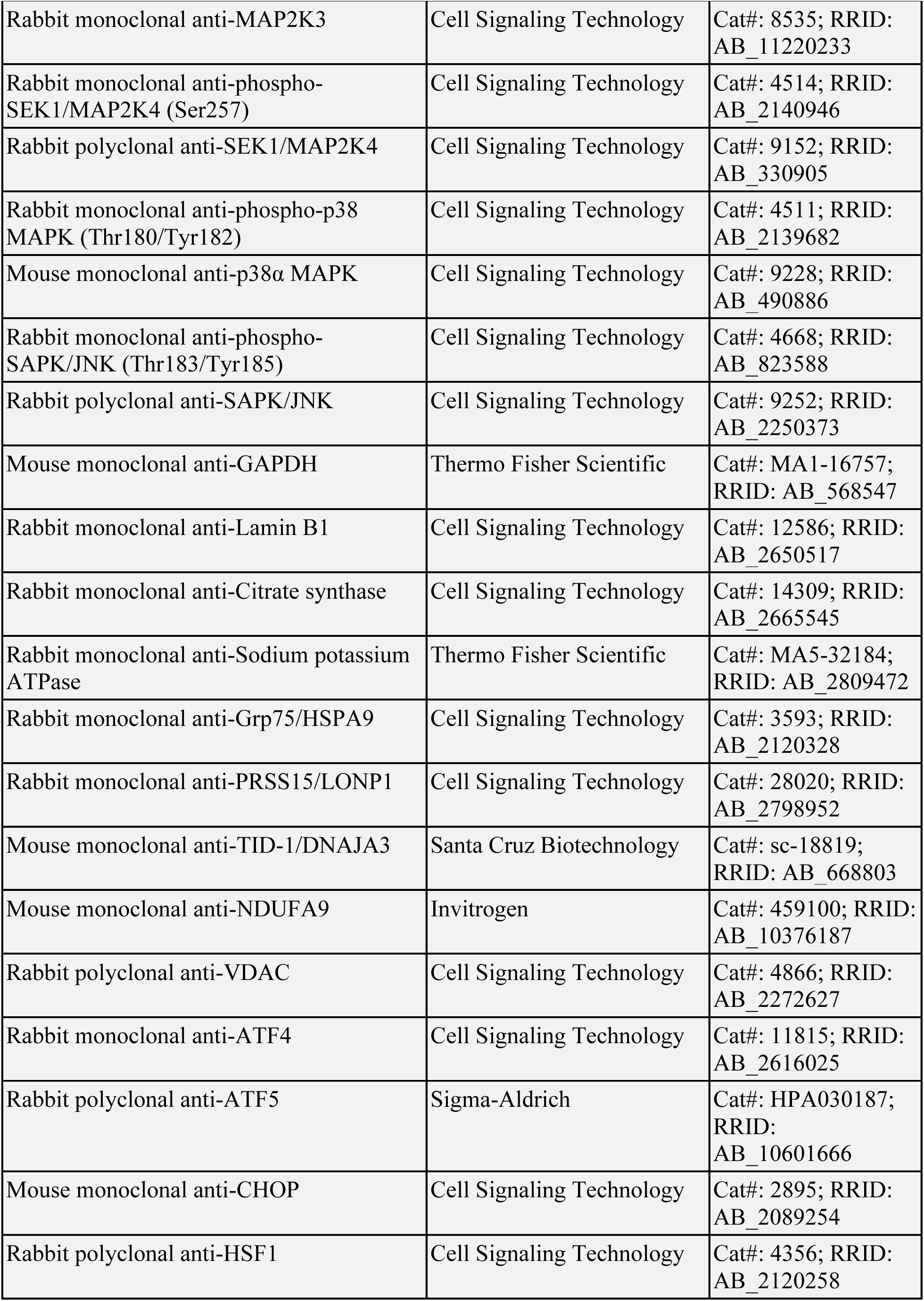
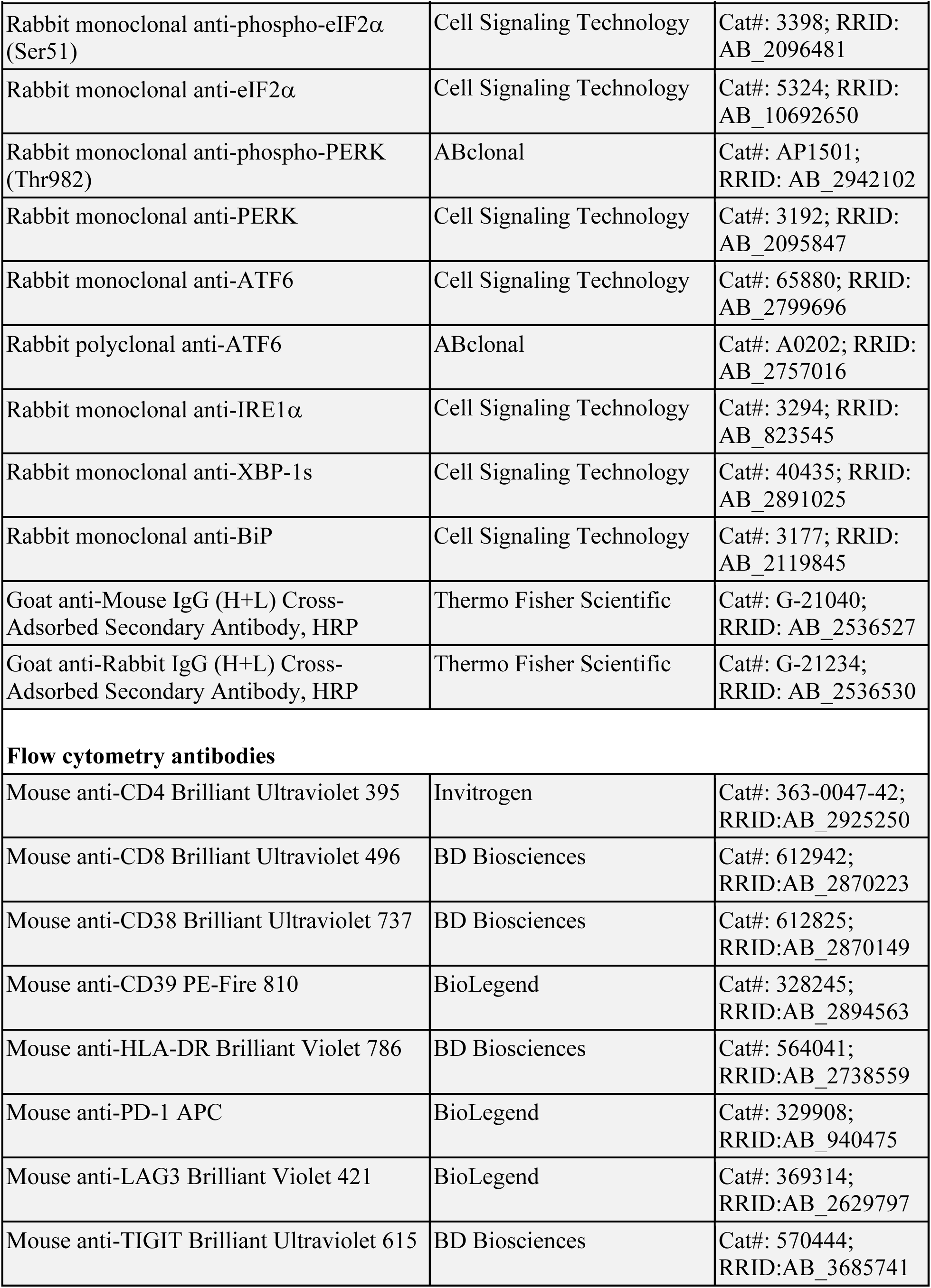
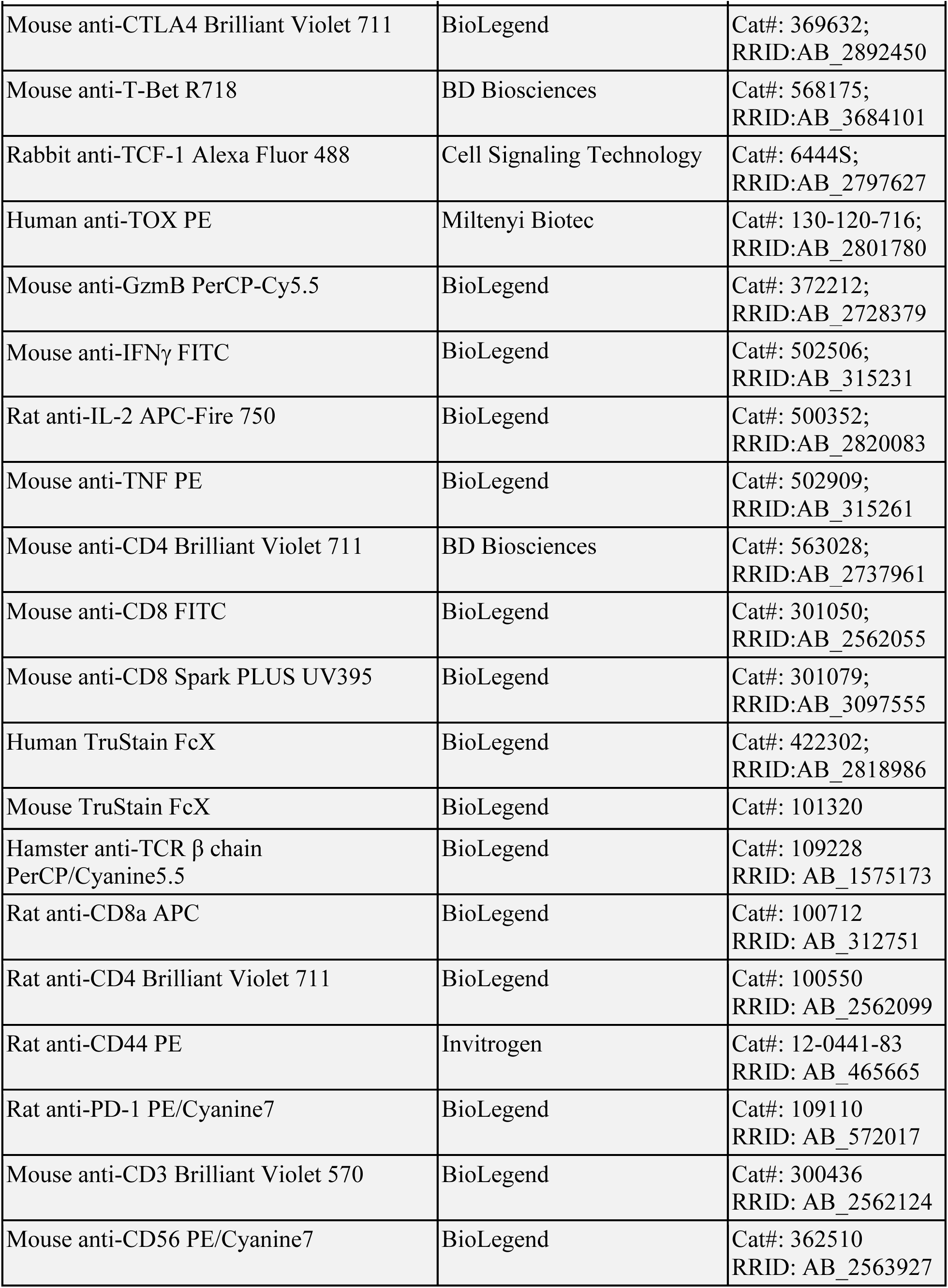
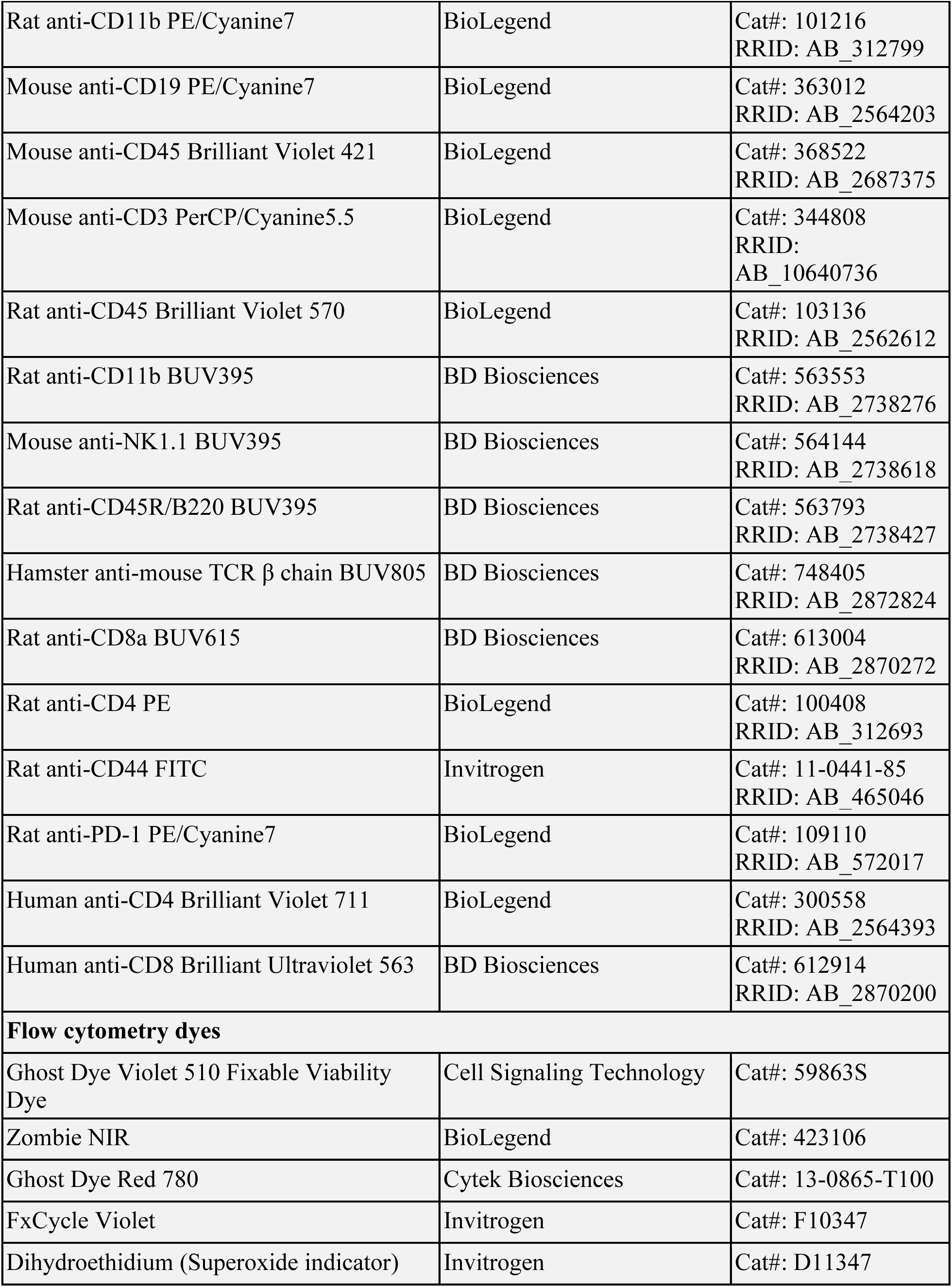
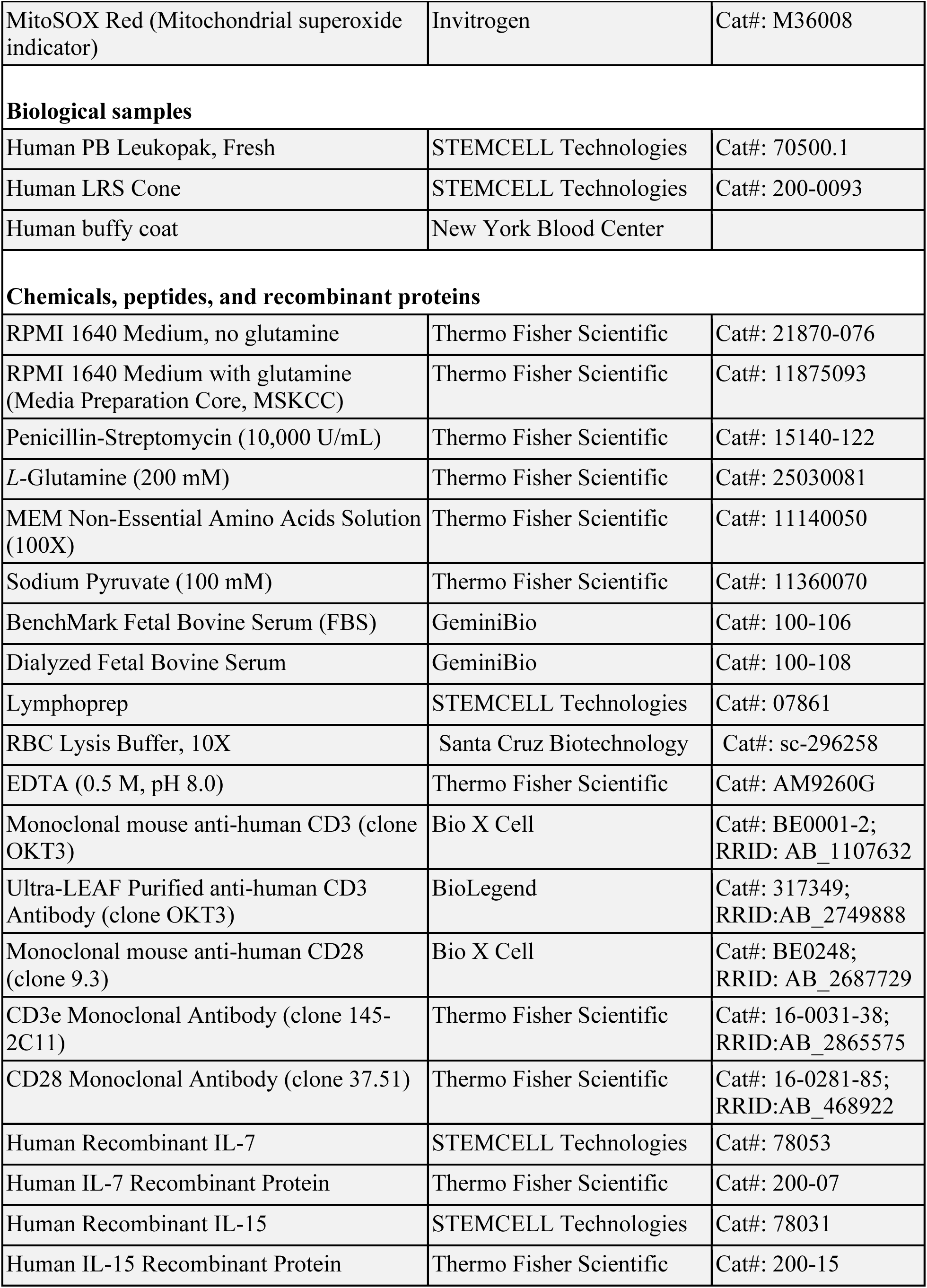
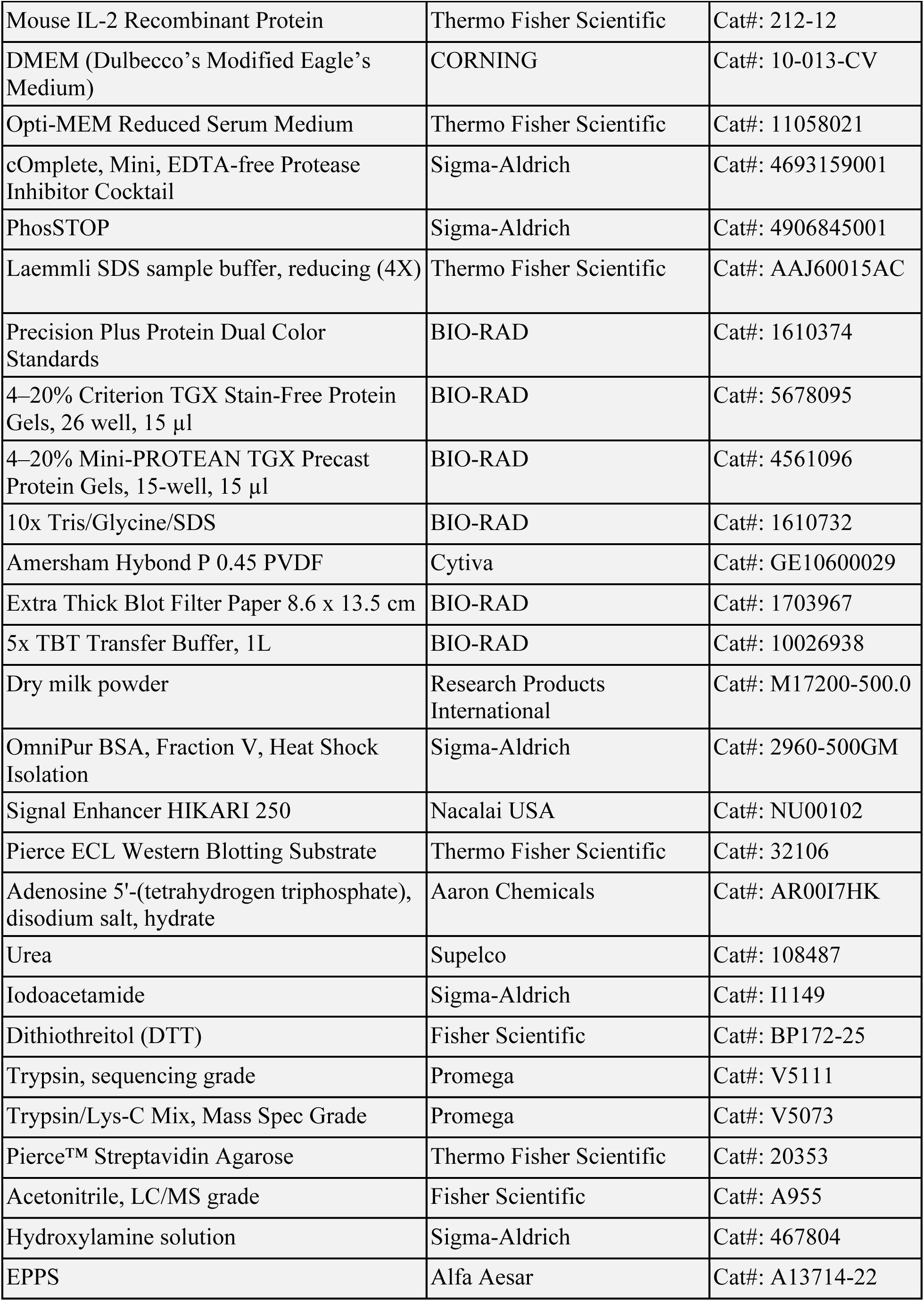
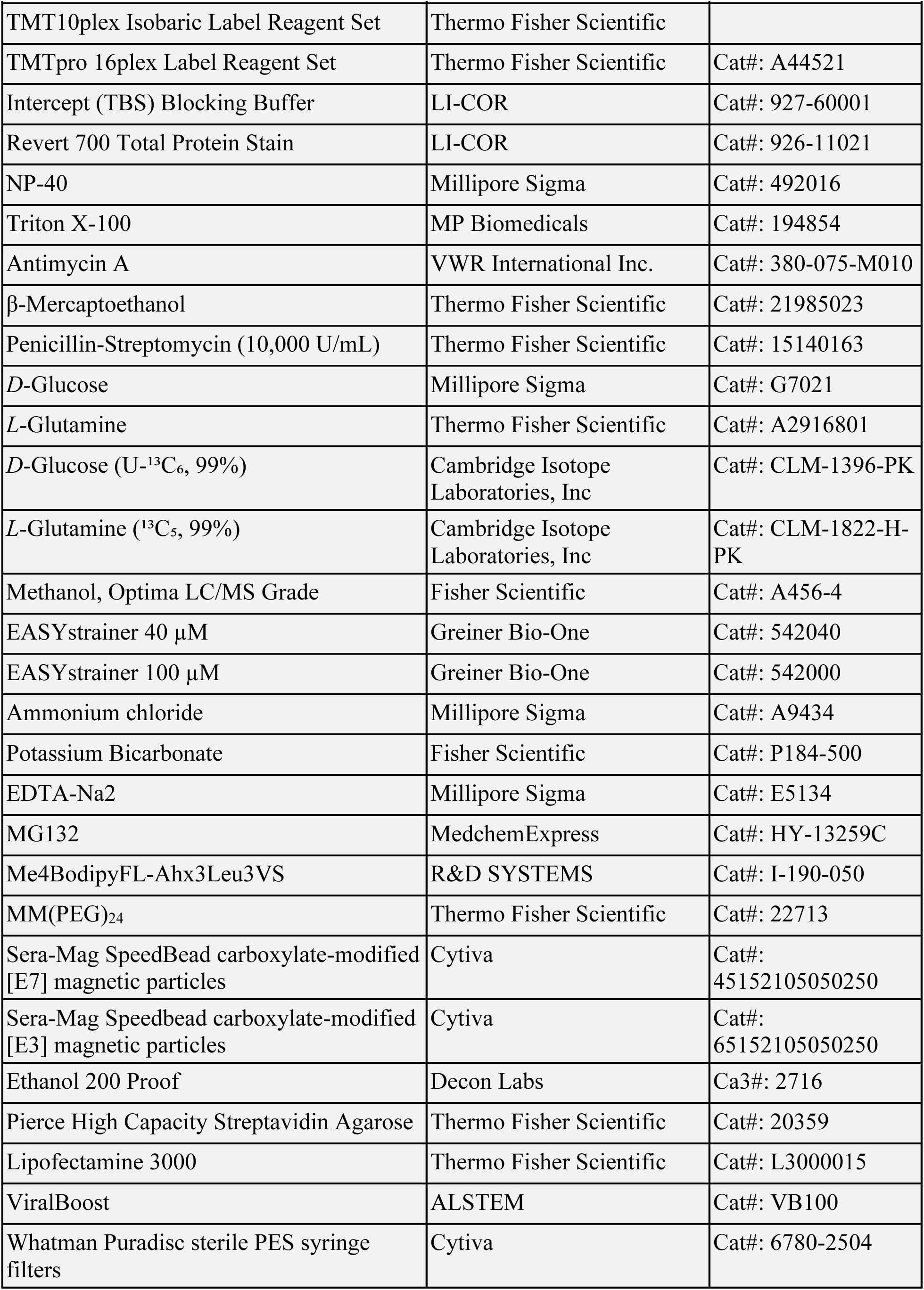
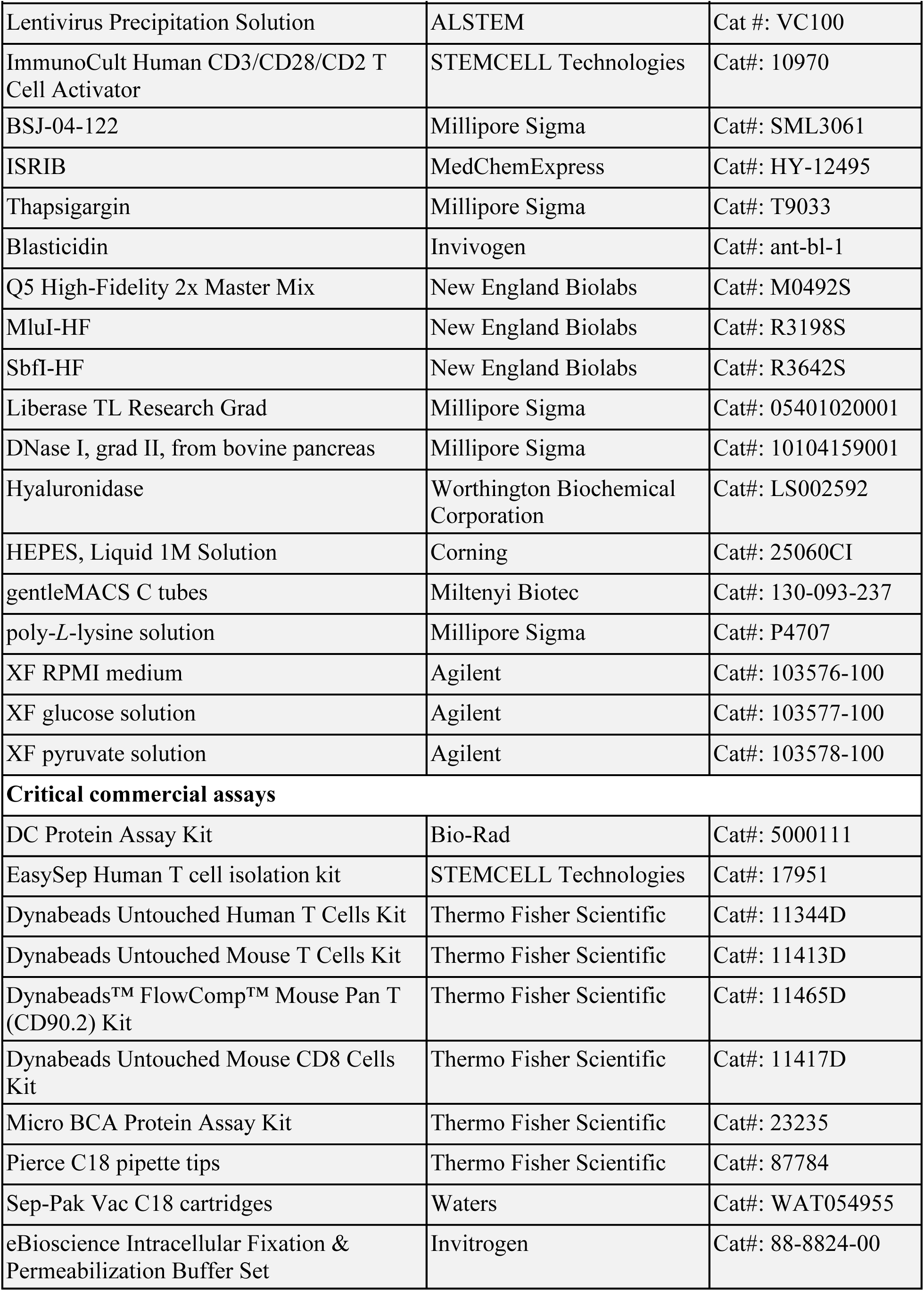
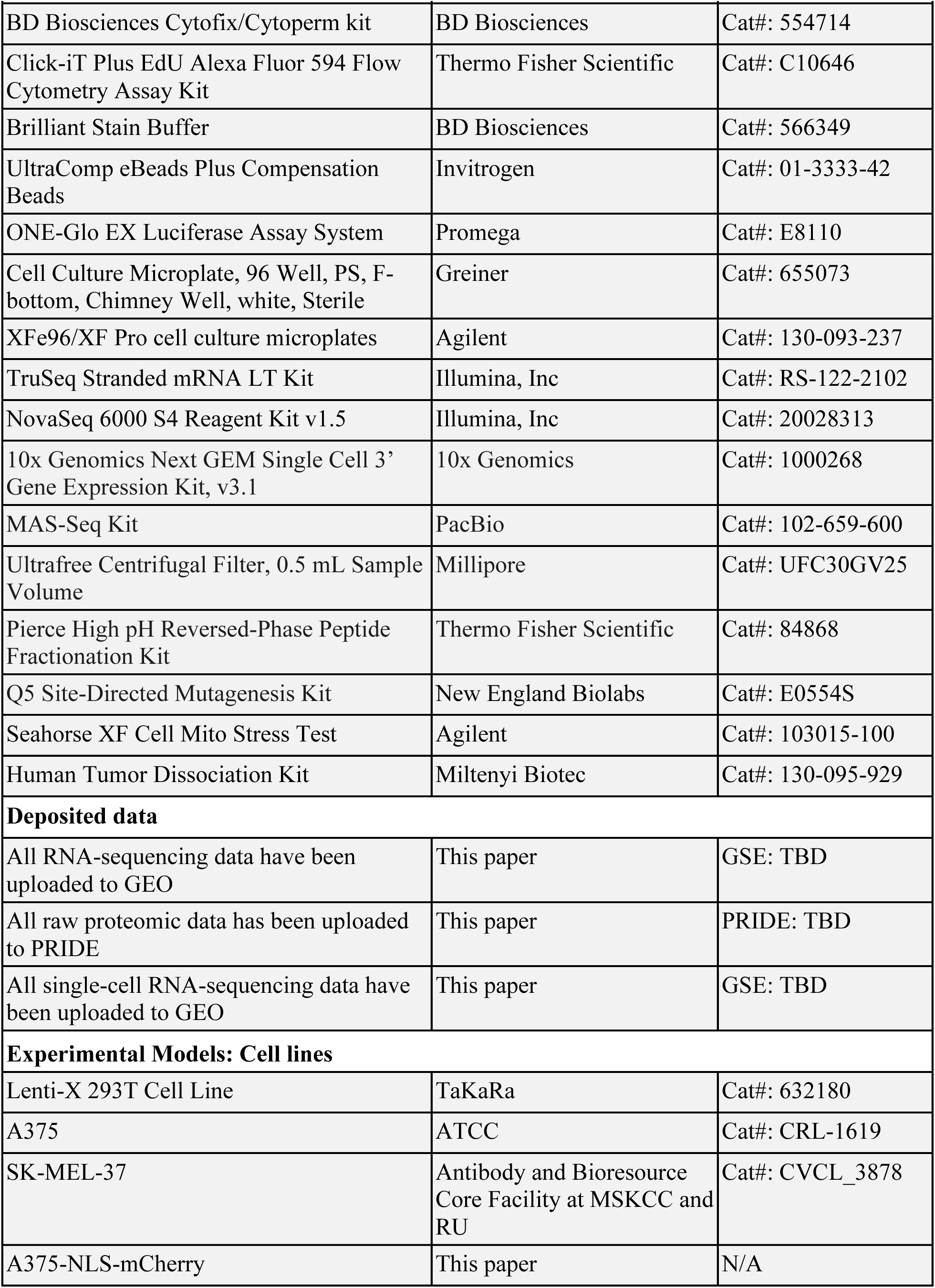
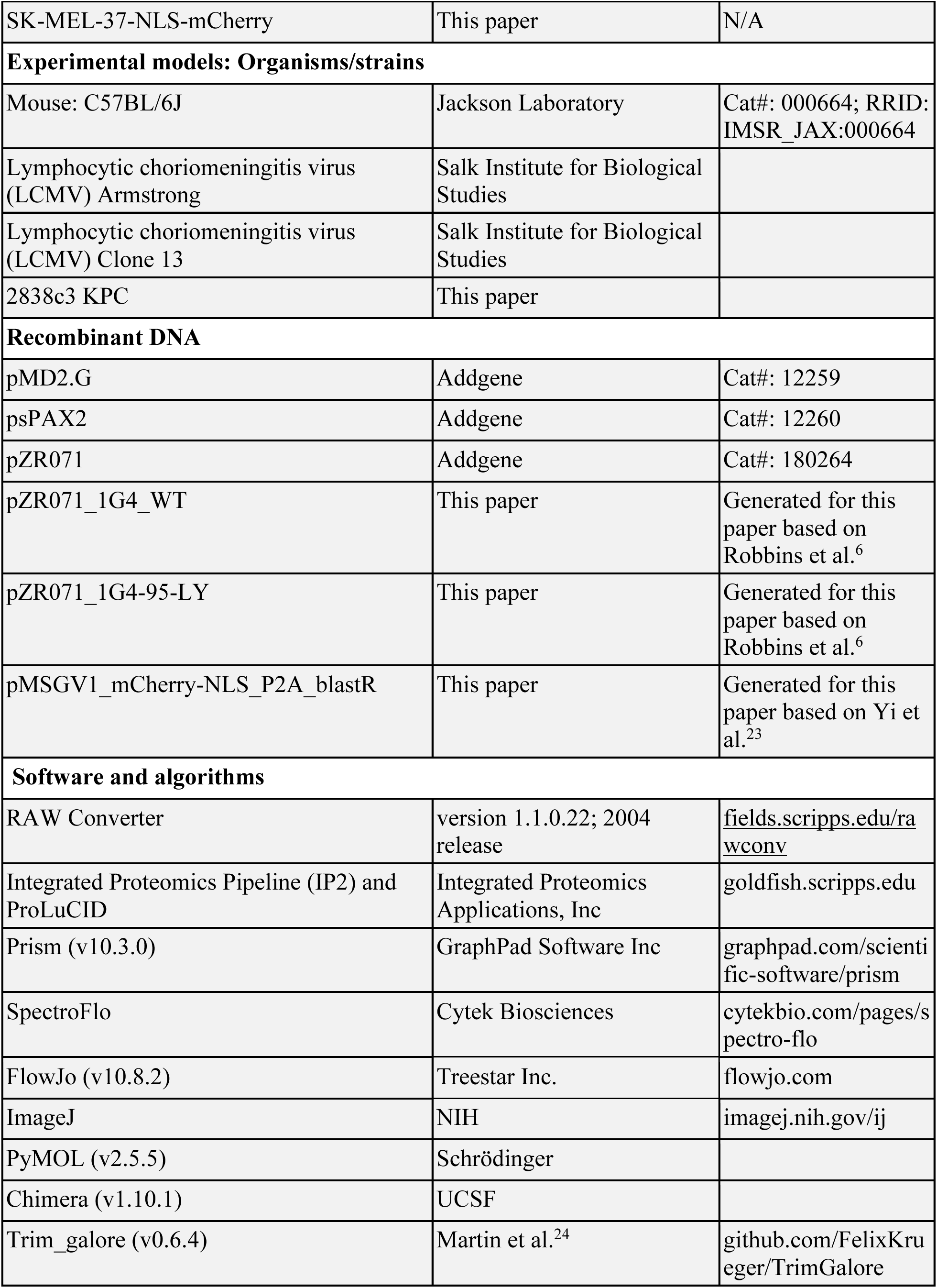
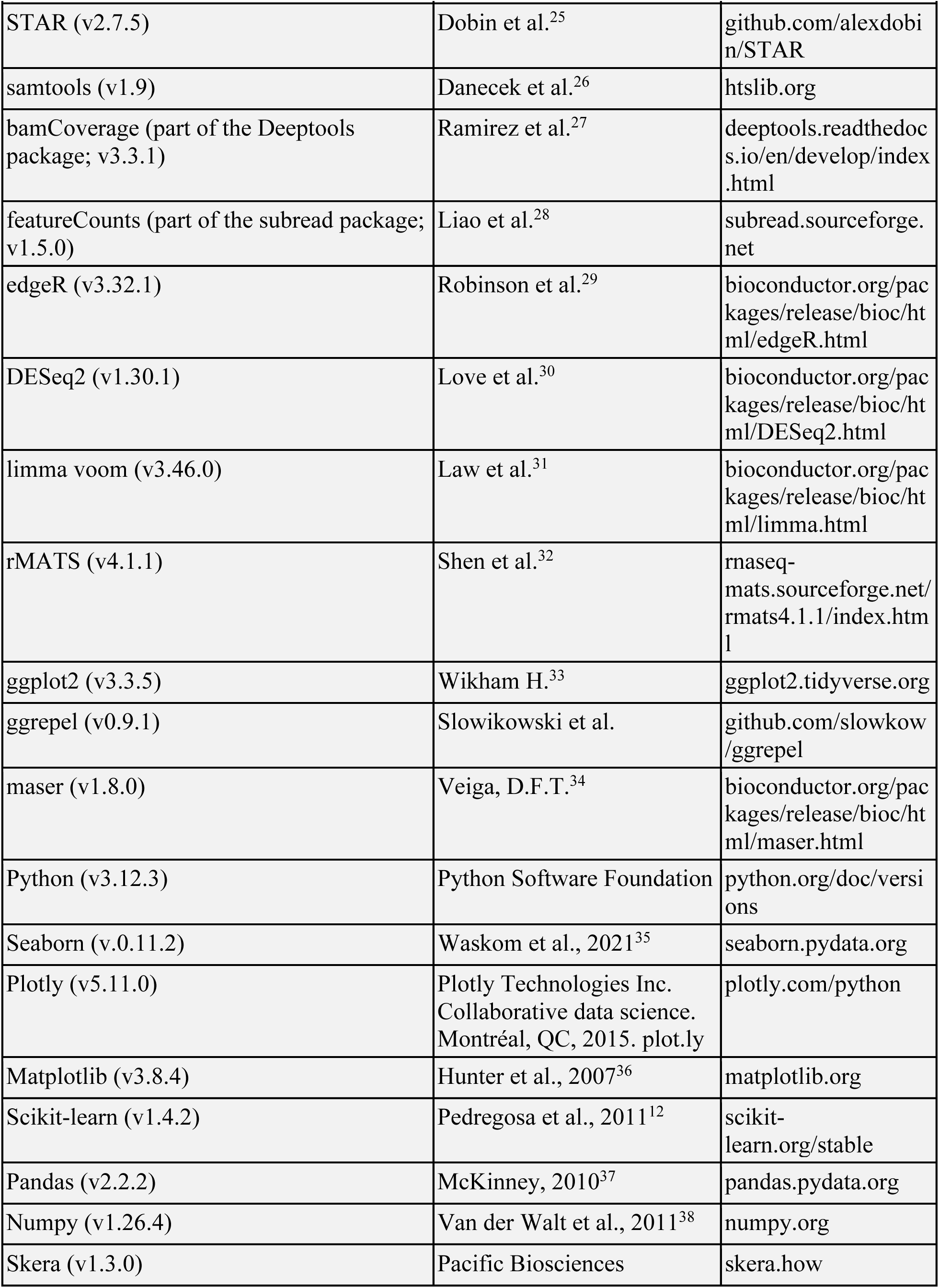
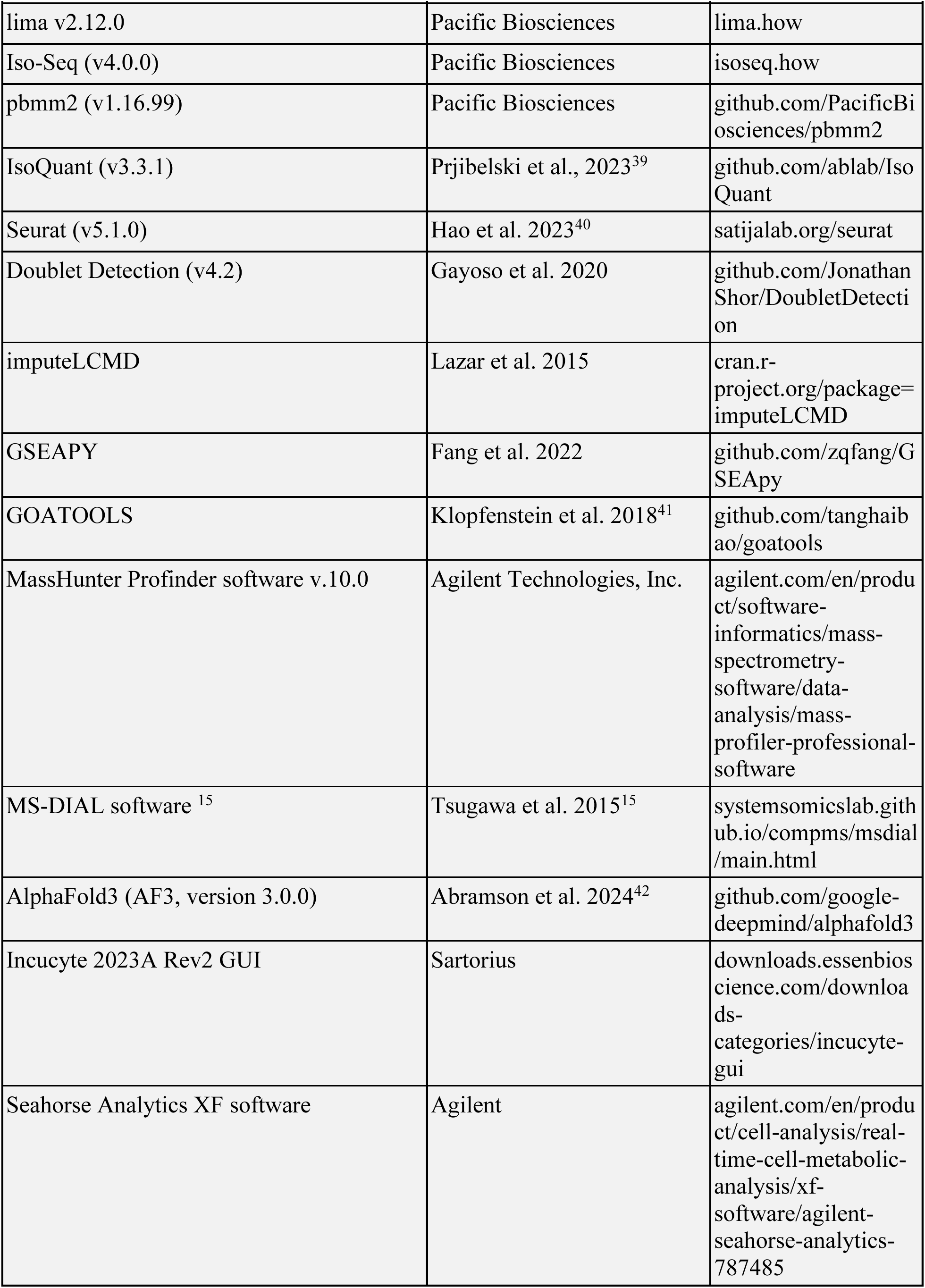

## Notes

### Summary of Updates

The manuscript has been revised to streamline the narrative.

